# Formation of biomolecular condensates in bacteria by tuning protein electrostatics

**DOI:** 10.1101/2020.05.02.072645

**Authors:** Vivian Yeong, Emily G. Werth, Lewis M. Brown, Allie C. Obermeyer

## Abstract

Biomolecular condensates provide a strategy for cellular organization without a physical membrane barrier while allowing for dynamic, responsive organization of the cell. To date, very few biomolecular condensates have been identified in prokaryotes, presenting an obstacle to engineering these compartments in bacteria. As a novel strategy for bacterial compartmentalization, protein supercharging and complex coacervation were employed to engineer liquid-like condensates in *E. coli*. A simple model for the phase separation of supercharged proteins was developed and used to predict intracellular condensate formation. Herein, we demonstrate that GFP-dense condensates formed by expressing GFP variants of sufficient charge in cells are dynamic and enrich specific nucleic acid and protein components. This study provides a fundamental characterization of intracellular phase separation in *E. coli* driven by protein supercharging and highlights future utility in designing functional synthetic membraneless organelles.

## Introduction

Cellular sub-compartmentalization has provided a method for spatially sequestering biomolecules from their surroundings, permitting the co-existence of separate, distinct environments within the cytoplasm and allowing reactions to occur that would otherwise be thermodynamically unfavorable (*1*). Manipulating the spatial localization of pathway components has been shown to improve product flux, protect against the buildup of toxic intermediates, and prevent flux through alternative metabolic pathways by sequestering intermediates (*1-3*). Consequently, spatial separation confers a variety of benefits for industrial and metabolic engineering applications.

Despite the presence of significantly fewer subcellular compartments in prokaryotes, a few endogenous compartments have been comprehensively studied. One example of bacterial compartments includes magnetosomes, which orient magnetotactic bacteria along an external magnetic field, and are decorated with lipid bilayer membranes that encapsulate ∼50 nm magnetite or greigite crystals (*4*). In contrast to lipid delimited compartments, another class of naturally existing bacterial microcompartments (BMCs) are bounded by a hollow protein shell (*1*). Examples of endogenous BMCs include carboxysomes, which house rubisco required for carbon dioxide fixation; PDU microcompartments, which facilitate 1,2-propanediol metabolism; and Eut BMCs in *E. coli*, which localize enzymes for ethanolamine utilization and sequester volatile pathway intermediates (*5*). While the majority of known bacterial compartments are delineated by a physical boundary, recent studies have reported compartments formed through intracellular phase separation in *Caulobacter crescentus* (*6, 7*). Similar to their eukaryotic counterparts, these bacterial condensates are thought to form through liquid-liquid phase separation (LLPS) and offer unique properties for engineering.

In addition to modifying or re-engineering native prokaryotic compartments, several orthogonal methods to organize the bacterial cytoplasm have been developed. Protein and nucleic acid scaffolds offer an alternative method for cellular organization without compartmentalization (*3, 8, 9*). Dueber et. al., demonstrated a 77-fold improvement in product titer by controlling the stoichiometry and recruitment of mevalonate biosynthetic enzymes to a scaffold using protein-protein interaction domains (*3*). RNA scaffolds have similarly been used to improve metabolic flux through pentadecane and succinate production pathways (*10*). Besides scaffolds, intrinsically disordered proteins have also been engineered in bacteria to confer spatial organization. In particular, phase separation of artificial polypeptides such as elastin-like polypeptides (ELPs) - which undergo simple coacervation at temperatures above their lower critical solution temperature (LCST) - has been used to form both synthetic vesicles (*11*) and simple coacervates (*12*) in *E. coli*. However, despite numerous publications over the last decade investigating the structure and function of endogenous condensates, our understanding of this nascent field is limited; consequently, engineering the *a priori* formation and dissolution of membraneless organelles *in vivo* remains a challenge, particularly in bacteria.

As a novel strategy for bacterial compartmentalization, we aim to engineer complex coacervation in *E. coli* by employing physical principles from polymer physics and inspiration from eukaryotic biomolecular condensates. Complex coacervation is driven by the electrostatic attraction between oppositely-charged macromolecules and the entropic release of bound counterions (*13*). This electrostatically driven phase separation is reported to play a key role in the formation of several membraneless organelles (*14-16*). Biomolecular condensates have been reported to form through protein-protein interactions between intrinsically disordered regions (IDRs) and through protein-RNA interactions (*17*). Given that a significant portion of biological condensates arise from RNA-protein interactions, we hypothesize that protein/RNA coacervates could be formed in bacteria to mimic naturally existing biomolecular condensates. The recent discovery of endogenous bacterial condensates further supports the feasibility of engineering synthetic membraneless organelles in *E. coli* through complex coacervation.

In this study, we use protein phase separation to create distinct compartments in *E. coli*, identify design criteria for the formation of such compartments, and evaluate compartment composition and dynamics. We begin by simplifying macromolecular interactions in the crowded intracellular milieu to solely electrostatic interactions between anionic RNA and a cationic protein. Using an engineered panel of supercationic green fluorescent protein (GFP) with positive charges isotropically distributed across the protein surface, we explore the propensity of supercationic GFP variants to undergo complex coacervation with RNA (*18*). We demonstrate the phase separation of engineered supercharged proteins with RNA under conditions that mimic the intracellular environment. Our understanding of parameters that govern phase separation *in vitro* guides our investigation and characterization of coacervate formation in cells. Our data indicates that the formation of engineered complex coacervates in *E. coli* is dependent on protein surface charge. Intracellular condensates also exhibit unique solubility behavior and GFP enriched in the coacervates freely diffuses through the surrounding cytoplasm. Finally, we find that engineered GFP coacervates are selective in their partitioning of specific nucleic acids and endogenous proteins, altogether highlighting their potential application as functional, synthetic membraneless organelles.

## Results & Discussion

### Design and *in vitro* demonstration of synthetic biomolecular condensates

We defined a simplified system for probing the phase behavior of protein condensates by first studying the *in vitro* behavior of a pair of oppositely-charged biomacromolecules consisting of engineered cationic GFP and an anionic biopolymer (Figure 1A). RNA was chosen as the anionic partner because it is distributed throughout the bacterial cytoplasm and comprises approximately 20% of dry cell weight. Additionally, RNA-protein interactions have been shown to regulate phase separation *in vitro* and in cells (*19, 20*).

**Figure 1.**
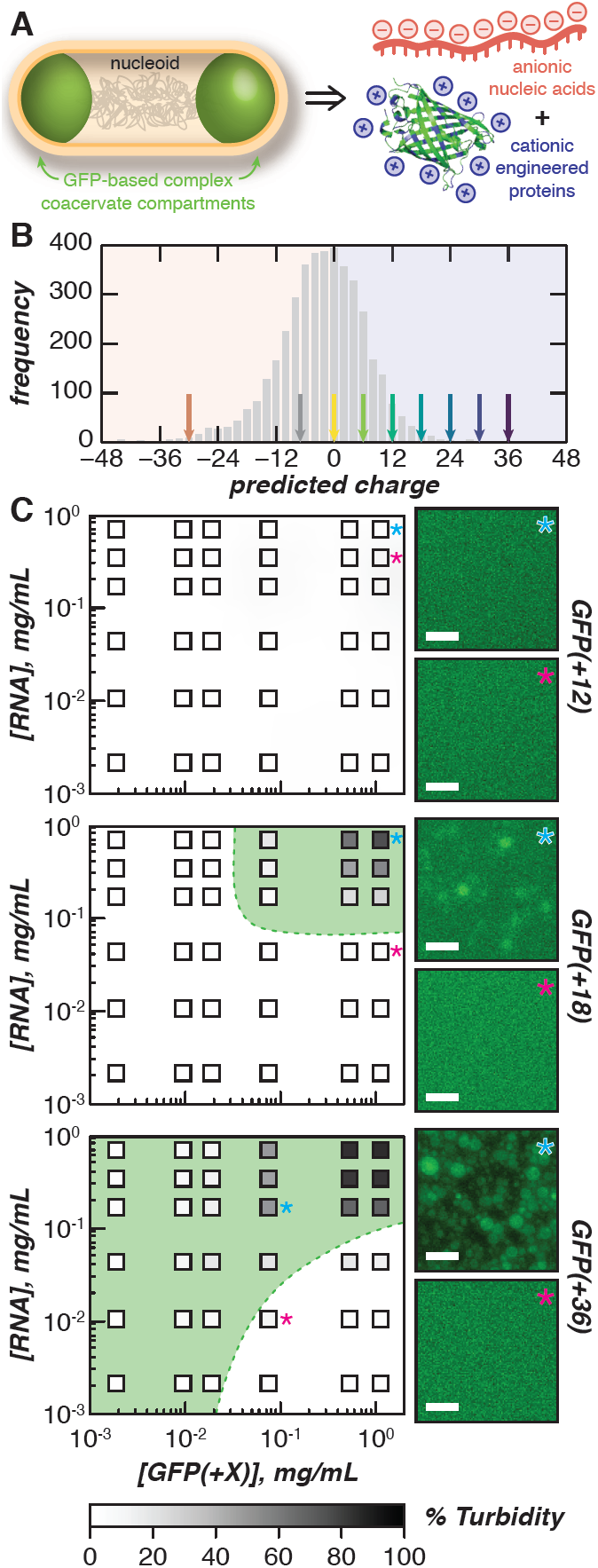
Phase separation of engineered proteins *in vitro*. (**A**) Schematic for the design of intracellular complex coacervates in *E. coli* between anionic nucleic acids and cationic engineered proteins. (**B**) Distribution of proteins in the *E. coli* proteome (UP000002032) by expected charge (bin width = 2). Arrows indicate the predicted charge of engineered GFPs used in this study. (**C**) Phase diagrams of purified GFP variants with purified total cellular RNA mixed at the indicated concentrations (boxes) in a physiological buffer (70 mM K_2_HPO_4_, 60 mM KCl, 40 mM NaCl, pH 7.4) as determined by turbidity (left); shading (within boxes) depicts turbidity values; green dashed lines represent observed phase boundaries; green shading represents two phase regions. Fluorescence microscopy images of indicated mixtures (right). Phase diagrams for GFP(+24) and GFP(+30) can be found in the supplementary information. Scale bars, 10 *µ*m.

To study the effects of protein charge on the coacervation of biomacromolecules, we used a panel of 7 isotropically supercharged GFP variants (*18*). Briefly, 3-6 amino acid substitutions were introduced at negative or neutral surface residues on superfolder GFP (sfGFP) to successively generate positively supercharged GFP variants in charge increments of +6. Using this panel of GFP variants, we were able to test the phase behavior of proteins with a range of charges that span those observed in the *E. coli* proteome (Figure 1B), with the most supercharged variant bearing a charge of +36.

We began our *in vitro* demonstration by probing the extent of phase separation of a simplified protein-RNA system using a turbidity assay. In our simplified model, we accounted for the most abundant free ions in the intracellular environment – potassium, sodium, and chloride, which are all found at millimolar concentrations in the cell. Similarly, we included phosphates as free ions because phosphates are also present at millimolar concentrations in the cell as components of nucleic acids and small metabolites. Each supercharged GFP variant was mixed with total RNA (torula yeast type VI) at various GFP/RNA ratios. We explored the phase boundary of different GFP variants by testing 6 different RNA and protein concentrations spanning 2 orders of magnitude. Turbidity measurements were collected immediately after mixing (Figure 1C and Figure S3). These *in vitro* assays were performed in a buffer (70 mM potassium phosphate dibasic, 60 mM potassium chloride, and 40 mM sodium chloride adjusted to pH 7.4) that replicated the concentration of common ions found in the intracellular environment of *E. coli*.

Initial turbidity assays performed by mixing each GFP variant with RNA at a fixed total macromolecule concentration (1 mg/mL) demonstrated that variants with an expected net charge below +18 did not phase separate with RNA under simulated physiological conditions (Figure S3). As a result, phase diagrams were investigated for GFP(+18) and variants with higher net charge in order to observe changes in phase boundaries with increased supercharging. As a negative control, phase diagrams were also constructed for GFP(+12). Consistent with our initial assays, very low turbidity values were observed at all concentrations tested for GFP(+12). In contrast, supercationic variants with higher net charges phase separated over a broader range under the concentrations and conditions tested (Figure 1C, Figure S3). This is depicted by the increase in the high turbidity regions as GFP charge is increased from +18 to +36, indicating that supercationic variants with higher net charge phase separate with RNA at lower total macro-molecule concentrations.

Optical microscopy was used to corroborate turbidity measurements and examine the nature of phase separation at all mixing ratios. Consistent with our turbidity assay, GFP(+12) did not phase separate at all of the mixing ratios tested (Figure 1C). In contrast, the formation of spherical droplets and droplet coalescence along the bottom of the plate were observed for GFP(+18) and variants of higher charge at a wider range of macromolecular concentrations (Figure 1C). Broadening of high turbidity regions is supported by the shifting of the phase boundary to include regions of lower macromolecule concentrations.

These supercationic proteins and RNA coacervate *in vitro* under conditions that mimic the intracellular environment and form liquid droplets that fuse, coalesce, and wet surfaces. We then wanted to investigate if these findings translated *in vivo*. We hypothesized that co-opting cellular machinery to produce supercationic GFP variants would be sufficient for intracellular phase separation without additional engineering.

### *In vivo* demonstration of protein condensation driven by electrostatics

Since RNA and negatively charged proteins comprise a significant portion of intracellular macromolecules, we hypothesized that expression of supercationic GFP variants alone could induce the formation of subcellular micro-assemblies *in vivo*. We further hypothesized that this could be accomplished without having to introduce an exogenous anionic partner or without having to append a phase separating domain, such as an intrinsically disordered polypeptide (IDP). Finally, we hypothesized that the formation of cellular compartments could be predicted by our simple *in vitro* protein-nucleic acid model. We report that expressing supercharged GFP (≥ +12) alone is sufficient to form sub-micron sized compartments in *E. coli* (Figure 2A). These phase separated compartments represent local intracellular regions containing higher GFP concentrations than the surrounding cytoplasm.

**Figure 2.**
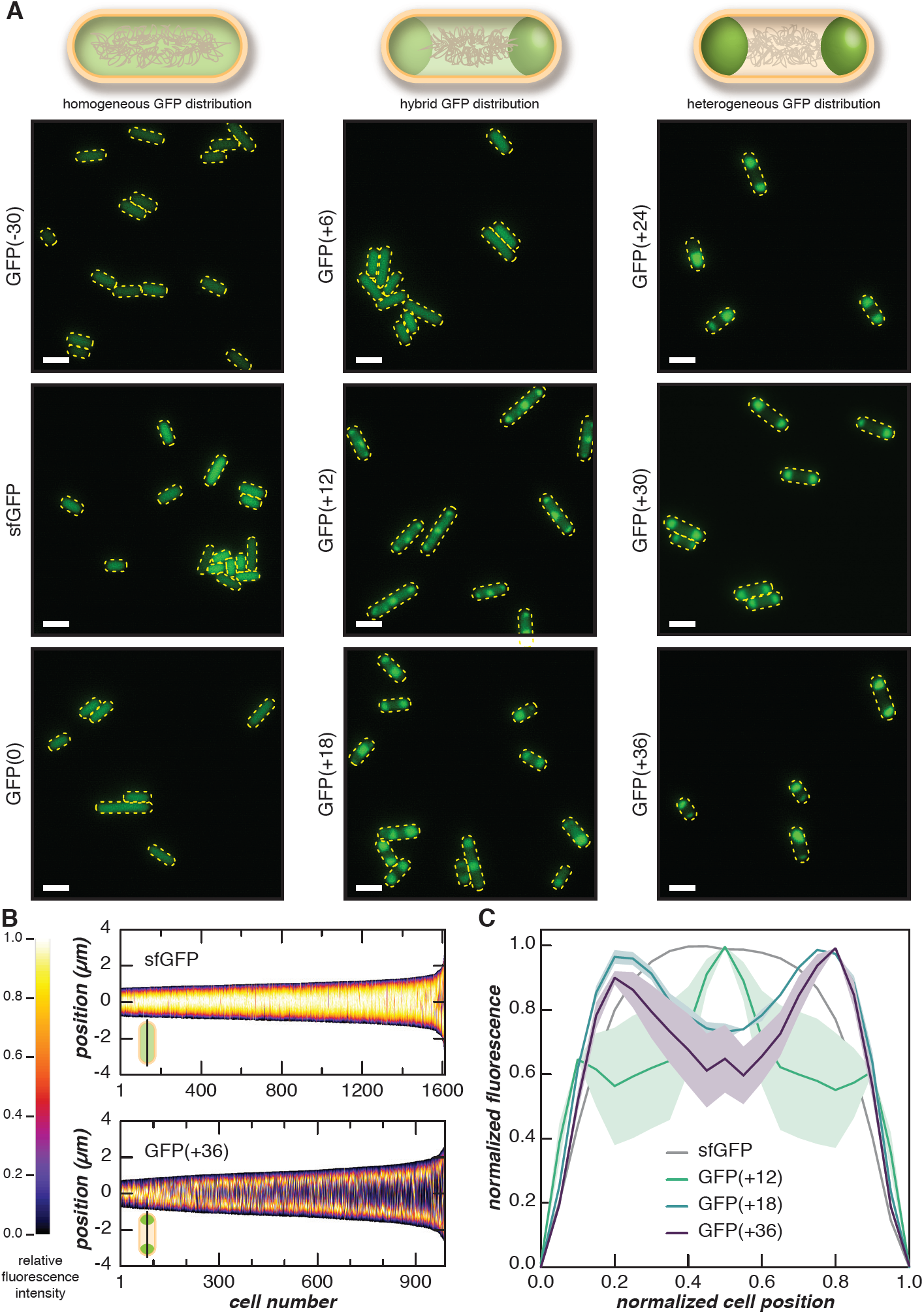
Phase separation of supercationic GFP in *E. coli*. (**A**) Fluorescent microscopy images of cells expressing GFP variants with different net charge at 24 h post-induction. Negatively charged or neutral variants (left column) were evenly distributed throughout the cell, while supercationic variants demonstrated punctate fluorescence localized to the cell poles (right column). The localization of fluorescence was more defined with increasing cationic charge as exemplified by the charge-dependent increase in localization observed with GFP variants of intermediate charge (middle column). Scale bars, 2 *µ*m. (**B**) Localization patterns of sfGFP and GFP(+36) were quantified from microscopy images, such as those shown in **A**. Vertical heatmaps representing GFP intensities across the long cell axis were generated using microbeJ. Demographs display the GFP intensity across a population of cells arranged by cell length. (**C**) The localization of GFP when normalized to cell position demonstrated transitions as GFP charge reaches +12 and +18. Each line represents the normalized medial fluorescence with respect to normalized cell position and the shaded region represents the SEM of observed values. For **B** and **C** three independent experiments were performed with at least 120 cells analyzed per experiment. Analysis for all GFP variants is found in Figures S4-9.

To test the dependence of phase behavior on protein charge, we over-expressed each supercationic GFP variant in *E. coli* cells (NiCo21(DE3)) and imaged the cells by optical microscopy at various time points after induction. Compartment formation is observed primarily in cells expressing GFP with a net charge of +12 or higher, which is largely consistent with trends observed *in vitro* through both turbidity assays and microscopy (Figure 2). In contrast, cells expressing a negatively charged superfolder GFP and variants with a net charge of +6 or below exhibit an even distribution of GFP across the length of the cell (Figures S4-9). To control for the effects of protein supercharging on intracellular compartment formation, we expressed a superanionic-GFP variant with a charge of −30. We observed an even GFP distribution within each cell, suggesting that merely supercharging is not sufficient for condensate formation and that supercationic proteins are required (Figures S4-9). These observations were then quantified by analyzing the spatial distribution of each GFP variant along the medial cell axis (Figure 2B). Distinct differences in spatial GFP distribution were exemplified by sf-GFP and GFP(+36). Image analysis confirmed homogeneous sfGFP distribution across the length of the cell whereas GFP(+36) was concentrated primarily at the poles.

Our results further demonstrated that GFP distribution in the cell is dependent on protein charge. Heterogeneous GFP distribution became more distinct with increasing protein net charge as shown by the decrease in fluorescence intensity at the cell center of the intermediate charge GFP variants (Figure 2A and Figures S4-9). A transition in GFP localization was also observed with increased supercharging, whereby expression of GFP(+6) in cells resulted in homogeneous GFP distribution at 24 h post-induction whereas GFP(+12) in cells produced 3 intracellular condensates and the higher supercharged variants produced only 2 condensates (Figure 2C).

Formation of phase separated condensates was observed to be dependent on the duration of protein expression in addition to the protein charge. Cells expressing GFP(+6) were imaged and analyzed at 2, 8, and 24 h after induction of GFP expression. During this time, an increase in intracellular GFP concentration was observed (Figure S10) and a transition between homogeneous distribution to heterogeneous distribution was observed from 2 h to 8 h. As intracellular GFP(+6) concentration continued to increase, the cells transitioned back to a uniform distribution at 24 h post-induction (Figures S4-9). Moreover, all GFP variants at 2 hours exhibited either a uniform distribution throughout the cytoplasm or an altered distribution in which compartments had formed but GFP was largely observed throughout the cytoplasm. We denote the altered distributions as “hybrid” distributions, representing regions on a phase diagram near the critical point where the difference in GFP concentration between the dilute and coacervate phase is minimal. Transitions between different GFP distributions may be explained by the change in intracellular GFP concentration over time as phase separation is dependent on macromolecule concentration as demonstrated *in vitro*. Changes in cytoplasmic macro-molecule concentrations over the cell growth cycle during the course of the experiment allow the cell to traverse phase boundaries (Figure S11).

Intracellular phase separation had a minimal impact on both cell viability and expression of most super-charged GFP variants. Growth assays conducted at 25 °C revealed that cells expressing supercationic GFP grew similarly to those expressing sfGFP for the first ∼8 h after induction of protein expression. All cationic variants then showed slightly depressed growth at longer time points (Figure S10). Importantly, condensate formation did not impact cell morphology and resulted in minimal differences in cell length (Figures S12-15). Moreover, fluorescence intensity measurements normalized to cell density increased for all GFP variants for 24 h post-induction (Figure S4). In general, increases in this normalized fluorescence over time was comparable between all GFP variants with the exception of GFP(+30) and GFP(+6). GFP(+30) consistently demonstrated the lowest optical density at all time points and exhibited the lowest change in the normalized fluorescence intensity over 8 and 24 h (Figure S10). Additionally, condensates formed in cells expressing GFP(+30) at 8 h were much less distinct than those found in cells expressing GFP(+24) and GFP(+36) (Figures S5 & S7). Demographs of GFP(+30) at 8 h depicted GFP(+30) concentrated at the poles and separated by a middle region containing GFP concentrations higher than observed for GFP(+24) and GFP(+36). Altogether, GFP(+30) did not follow the predicted trend in intracellular condensate formation even though *in vitro* turbidity assays and phase diagrams suggested otherwise. GFP(+30) may suffer from relatively poor GFP expression in cells and does not achieve the intracellular GFP concentration required to form larger condensates. In contrast, GFP(+6) demonstrated the highest increase in normalized GFP fluorescence intensity from 2 to 8 h and another increase from 8 to 24 h. Interestingly, these changes in normalized fluorescence intensity may help explain the reversible formation of compartments in cells expressing GFP(+6) at 8 h and their dissolution at 24 h (Figure S4-9). Using the normalized fluorescence as a proxy for GFP concentration per cell, we hypothesized that the protein concentration in the cell may traverse the phase boundary to a demixed state at 8 h upon an initial increase in GFP(+6) concentration between 2 to 8 h (Figure S11). The disappearance of compartments at 24 h upon a further increase in intracellular GFP(+6) concentration then corresponds to the cellular GFP concentration crossing the phase boundary again, returning to a single phase state. This hypothesis is based on our observations of the *in vivo* formation and dissolution of condensates of GFP(+6) as well as reversible phase transitions of membraneless organelles reported in eukaryotic cells (*17*). Moreover, phase separation is difficult to test *in vitro* because phase separation of GFP(+6) with RNA is not observed in our simple *in vitro* model.

Taken together, our *in vivo* results revealed that the formation of protein-dense compartments in *E. coli* is dependent on protein charge and concentration. We also demonstrated the reversibility of intracellular compartment formation as a consequence of changing intracellular GFP concentration, providing initial evidence that the compartments may arise from protein phase separation. Moreover, the intracellular formation of protein condensates aligned with trends in our *in vitro* protein-RNA model. This indicates that our simplified model, which accounts for the ionic strength of the intracellular environment and electrostatic interactions between RNA and supercationic GFP, is predictive for engineered intracellular condensates.

### Condensed phase is distinct from inclusion bodies and is dynamic

We proceeded to characterize the properties of these protein condensates and the dynamics of the encapsulated supercharged GFP. Literature on condensate formation in bacteria remains sparse, but recent reports have shed light on endogenous RNP-bodies (known as BR-bodies) formed in *C. crescentus* from a protein, RNase E, which contains an unstructured C-terminal domain responsible for intracellular phase separation (*6, 7*). In addition to BR-bodies, numerous studies have reported both artificially engineered and common endogenous bacterial compartments (*2, 11, 21, 22*). Hollow bacterial compartments, such as vesicles, have been engineered from simple coacervation of elastin-like polypeptides (ELPs) fused with GFP (*11*). Other GFP-ELP fusions have also been reported to form phase separated compartments in *E. coli* (*12*). In addition to vesicles and coacervates, compartments of insoluble, mis-folded protein, termed inclusion bodies (IBs), can form in *E. coli* when expressing recombinant proteins, frequently complicating protein purification processes in industrial settings. Protein purification from IBs involves isolation of IBs from cells and solubilization of IBs using chemical denaturants to unfold aggregated proteins. To distinguish phase separated condensates from other bacterial compartments, condensate solubility and intracellular protein dynamics were investigated.

Coacervate-like properties of the intracellular compartments were probed by examining compartment solubility under varying conditions. We hypothesized that, as protein-based complex coacervates, intracellular condensates would be soluble in high ionic strength buffers due to the screening of electrostatic interactions by ions in solution. In contrast, charge screening would have minimal effect on the dissolution of inclusion bodies (IBs) which form by aggregation of partially folded recombinant proteins. IB solubilization would instead require denaturation of the partially and mis-folded proteins using a chaotrope such as urea.

To distinguish differences in solubility between intracellular compartments and IBs, we first engineered an inclusion body-forming supercationic GFP variant, IB-GFP(+36), by deleting a hydrophobic loop (GPVLLP) that lies outside the barrel of GFP. This mutation was previously reported to result in IB formation in cells expressing mutated GF-Phs1, a stabilized derivative of sfGFP (*23*). As an added control, the solubility of streptavidin, a protein commonly reported to form IBs when recombinantly expressed in *E. coli* (*24*), was tested using the same set of solubilization conditions. Similarities in solubility of IB-GFP(+36) and streptavidin suggest that IB-GFP(+36) forms insoluble aggregates (Figures S16-17). Additionally, IB-GFP(+36) easily photobleached in cells during microscopy and exhibited low fluorescence, which is likely due to low expression of folded IB-GFP(+36) as evident by the overall reduction in IB-GFP observed in our SDS-PAGE analysis (Figures S19 & S16). Rapid photobleaching of GFP within inclusion bodies has also been observed for GFP fusion proteins expressed in *E. coli* (*25*). We performed comparative solubility studies on GFP variants that exhibited the most prominent differences in phenotypes - sfGFP, condensate-forming GFP(+36), and inclusion body-forming IB-GFP(+36). After expression of each GFP variant in cells, cells were harvested, lysed and centrifuged to separate the soluble protein fraction in the supernatant from the dense, insoluble components in the pellet. Under standard protein purification conditions, IBs separate into the insoluble pellet and we hypothesized that dense GFP coacervates would also localize to the pellet. Pellets were washed to remove residual soluble proteins and treated with a low salt, 1 M NaCl, or 8 M urea buffer. The soluble and insoluble fractions were collected and the amount of GFP in each fraction was analyzed by SDS-PAGE (Figure 3A and Figure S16).

**Figure 3.**
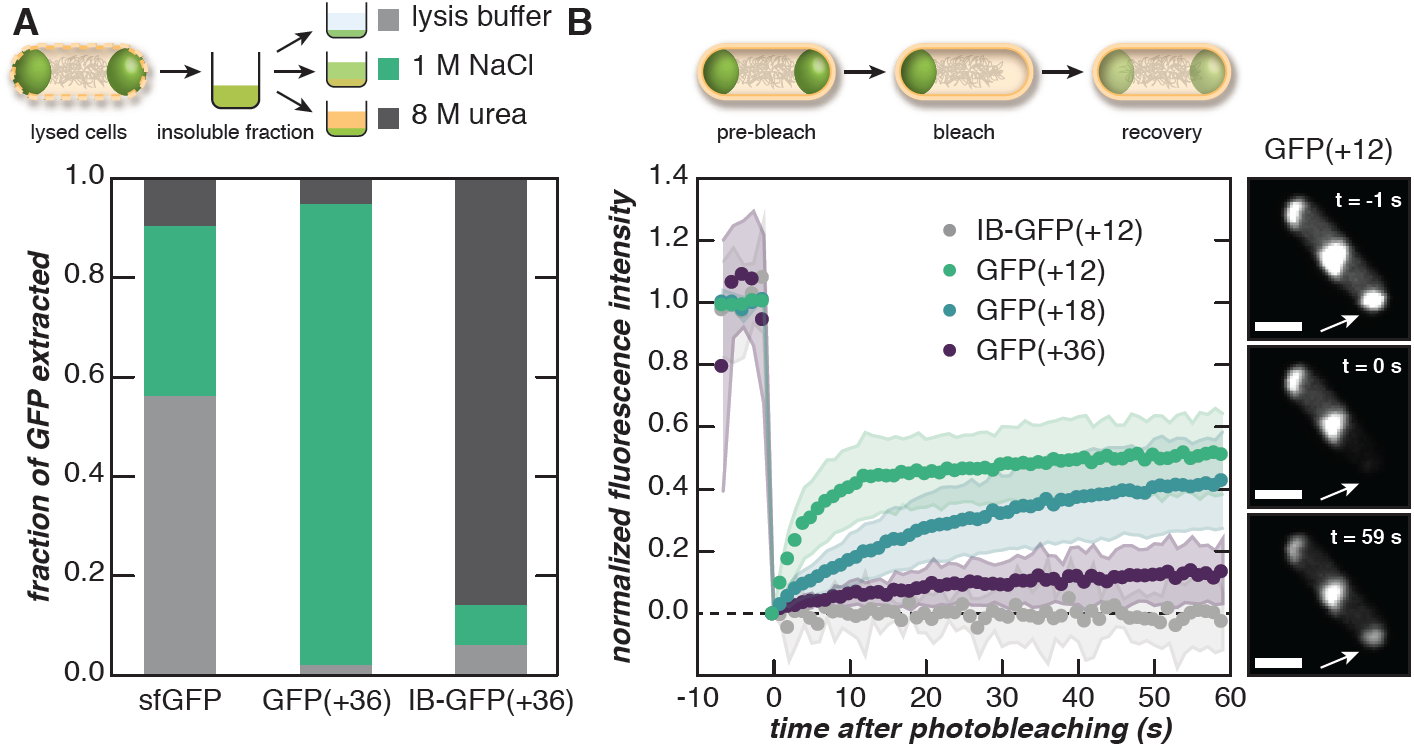
Fluorescent puncta behave as complex coacervates. (**A**) The dense, insoluble fraction of lysed *E. coli* cells expressing engineered GFPs was solubilized in a range of buffers to distinguish the behavior of supercationic GFP(+36) from an inclusion body (IB) forming variant. The insoluble fraction was treated with lysis buffer, 1 M NaCl, or 8 M urea and the fraction of GFP (sfGFP, GFP(+36), or IB-GFP(+36)) solubilized with each treatment was determined by SDS-PAGE analysis. (**B**) One pole of an *E. coli* cell was bleached and the fluorescence recovery was monitored over time (left). The panels show the fluorescence of a representative cell expressing GFP(+12) at different time points during FRAP (right). Supercharged GFP droplets were dynamic relative to IB-forming GFPs and became less dynamic with increasing protein charge. Scale bar, 1 *µ*m.

Solubility differences indicated that supercationic GFPs formed compartments distinct from inclusion bodies. Compartment-forming GFP variants were difficult to solubilize in a low ionic strength buffer whereas sfGFP was soluble in all buffers tested (Figure 3A). The fraction of GFP(+36) extracted with a high ionic strength buffer (∼0.9) was much higher than extraction with urea (∼0.05). This indicated that solubilization of these compartments required increased ionic strength, providing further evidence for complex coacervate-like properties. The formation of a single phase upon exposure to increased salt concentration is likely caused by salt screening of electrostatic interactions between oppositely charged polyelectrolytes (*13*). In contrast, the fraction of extracted IB-GFP(+36) was highest in 8 M urea buffer (∼0.9) and appreciably lower in 1 M NaCl buffer (∼0.1), demonstrating that inclusion bodies resisted dissolution in high ionic strength buffers and required protein denaturation with urea. Taken together, these solubility assays provided evidence for the coacervate-like phase behavior of the intracellular compartments formed by GFP(+36) and differentiated these protein condensates from inclusion bodies.

The protein condensates formed by intracellular GFP(+36) are GFP-dense, which is consistent with *in vitro* protein encapsulation experiments demonstrating that the coacervate phase can efficiently encapsulate >90% of proteins in protein-nucleic acid mixtures (*18*). In our solubility assays, the majority of GFP(+36) was extracted upon solubilization with 1 M NaCl and very little GFP remained in the pellet as observed by SDS-PAGE analysis (Figure S16). These results suggest that protein supercharging may provide a strategy for enriching recombinantly expressed proteins in engineered intracellular compartments. Additionally, gel analysis revealed that the vast majority of the protein solubilized under high salt conditions was GFP. These results also indicate that it is possible to achieve selective condensation without inherent biomolecular specificity (*26*).

To further demonstrate that compartments formed by supercharged GFP are condensates capable of dynamic restructuring, fluorescence recovery after photobleaching (FRAP) was conducted to monitor the diffusion of GFP molecules between compartments. Cells were bleached at one of the poles and fluorescence intensity of the bleached condensate was monitored over time (Figure 3B). GFP(+12) exhibited the fastest average recovery (t1/2 ∼4 s) and highest average mobile fraction (∼0.5) as estimated by a single exponential fit (Figures S18 & S20). Additionally, the recovery of GFP fluorescence could be visually observed in microscopy images approximately 1 minute after bleaching and fluorescence of adjacent condensates within the same cell were visibly decreased, indicating diffusion of GFP between condensates. GFP diffusion between condensates was reduced as protein supercharging increased. The mobile fraction decreased and the half-life of recovery increased with increasing protein net charge (Figure 3B).

To further distinguish compartments formed from supercationic proteins, the dynamics of inclusion body-forming GFP variants and sfGFP were also tested. IB-GFP(+12) was used as a negative control and did not recover after bleaching. Similarly, additional negative controls also photobleached easily and did not recover (see IB-GFP(+18) and IB-GFP(+36) in Figures S19-20). This is consistent with other reports that have attempted photobleaching of intracellular inclusion bodies in mammalian cells and observed no material exchange (*27*). Cells expressing sfGFP showed immediate, complete bleaching and exhibited no visible recovery (Figures S18 & S20). Recovery was not observed due to the rapid diffusion of GFP, which resulted in bleaching of the entire cell. This is consistent with previous reports measuring GFP diffusion in the *E. coli* cytoplasm, where complete fluorescence recovery occurs within 1.5 seconds following the bleaching event (*28, 29*).

Taken together, protein solubility and FRAP experiments demonstrate that compartments formed by GFP(+36) likely arise from electrostatic interactions consistent with complex coacervation. Solubility assays revealed that the coacervate phase is dense and requires a dissolution mechanism different from that of common, insoluble bacterial inclusion bodies. Moreover, analysis by FRAP confirmed that the condensed phase is capable of material exchange through the surrounding cytoplasm and the dynamic exchange is dependent on protein charge.

### Identification of endogenous biomolecules in engineered protein condensates

Since endogenous nucleic acids and proteins participate in and regulate intracellular condensation (*19, 30-32*), we hypothesized that in addition to supercationic GFP, endogenous biomacromolecules would be localized to the engineered condensates. To characterize condensate composition, a combination of nucleic acid staining and proteomics assays were performed to identify constituents that participate or partition into the condensate phase. The roles of nucleic acids in driving protein phase separation have been reported both *in vivo* and *in vitro* (*17, 18, 31, 33, 34*).

Nucleic acid constituents of the protein condensates were investigated by imaging cells expressing GFP(+36) with either a DNA stain (DAPI) or a general nucleic acid stain (SYTO17) that binds to both DNA and RNA. DAPI staining revealed that DNA was excluded from the coacervate phase (Figure 4A). DNA localized to the center of the cell and was excluded from GFP condensates at the cell poles. However, spatial overlap between GFP(+36) condensates and SYTO17 dyes suggest that, unlike DNA, RNA colocalizes with the protein condensates and may be a constituent (Figure 4B, Figure S21). In contrast both DAPI and SYTO17 staining of cells expressing sfGFP depict fluorescence throughout the entire cell, indicating colocalization of sfGFP with both DNA and RNA.

**Figure 4.**
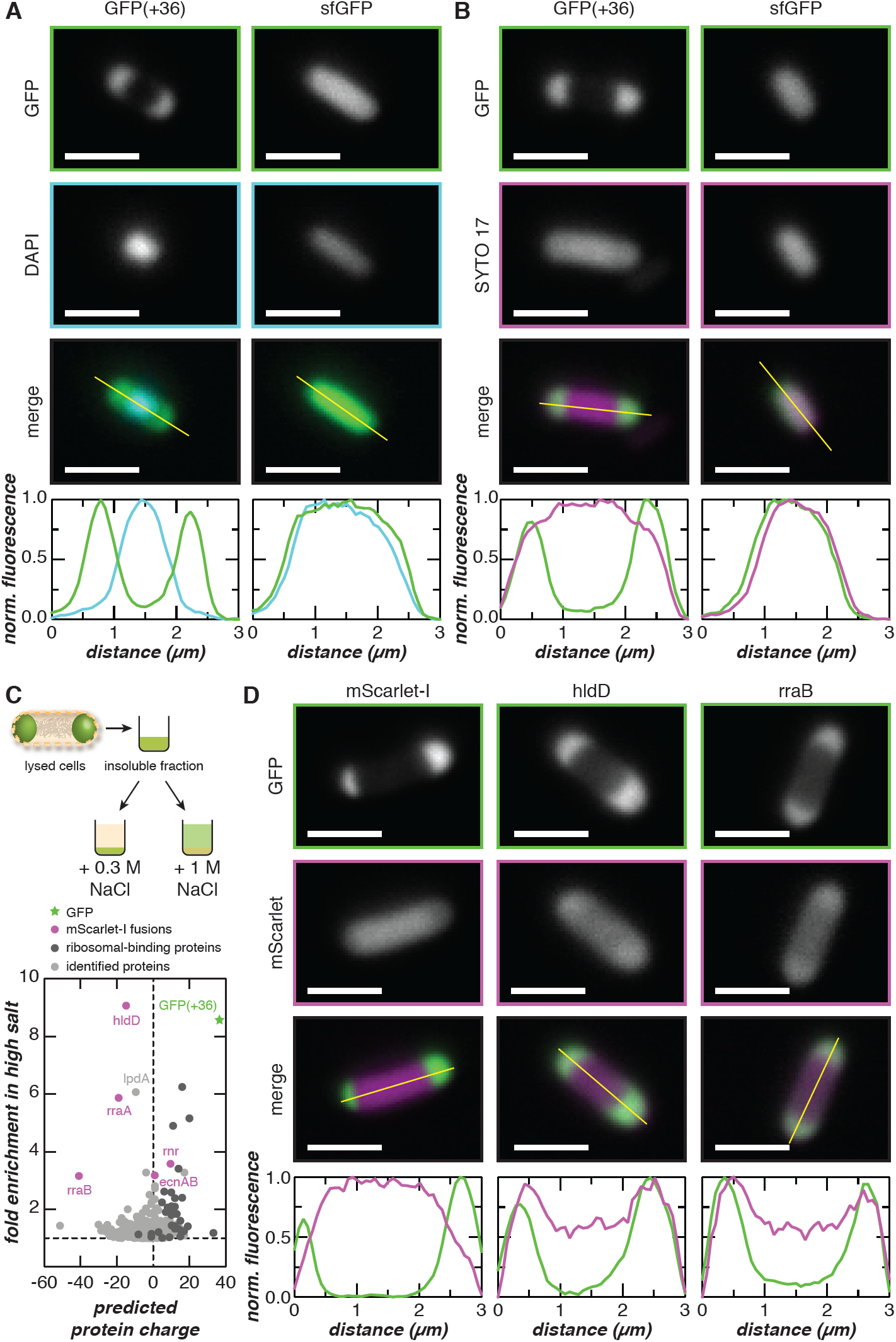
Colocalization of endogenous biomolecules with supercationic GFP condensates. (**A**) The colocalization of DNA with the GFP(+36) condensates was evaluated by staining cells with DAPI. Microscopy images depict cells expressing GFP(+36) or sfGFP and stained with DAPI. Intensity line-cuts demonstrate exclusion of DNA from the GFP(+36) condensates. (**B**) The colocalization of RNA with the GFP(+36) condensates was evaluated by staining cells with SYTO 17, which binds both DNA and RNA. Microscopy image of cells expressing GFP(+36) or sfGFP and stained with SYTO 17 are shown along with intensity line-cuts demonstrating colocalization of RNA and GFPs. (**C**) Proteins from lysed *E. coli* cells enriched following high salt fractionation were quantified by LC-MS/MS. The fold enrichment under high salt treatment is plotted against the predicted protein charge. (**D**) Selected proteins from **C** were co-expressed with GFP(+36) as mScarlet-I fusions. Fluorescence microscopy and intensity line-cuts demonstrate colocalization of GFP(+36) with hldD and rraB mScarlet-I fusions. Additional co-expression data is found in Figure S23. Scale bars, 2 *µ*m.

While nucleic acids have been reported as major participants in biomolecular phase separation, weak, transient interactions between protein domains have also been shown to give rise to biomolecular condensates (*17*). Moreover, an analysis of the *E. coli* proteome reveals that the frequency distribution of protein expected charge (Figure 1B) is skewed such that the proteome is net negative. As a result, we hypothesized that proteins may also function as anionic counterparts to the engineered supercationic GFPs.

Quantitative proteomics was performed to identify protein constituents of the condensates formed by GFP(+36). Proteins extracted from the insoluble fraction containing the GFP(+36)-based condensed phase under high salt conditions were compared to those isolated via low salt extraction (Figure S22). Because the high concentration of GFP(+36) in the condensate could mask the presence of other proteins, GFP(+36) was depleted from the samples using affinity chromatography. Approximately 1100 proteins were identified by proteomics with ∼450 proteins showing slight enrichment in the extracted condensates (>1-fold) and only ∼30 proteins demonstrating significant enrichment (>2-fold) (Figure 4C; Supplementary Data S1). Many ribosomal binding proteins (∼30) were identified but were excluded as candidates for validation experiments because they are easily extracted from the insoluble fraction through mild salt fractionation (*35*). Both positively and negatively charged proteins were enriched in the condensate, with an anionic protein exhibiting the highest (∼9-fold) enrichment.

To validate our quantitative proteomics results, several identified proteins were genetically fused to mScarlet-I to evaluate intracellular colocalization with GFP(+36) condensates. Proteins chosen for validation experiments, including ADP-L-glycero-D-manno-heptose-6-epimerase (hldD), regulator of ribonuclease activity A (rraA), ribonuclease R (rnr), regulator of ribonuclease activity B (rraB), entericidin A/B (ecnAB), and biosynthetic arginine decarboxylase (speA) (*36*), spanned nearly two orders of fold-enrichment (Figure 4C). To allow for orthogonal gene expression, mScarlet-I fusions were assembled onto an arabinose-inducible plasmid and expressed in *E. coli* along with an IPTG-inducible GFP(+36) plasmid. Of the protein fusions tested, two endogenous *E. coli* proteins colocalized with GFP condensates. When fused to mScarlet-I, the protein exhibiting the highest fold change (hldD) and the most anionic protein tested (rraB) colocalized with GFP(+36) condensates (Figure 4D). Interestingly, hldD is involved in lipopolysaccharide core biosynthesis and has been reported to promote *E. coli* viability under high temperatures when induced by heat shock (*37*). Parallels can be drawn to proteins in eukaryotic cells that mount adaptive responses upon exposure to environmental stressors by forming biomolecular condensates (*19*). In the case of a polyA-binding protein (Pab1) in yeast, phase separation of Pab1 through protein-protein interactions improved organism fitness under prolonged thermal stress (*38*). Another protein tested, rraB regulates intracellular RNA abundance by inhibiting RNase E and preventing degradation of specific RNA transcripts (*39, 40*). More excitingly, RNase E has been implicated in the formation and degradative function of recently discovered bacterial condensates (BR bodies) that sequester and control the degradation of RNA in *C. crescentus* (*6*). The same study also showed that BR bodies improved cell growth in response to acute ethanol stress. More comprehensive studies are required to understand the presence and function of hldD and rraB in our engineered GFP condensates. However, their functional similarities to proteins responsible for intracellular phase separation in other organisms also indicate that hldD and rraB are promising candidates for possible endogenous biomolecular condensates in *E. coli*.

In contrast, rraA-mScarlet-I did not show a spatial preference and exhibited a distribution profile similar to the control sample, in which mScarlet-I alone was co-expressed with GFP(+36) (Figure 4D; Figure S23). The co-expression control demonstrated homogeneous distribution of mScarlet-I throughout the cell and heterogeneous distribution of GFP(+36) to the poles. These results indicated that mScarlet-I expression did not affect GFP(+36) localization and that mScarlet-I co-localization at the poles in hldD- and rraB-mScarlet-I samples was not an artifact of our expression system. Moreover, like rraB, rraA controls mRNA abundance by binding and inhibiting RNaseE activity; however, rraA regulates a set of RNA transcripts distinct from those of rraB (*41*). The lack of spatial preference suggests that the rraA fusion may have a lower preference for interactions with constituents of the coacervate phase than its rraB counterpart.

The remaining protein fusions (ecnAB-, speA-, and rnr-mScarlet-I) did not colocalize with GFP(+36) condensates. Since ecnAB is a bacterial lipoprotein that localizes to the cell membrane (*42*), the mScarlet-I fusion exhibits fluorescence that flanks the GFP condensates, representing the cell border (Figure S23). SpeA, was used as a negative control because it was identified by proteomics but was not enriched in the condensate while being superanionic (expected charge of −41). As predicted, speA-mScarlet-I demonstrated anti-colocalization whereby the fusion protein was predominantly localized to the center of the cell and partially excluded from the poles. Surprisingly, rnr-mScarlet-I was also observed to be excluded from the cell poles despite it’s ∼3.6-fold enrichment. Rnr has a positive expected charge of 9.64 at physiological pH. Since condensates are primarily comprised of supercationic GFP, we hypothesized that partitioning of rnr-mScarlet-I into the coacervate was disfavored because it would be outcompeted by supercationic GFP for attractive interactions with anionic macromolecules in the condensate. Moreover, similar to the ribosomal binding proteins identified by proteomics, rnr may be easily solubilized in the presence of high salt, which could explain its observed enrichment in our proteomics screen.

As additional controls, mScarlet-I fusion proteins were co-expressed with sfGFP (Figure S23). All mScarlet-I fusions demonstrated homogeneous fluorescence intensity distributions that correlated with the homogeneous distribution of sfGFP, with the exception of ecnAB-mScarlet-I, which localized to the membrane. Similarities in spatial distribution profiles provided further evidence that colocalization of endogenous genes with condensates were not an artifact of our chosen expression system. It should be noted that coexpression of speA-mScarlet-I with sfGFP often yielded cells that were much wider and longer than cells from other co-expressions. Additionally, a small percentage of cells expressing speA-mScarlet-I exhibited very low sfGFP fluorescence and exclusion of the fusion protein from non-specific regions within the cell. However, co-expression of speA-mScarlet-I with GFP(+36) significantly minimized the presence of these anomalous cells and reduced cell size to that of the mScarlet-I control. This suggested that the cellular burden of synthetically expressing speA-mScarlet-I was reduced when co-expressed with GFP(+36) and that the observed anti-colocalization was not an experimental artifact. Validation of additional endogenous proteins exhibiting low fold-enrichment may help garner further insights into protein properties that promote intracellular complex coacervation.

*In vivo* nucleic acid staining and quantitative proteomics experiments demonstrated that synthetic GFP condensates are selective in their nucleic acid and protein composition. Colocalization analysis of nucleic acid dyes with GFP condensates depicted the exclusion of DNA and incorporation of RNA into the compartments, supporting our simple *in vitro* RNA-protein model (Figure 1A). Moreover, identification of protein constituents by high salt fractionation and quantitative proteomics revealed that highly charged endogenous proteins are present in the condensates along with RNA. Validation of proteomics candidates spanning a range of fold-enrichment and expected charges provides further evidence for the participation and/or partitioning of highly charged endogenous proteins into the coacervate phase.

## Conclusion

In summary, we have developed a promising method for engineering dynamic condensates in *E. coli* that can enrich heterologously expressed proteins by increasing protein surface charge. To our knowledge, the engineering of complex coacervates in bacteria has not been attempted. We demonstrate here that condensate formation is dependent on the extent of protein supercharging and intracellular concentration of the protein expressed. Characterization of the condensates suggest that they are held together by electrostatic interactions and are composed of RNA and endogenous protein, while excluding DNA. Moreover, the propensity for each supercharged GFP variant to form condensates can be approximated by its *in vitro* phase behavior when mixed with total RNA. The use of supercharged proteins for engineered complex coacervation *in vivo* has great potential for the functional engineering of intracellular condensates as bioreactors, intracellular protein depots, and biosensors.

## Acknowledgements

Images were collected in the Confocal and Specialized Microscopy Shared Resource of the Herbert Irving Comprehensive Cancer Center at Columbia University, supported by NIH grant #P30 CA013696 (National Cancer Institute). The confocal microscope was purchased with NIH grant #S10 RR025686. We also acknowledge Katia Kovrizhkin for her preliminary work on constructing phase diagrams. We thank Shahar Goeta for preparing samples for mass spectrometry and for processing mass spectrometry data files.

## Author contributions

V.Y. and A.C.O. conceived of the project. V.Y. performed *in vitro* and *in vivo* experiments. V.Y. and A.C.O. analyzed the data. L.M.B. and E.G.W conducted the proteomics experiments and analysis. V.Y. and A.C.O. wrote the manuscript. All authors edited and approved the final manuscript.

## Data and materials availability

All data is available in the manuscript or the supplementary materials. Raw data and python code used to generate heatmaps depicted in phase diagrams are available upon request. Mass spectrometry raw data files have been deposited in two public repositories, Chorus (https://chorusproject.org under project ID# 1656) and the MassIVE database (ftp://massive.ucsd.edu/MSV000085333/ with doi:10.25345/C5297W).

