## Supplementary Materials for "Formation of biomolecular condensates in bacteria by tuning protein electrostatics"

**This PDF file includes:**

Materials and Methods
Supplementary Text
Figures S1 to S23
Caption for Data S1

**Other Supplementary Materials for this manuscript include the following:**

Data S1 [Protein enrichment in the condensate as determined by quantitative proteomics]

### Materials and Methods

Cloning of supercharged GFP variants

Supercharged GFP variants described in this paper were previously purchased from GenScript and all contain an N-terminal 6xHis tag (*18*). Briefly, site-specific mutations were introduced, replacing neutral (Ile, Leu, Ala, Val, Thr, Met, Asn, Gln) and acidic (Glu, Asp) residues on surface of superfolder GFP (sfGFP) with basic (Arg, Lys) residues to increase the overall net positive charge on the protein. Mutation sites were chosen from a set of mutations previously reported to generate highly supercharged superfolder GFP with an expected charge of (+36) (*43*). The crystal structure of sfGFP (PDB ID: 2B3P) was used as a guide to evenly distribute charges across the protein surface to minimize the introduction of charge patches. As a control, superanionic GFP(-30) was created using mutations described by the Liu group (*43*). The inclusion body (IB)-forming GFP variants (IB-GFP(+12), IB-GFP(+18), and IB-GFP(+36)) were generated by deleting the nucleic acid sequence encoding GPVLLP (loop 32). Deletions were performed as instructed using NEBuilder HiFi DNA assembly and primers were purchased through Integrated DNA Technologies. PCR products were amplified using Phusion high-fidelity DNA polymerase (NEB), and were purified using QIAquick PCR purification kit or QIAquick gel extraction kit (Qiagen) prior to assembly with NEBuilder HiFi DNA assembly master mix. Transformations into NiCo21(DE3) and NEB5α cells were performed as instructed by NEB. All *in vivo* experiments were conducted using NiCo21(DE3) cells purchased from NEB. All GFP variants were cloned with an N-terminal 6xHis-tag.

Primer sequences and templates used for HiFi assembly of IB-GFP variants are provided below. All primers were purchased from IDT.

*IB-GFP(+12):*
Template for insert and vector PCRs: GFP(+12)
Insert forward primer: 5' – tgcgttgcgcTCACTGCCCGCTTTCCAG – 3'
Insert reverse primer: 5' – agtggttgcgATCACCAATCGGCGTGTTTTG – 3'
Vector forward primer: 5' – gattggtgatCGCAACCACTACCTGAGC – 3'
Vector reverse primer: 5' – cgggcagtgaGCGCAACGCAATTAATGTAAG – 3'

*IB-GFP(+18):*
Template for insert and vector PCRs: GFP(+18)
The same primers used to construct IB-GFP(+12) were also used to construct IB-GFP(+18).

*IB-GFP(+36):*
Template for insert and vector PCRs: GFP(+36)
Insert forward primer: same insert forward primer for IB-GFP(+12)
Insert reverse primer: 5' – agtggttgcgGCGACCAATCGGCGTGTT – 3'
Vector forward primer: 5' – gattggtcgcCGCAACCACTACCTGAGC – 3'
Vector reverse primer: same vector reverse primer for IB-GFP(+12)

Amino acid sequences of all GFP variants are provided below. Mutations from the parent sequence are bolded and underlined.

*sfGFP:*
MGHHHHHHGGASKGEELFTGVVPILVELDGDVNGHKFSVRGEGEGDATNGKLTLKFICTTGKLPVP-WPTLVTTLTYGVQCFSRYPDHMKQHDFFKSAMPEGYVQERTISFKDDGTYKTRAEVKFEGDTLVNRI-ELKGIDFKEDGNILGHKLEYNFNSHNVYITADKQKNGIKANFKIRHNVEDGSVQLADHYQQNTPIG-DGPVLLPDNHYLSTQSALSKDPNEKRDHMVLLEFVTAAGITHGMDELYK

*GFP(0):* (mutations to sfGFP)
MGHHHHHHGGASKGE**R**LFTGVVPILVELDGDVNGHKFSVRGEGEGDATNGKLTLKFICTTGKLPVP-

WPTLVTTLTYGVQCFSRYPDHMKQHDFFKSAMPEGYVQERTISFKDDGTYKTRAEVKFEGDTLVNRI-
ELKG**R**DFKEDGNILGHKLEYNFNSHNVYITADKQKNGIKANFKIRHNVEDGSVQLADHYQQNTPIG-
DGPVLLP**R**NHYLSTQSALSKDP**K**EKRDHMVLLEFVTAAGITHGMDE**R**YK

*GFP(+6):* (mutations to GFP(0))
MGHHHHHHGGASKGERLFTGVVPILVELDGDVNGHKFSVRGEGEGDATNGKLTLKFICTTGKLPVP-
WPTLVTTLTYGVQCFSRYP**K**HMKQHDFFKSAMPEGYVQERTISFK**K**DGTYKTRAEVKFEG**R**TLVN-
RIELKGRDFKEDGNILGHKLEYNFNSHNVYITADKQKNGIKANFKIRHNVEDGSVQLADHYQQNT-
PIGDGPVLLPRNHYLSTQSALSKDPKEKRDHMVLLEFVTAAGITHGMDERYK

*GFP(+12):* (mutations to GFP(+6))
MGHHHHHHGGASKGERLFTGVVPILVELDGDVNGHKFSVRGEG**K**GDATNGKLTLKFICTTGKLPVP-
WPTLVTTLTYGVQCFSRYPKHMKQHDFFKSAMPEGYVQERTISFKKDG**K**YKTRAEVKFEGRTLVN-
RIELKGRDFKEDGNILGHKL**R**YNFNSH**K**VYITADKQKNGIKANFKIRHNVEDGSVQLADHYQQNT-
PIGDGPVLLPRNHYLSTQSALSKDPKEKRDHMVLLEFVTAAGITHGMDERYK

*GFP(+18):* (mutations to GFP(+12))
MGHHHHHHGGASKGERLFTGVVPILVEL**K**GDVNGHKFSVRG**K**GKGDATNGKLTLKFICTTGKLPVP-
WPTLVTTLTYGVQCFSRYPKHMKQHDFFKSAMPEGYVQERTISFKKDGKYKTRAEVKFEGRTLVNRI-
ELKGRDFKE**K**GNILGHKLRYNFNSHKVYITADKQKNGIKANFKIRHNVEDGSVQLADHYQQNTPIG-
DGPVLLPRNHYLSTQSALSKDPKEKRDHMVLLEFVTAAGITHGMDERYK

*GFP(+24):* (mutations to GFP(+18))
MGHHHHHHGGASKGERLFTG**K**VPILVELKGDVNGHKFSVRGKGKGDATNGKLTLKFICTTGKLPVP-
WPTLVTTLTYGVQCFSRYPKHMKQHDFFKSAMPEGYVQERTISFKKDGKYKTRAEVKFEGRTLVN-
RI**K**LKGRDFKEKGNILGHKLRYNFNSHKVYITADK**R**KNGIKANFKIRHNVEDGSVQLADHYQQNTPI-
G**R**GPVLLPRNHYLSTQSALSKDPKEKRDHMVLLEFVTAAGITHGMDERYK

*GFP(+30):* (mutations to GFP(+24))
MGHHHHHHGGASKGERLFTGKVPILVELKGDVNGHKFSVRGKGKGDATNGKLTLKFICTTGKLPVP-
WPTLVTTLTYGVQCFSRYPKHMKQHDFFKSAMP**K**GYVQERTISFKKDGKYKTRAEVKFEGRTLVN-
RIKLKGRDFKEKGNILGHKLRYNFNSHKVYITADKRKNGIKANFKIRHNV**K**DGSVQLADHYQQNTPI-
GRGPVLLPRNHYLSTQS**K**LSKDPKEKRDHMVLLEFVTAAGITHGRDERYK

*GFP(+36):* (mutations to GFP(+30))
MGHHHHHHGGASKGERLF**R**GKVPILVELKGDVNGHKFSVRGKGKGDAT**R**GKLTLKFICTTGKLPVP-
WPTLVTTLTYGVQCFSRYPKHMK**R**HDFFKSAMPKGYVQERTISFKKDGKYKTRAEVKFEGRTLVN-
RIKLKGRDFKEKGNILGHKLRYNFNSHKVYITADKRKNGIKA**K**FKIRHNVKDGSVQLADHYQQNTPI-
GRGPVLLPRNHYLST**R**SKLSKDPKEKRDHMVLLEFVTAAGI**K**HGRDERYK

*GFP(-30):* (mutations to sfGFP)
MGHHHHHHGGASKGEELF**D**GVVPILVELDGDVNGH**E**FSVRGEGEGDAT**E**G**E**LTLKFICTTG**E**LPVP-
WPTLVTTLTYGVQCFS**D**YPDHM**D**QHDFFKSAMPEGYVQERTISFKDDGTYKTRAEVKFEGDTLVN-
RIELKGIDFKEDGNILGHKLEYNFNSH**D**VYITADKQ**E**NGIKA**E**F**E**IRHNVEDGSVQLADHYQQNTPIG-
DGPVLLPD**D**HYLST**E**SALSKDPNE**D**RDHMVLLEFVTAAGI**D**HGMDELYK

*IB-GFP(+12):* (deletion to GFP(+12) denoted by **Residue 1|Residue 2**)
MGHHHHHHGGASKGERLFTGVVPILVELDGDVNGHKFSVRGEGKGDATNGKLTLKFICTTGKLPVP-
WPTLVTTLTYGVQCFSRYPKHMKQHDFFKSAMPEGYVQERTISFKKDGKYKTRAEVKFEGRTLVNRI-
ELKGRDFKEDGNILGHKLRYNFNSHKVYITADKQKNGIKANFKIRHNVEDGSVQLADHYQQNTPIG-

**D|R**NHYLSTQSALSKDPKEKRDHMVLLEFVTAAGITHGMDERYK

*IB-GFP(+18):* (deletion to GFP(+18) denoted by **Residue 1|Residue 2**)
MGHHHHHHGGASKGERLFTGVVPILVELKGDVNGHKFSVRGKGKGDATNGKLTLKFICTTGKLPVP-WPTLVTTLTYGVQCFSRYPKHMKQHDFFKSAMPEGYVQERTISFKKDGKYKTRAEVKFEGRTLVNRI-ELKGRDFKEKGNILGHKLRYNFNSHKVYITADKQKNGIKANFKIRHNVEDGSVQLADHYQQNTPIG-**D|R**NHYLSTQSALSKDPKEKRDHMVLLEFVTAAGITHGMDERYK

*IB-GFP(+36):* (deletion to GFP(+36) denoted by **Residue 1|Residue 2**)
MGHHHHHHGGASKGERLFRGKVPILVELKGDVNGHKFSVRGKGKGDATRGKLTLKFICTTGKLPVP-WPTLVTTLTYGVQCFSRYPKHMKRHDFFKSAMPKGYVQERTISFKKDGKYKTRAEVKFEGRTLVN-RIKLKGRDFKEKGNILGHKLRYNFNSHKVYITADKRKNGIKAKFKIRHNVKDGSVQLADHYQQNTPI-G**R|R**NHYLSTRSKLSKDPKEKRDHMVLLEFVTAAGIKHGRDERYK

Protein expression in *E. coli*
Cells were streaked from a glycerol stock onto an agar plate supplemented with 100 µg/mL ampicillin (Gold Biotechnology) and grown at 37 ºC overnight. A single colony was then inoculated into sterilized LB media supplemented with 100 µg/mL ampicillin and grown to saturation in an incubator (Thermo Scientific MaxQ 6000) at 37 ºC overnight with shaking at 250 rpm. The overnight culture was inoculated into 1 L of LB media supplemented with 100 µg/mL ampicillin and allowed to grow at 37 ºC to $OD_{600}$ between 0.8-1.0. At this point, the cultures were induced with 1 mL of 1 M isopropyl ß-D-1-thiogalacopyranoside (IPTG, Gold Biotechnology). Induction temperatures were optimized for protein expression as follows: sfGFP, GFP(0), GFP(+6), GFP(+12) were incubated at 37 ºC after induction and GFP(-30), GFP(+18), GFP(+24), GFP(+30), GFP(+36), and IB-GFP(+36) were incubated at 25 ºC post-induction. Cells were grown for an additional 16-18 h after induction. NiCo21(DE3) cells (New England Biolabs) were used for protein expression.

Protein purification
Following protein expression, cells were harvested by centrifugation (Thermo Scientific Sorvall Legend XTR) in a swinging bucket rotor (Thermo Scientific TX-750) at 4000 rpm for 15 min and resuspended in lysis buffer (50 mM $NaH_2PO_4$, 300 mM NaCl, pH 8.0). Purification of GFP(+18), GFP(+24), GFP(+30) and GFP(+36) was optimized by increasing the NaCl concentration of all buffers to 1 M. The cell pellet was then subjected to one freeze thaw cycle and subsequently lysed by sonication (cycle: 2s on, 4s off) at 60% amplitude using a 0.5 in probe for 10 min (Fisher Scientific). Soluble proteins were separated from cell debris by centrifugation in a fixed angle rotor (Thermo Scientific, Fiberlite F15-8x50cy) at 10,000 rpm for 30 min at room temperature. Proteins in the soluble fraction were collected and purified using immobilized metal affinity chromatography. Briefly, 8-10 mL of His-Pur Ni-NTA resin (Thermo Scientific) was used per L of cell culture. The resin was initially equilibrated in cell lysis buffer before incubation with the soluble cell lysate. Flow throughs, wash fractions (in lysis buffer containing 50 mM imidazole) and elution fractions (in lysis buffer containing 250 mM imidazole) were collected. Fractions were analyzed on a Bolt™ 4-12% Bis-Tris Plus gel (Invitrogen) to determine protein purity, and pure fractions were concentrated using Amicon Ultra centrifugal filter units with 10 kDa molecular weight cutoffs (Millipore Sigma). Proteins were then dialyzed against a buffer that mimics the physiological ion concentration in *E. coli* (physiological buffer: 70 mM $K_2HPO_4$, 60 mM KCl, 40 mM NaCl, pH 7.4) for at least 3 h per buffer change and a total of 7 buffer changes.

SDS-PAGE analysis
Samples were prepared in 4X LDS loading dye and incubated at room temperature for 30-60 min. Bolt™ 4-12% Bis-Tris Plus gels (Invitrogen) were loaded with 15 µL of each sample and were run at 200 V for 30 min. Gels were then stained and destained following the SimplyBlue SafeStain protocol.

Turbidimetry assay

Stock solutions (1 mg/mL) of GFP and total RNA from torula yeast type VI (Sigma-Aldrich) were prepared in physiological buffer (70 mM $K_2HPO_4$, 60 mM KCl, 40 mM NaCl, pH 7.4). GFP and RNA stock solutions were filtered using a 0.22 µm SFCA Thermo Scientific Nalgene 25 mm Syringe Filter and mixed at GFP mass fractions in increments of 0.04 varying from 0 to 1. All samples were prepared in triplicate on a tissue culture-treated polystyrene 96-well half-area plate (Corning, REF#3697), and the absorbance of each sample was measured at 600 nm on a plate reader (Tecan Infinite M200 Pro). The absorbance (A) was converted to turbidity using the following equation: $Turbidity = 100 - 10^{2-A}$.

Construction of phase diagrams for GFP variants

GFP variants and RNA were mixed at various mass ratios using 2 mg/mL or 0.1 mg/mL stock solutions prepared in physiological buffer. All solutions were filtered using 0.22 µm SFCA Thermo Scientific Nalgene 25 mm Syringe Filters. Protein concentrations were obtained by measuring absorbance at 488 nm on a Cary 60 UV-Vis spectrophotometer (Agilent Technologies) and calculating concentrations using Beer's Law. The extinction coefficient for sfGFP ($\varepsilon = 8.33 \times 10^5$ $M^{-1}$ $cm^{-1}$) was used to calculate the concentrations for all supercharged GFP variants. All samples were prepared in triplicate to a total of 50 µL on a 384 well glass-bottom plate (Cellvis) and the absorbance at 600 nm was measured immediately after mixing on a plate reader (Tecan Infinite M200 Pro) at 25 °C. The samples were then imaged as described below. Heatmaps of the turbidity values at each of the tested macromolecule concentrations were generated in Python and transferred to Adobe Illustrator.

Microscopy of coacervates

Mixtures were prepared in a 384-well glass-bottom plate (Cellvis) and imaged at room temperature (approximately 25 ºC) immediately after turbidimetry assays. Images were acquired using a 20X 0.40 NA Plan Fluor objective on GFP ($\lambda_{ex}$= 470-522 nm; $\lambda_{em}$= 525-550 nm) and brightfield channels on an EVOS FL Auto 2 inverted fluorescence microscope (Invitrogen).

Protein characterization by MALDI-TOF mass spectrometry

A buffer exchange was performed on protein stock solutions into milliQ water using Amicon Ultra centrifugal filter units with a 10 kDa molecular weight cutoff (Millipore Sigma). Samples were then prepared using a 10 mg/mL sinapinic acid matrix (7:3 water:acetonitrile with 0.1% trifluoroacetic acid) and were either premixed at a 4:6 sample:matrix ratio or directly spotted on a grounded stainless steel target at a 3:7 sample:matrix ratio. Samples were calibrated with Protein Standard II (Bruker) and MALDI-TOF spectra were obtained from the Columbia University Chemistry Department Mass Spectrometry Facility on a Bruker ultrafleXtreme MALDI TOF/TOF.

Sample preparation for *E. coli* microscopy

Agarose pads were prepared by pipetting 50 µL of 1 w/v% agarose (TopVision) in milliQ water on a clean 1 mm thick, 25x75 mm microscope slide. A coverslip (No. 1.5 18x18 mm, Thermo Scientific) was immediately placed on top. Pads were allowed to solidify for 10-60 min. The cover slip was gently removed, and 1.5 µL cell culture was pipetted onto the agarose pad, secured with a coverslip, and sealed on all four sides with nail polish.

Microscopy of *E. coli* cells

A single colony was selected from a plate and inoculated into 5 mL LB media supplemented with 100 µg/mL ampicillin and grown to saturation overnight at 37 ºC, with shaking at 250 rpm. The saturated culture was then diluted in 50 mL LB media supplemented with ampicillin in a 125 mL Erlenmeyer flask to an $OD_{600}$ of 0.1 and then divided into 2 x 25 mL cultures in sterile 125 mL Erlenmeyer flasks to provide an uninduced control culture. The cultures were grown to an $OD_{600}$ of 0.8-1.0, and then one culture was induced with 1 mM IPTG. Both induced and uninduced cultures were incubated at 25 °C, with shaking at 225 rpm. An image of the overnight cultures and diluted cultures ($OD_{600} \sim 0.1$) was taken to ensure that cultures were not contaminated. Microscopy samples were prepared as described above. Cells were imaged using a 100X oil 1.40 NA UPlanSApo objective (Olympus) on GFP ($\lambda_{ex}$= 470-522 nm; $\lambda_{em}$= 525-550 nm; EVOS GFP light cube) and brightfield channels using an EVOS

FL Auto 2 inverted fluorescence microscope. Cells were then imaged on agarose pads at 0, 2, 8, and 18 or 24 h post-induction. 10-15 images were acquired for each strain in order to image at least 120 cells per strain.

Image analysis

Microscopy images were analyzed using MicrobeJ (version 15.3l (14)), an ImageJ plugin. MicrobeJ was used to threshold the microscopy images, identify cells, and measure intracellular fluorescence intensity. Cells were then sorted in order of increasing length, and demographs and normalized fluorescence intensity profiles of all the GFP variants were generated. Images were thresholded using the Li method (-30) and constraints were placed on cell area (≥1.43 $\mu m^2$), length (≥1.43 µm), width (0-1.5 µm), circularity (0.25-1), and angularity (0-0.4 rad). For GFP(+24) and GFP(+36) 24 h post-induction, thresholding was adjusted to -10 slightly to improve the ability to identify cells.

Cell growth and protein expression assay

The cellular growth of NiCo21(DE3) *E. coli* and fluorescent protein production was monitored for the GFP variants studied here (sfGFP, GFP(0), GFP(+6), GFP(+12), GFP(+18), GFP(+24), GFP(+30), and GFP(+36)) using a plate based growth assay. Three colonies containing the plasmid for each variant were selected from a freshly streaked plate and grown to saturation overnight at 37 °C. The optical density ($OD_{600}$) was measured and each replicate was back-diluted to an $OD_{600}$ of 0.1 in a 96-well deep-well plate (1.5 mL total volume). After 4.75 h the cultures had reached a corrected $OD_{600}$ between 0.8-1.0. At this point, the cultures were induced with 1 mM IPTG. After additional growth with shaking at 25 °C for 2 h, 8 h, 18 h, and 24 h, 25 µL of culture was transferred to a half-area flat bottom clear plate for measurement of GFP fluorescence ($\lambda_{ex}$= 488 ± 9 nm, $\lambda_{ex}$= 530 ± 20 nm) and $OD_{600}$ on an Infinite M200 Pro microplate reader (Tecan). The data were analyzed by dividing the background-corrected GFP fluorescence values by the background-corrected $OD_{600}$ values.

Solubility assay

1 L cell cultures were grown and induced as described above (Protein expression in *E. coli*). 270 mL of the cultures were harvested and resuspended in 4.5 mL cell lysis buffer supplemented with a protease inhibitor cocktail (as directed by Sigma-Aldrich, P8849-1ML), and subjected to one freeze thaw cycle. Afterwards, cultures were sonicated using a microtip (cycle: 2 s on, 4 s off) at 40% amplitude for 5 min and pelleted by centrifugation at 10,000 rpm for 30 min. The pellet containing cell debris and protein condensates was then washed by resuspending in 4.5 mL cell lysis buffer (50 mM $NaH_2PO_4$, 300 mM NaCl, pH 8.0), divided into 3 x 1.5 mL aliquots and pelleted again by centrifugation at 10,000 rpm for 30 min. To solubilize the protein condensates, pellets were resuspended in 1.4 mL cell lysis buffer (control) or lysis buffer containing either 1 M NaCl or 8 M urea. The resuspended lysates were pelleted again to separate the solubilized protein components from the insoluble cell debris. All supernatants (soluble fractions) were collected and pellets (insoluble fractions) were resuspended in their respective buffers before analysis by SDS-PAGE to examine condensate solubility. These samples were run on Bolt™ 4-12% Bis-Tris Plus gels (Invitrogen) and imaged using a Gel Doc XR+ System (Bio-Rad). Gel images were inverted and background subtracted (rolling ball, 100 pixels) in Fiji prior to analysis. Rectangular ROIs were drawn for each lane to generate histograms and line segments were manually drawn at the base of peaks to define bands of interest. The area under the curve (AUC) was measured for all peaks in Fiji and used to calculate the GFP fraction for each sample. The fraction of GFP extracted in Figure 3A was obtained by normalizing the GFP fraction (ratio of GFP band intensity to the total protein band intensity in a given lane) for each GFP variant.

Extraction of streptavidin inclusion bodies

Similar to the solubility assay protocol, cell cultures expressing streptavidin were grown and induced as described above. Briefly, cultures were harvested, frozen and then thawed, sonicated, pelleted, and resuspended in 1.5 mL cell lysis buffer, 1 M NaCl lysis buffer, or 8 M urea lysis buffer (per 90 mL cell culture). After pelleting the resuspended lysate, soluble and insoluble fractions were collected and analyzed by SDS-PAGE.

Photobleaching and Recovery (FRAP)

A single colony was inoculated from a plate into 5 mL LB media supplemented with 100 µg/mL ampicillin and grown to saturation overnight at 37 °C, with shaking at 250 rpm. The saturated culture was then diluted 200-fold in 25 mL LB media supplemented with ampicillin in a sterile 125 mL erlenmeyer flask, grown to an $OD_{600}$ of 0.8-1, and then induced with 1 mM IPTG for 16-18 h at 25 °C with shaking at 225 rpm. Samples were then prepared on agarose pads as described above (Sample preparation for *E. coli* microscopy) and images were acquired with a 100X oil 1.45 NA objective on EGFP ($\lambda_{ex}$= 488 nm; $\lambda_{em}$= 500-550 nm) and brightfield (TD) channels using a Nikon A1RMP multiphoton confocal microscope located in the Confocal and Specialized Microscopy Shared Resource of the Herbert Irving Comprehensive Cancer Center at Columbia University (HICCC). At the beginning of each FRAP experiment, 5 frames were collected pre-bleach at 265.9 ms/frame. A laser spot 0.2 µm in diameter was positioned at one pole of the cell. Cells were bleached in 1 frame at 234.9 ms/frame using a 405 nm laser at 75% power and 486 nm laser at 100% power. 60 frames were collected post-bleach at 1.06 sec/frame for all GFP variants. All post-bleach frames were captured using a 486 nm laser at 0.5% power. Recovery curves were fit to a single exponential recovery function using easyFRAP and averaged over 5-9 cells (*44*). Replicates with a gap ratio and/or bleach depth below 0.6 were excluded from analysis because of excessive or insufficient bleaching.

Nucleic acid staining of cells

Cells were grown and GFP expression was induced as described above (Microscopy of *E. coli* cells). At 18 h post-induction, 250 µL aliquots of cell culture were washed and resuspended in an equal volume of cell lysis buffer (50 mM $NaH_2PO_4$, 300 mM NaCl, pH 8.0) by centrifugation at 13,000 rpm for 1 min. Either 0.5 µL of a 5 mg/mL DAPI stock (Invitrogen) in DMF or 0.5 µL of a 0.5 mM SYTO17 stock (Invitrogen) in DMSO were added to the resuspended cell culture and allowed to incubate in a dark cabinet at room temperature for 10 min. Cell imaging was performed as described above. In addition to GFP and brightfield channels, cells were also imaged on either DAPI ($\lambda_{ex}$= 357-444 nm; $\lambda_{em}$= 447-560 nm; EVOS DAPI light cube) or Texas Red channels ($\lambda_{ex}$= 585-629 nm; $\lambda_{em}$= 624-640 nm; Invitrogen Texas Red light cube).

Colocalization analysis

Microscopy images were pre-processed using rolling ball subtraction (100 pixels) and manually adjusted for optimal channel alignment in Fiji. Line-cuts were drawn down the medial axis of representative cells and fluorescence intensity was measured along the line-cut in GFP, DAPI, and/or Texas Red channels. Fluorescence intensities were then normalized for each channel and plotted as a function of distance. Analysis of images obtained from proteomics validation assays detailed below were also subjected to the same analysis procedure to generate normalized fluorescence intensity plots from the GFP and Texas Red channels.

Nucleic acid *in vitro* staining

The affinity of DAPI and SYTO17 for RNA and DNA were investigated. Briefly, 1 mg/mL stock solutions of RNA from torula yeast type VI (Sigma-Aldrich) and DNA sodium salt from *Oncerhynchus keta* (Sigma-Aldrich) were prepared by dissolution in physiological buffer (70 mM $K_2HPO_4$, 60 mM KCl, 40 mM NaCl, pH 7.4) and filtered using 0.22 µm SFCA Thermo Scientific Nalgene 25 mm Syringe Filters. 1 µL of a 0.5 mg/mL DAPI stock solution in DMF or 1 µL of a 0.05 mM SYTO17 stock solution in DMSO were added to 50 µL nucleic acid solutions in a 384 well glass bottom plate (Cellvis) and allowed to incubate in a dark drawer at room temperature for 10 min before imaging at 20X magnification on GFP, DAPI, Texas Red, and brightfield channels as described above. Fluorescence intensity measurements were obtained using Fiji. Background subtracted fluorescence intensity values were plotted.

Proteomics Sample Preparation

Cells were streaked from a glycerol stock onto an LB plate supplemented with 100 µg/mL ampicillin. The plate was incubated at 37 °C overnight and was then stored at 4 °C for at most 2 weeks. A single colony was then inoculated into 6 mL LB media supplemented with 100 µg/mL ampicillin and grown to saturation at 37 ºC overnight with shaking at 225 rpm. 5 mL of overnight culture was inoculated into 1 L of LB media supplemented with 100 μg/mL ampicillin and allowed to grow at 37 ºC to $OD_{600}$ between 0.8-1.0. At this point, the cultures were induced with 1 mL of 1 M IPTG and incubated at 25 ºC for an additional 16-18 h post-induction.

Cell harvesting: Following protein expression, 900 mL cells were harvested by centrifugation at 4000 rpm for 10 min at 4 °C, washed twice with 450 mL of cold lysis buffer (50 mM $NaH_2PO_4$, 300 mM NaCl, dissolved in MilliQ water, pH 8.0, stored at 4 °C) and resuspended in lysis buffer for a 0.5 g wet cell weight/mL cell lysate. 300 μL of protease inhibitor cocktail (Sigma-Aldrich, P8849-1ML) was added to 15 mL cell lysate. The cell lysate was then frozen in liquid nitrogen and stored at -80 °C.

Isolation of condensate components: Cells were then thawed in an ice water bath and lysed by sonication (60% amplitude; 2 s on, 4 s off; 8 min sonics) in an ice water bath supplemented with $CaCl_2$ (10 g $CaCl_2$ / 500 g ice). Soluble proteins were separated from cell debris by centrifugation at 10,000 rpm for 30 min at 4 °C in a fixed angle rotor. The pellet was washed in 15 mL lysis buffer (50 mM $NaH_2PO_4$, 300 mM NaCl prepared with optima grade water, pH 8) and centrifuged again at 10,000 rpm for 30 min at 4 °C. The pellets were then resuspended in either lysis buffer (50 mM $NaH_2PO_4$, 300 mM NaCl prepared with optima grade water, pH 8) or 1 M NaCl lysis buffer (50 mM $NaH_2PO_4$, 1 M NaCl prepared with optima grade water, pH 8) and pelleted again by centrifugation at the same speed, time, and temperature. The supernatants were collected in 1.5 mL Eppendorf tubes and spun again in a microcentrifuge tube at 13,000 rpm for 30 min at 4 °C to pellet residual cell debris. The top portion of supernatant was collected for further proteomics analysis, leaving ~100 μL near the undisturbed pellet.

Removal of GFP by Ni-NTA chromatography: 6xHis-tagged GFP in the collected fractions were then depleted from the samples using Nickel Affinity Chromatography. Briefly, 1.25 mL of Ni-NTA resin was used per 3-4 mL of soluble protein. The resin was initially equilibrated in 5 mL (~10 column volumes) of cell lysis buffer supplemented with 10 mM imidazole (50 mM $NaH_2PO_4$, 300 mM NaCl, 10 mM imidazole prepared with optima grade water, pH 8). The sample was then incubated with Ni-NTA resin for 10 min at 4 °C. The flow through, wash fractions (50 mM $NaH_2PO_4$, 1 M NaCl, 35 mM imidazole prepared with optima grade water, pH 8) and elution fractions (50 mM $NaH_2PO_4$, 1 M NaCl, 250 mM imidazole prepared with optima grade water, pH 8) were collected in 4 mL fractions. For each sample, 1 x 4 mL flow through fraction (F1), 1 x 4 mL rinse in lysis buffer (F2), 3 x 4 mL wash fractions (W), and 3 x 4 mL elution fractions (E) were collected. Fractions were analyzed by SDS-PAGE on Bolt 4-12% Bis-Tris Plus gels as described above (SDS-PAGE Analysis).

Protein extraction/precipitation: Proteins were extracted from the flow through (F1) and rinse (F2) fractions. For each sample, 1 mL of flow through and rinse fractions were mixed in a 15 mL Falcon tube and 3 mL Optima grade water (Fisher Scientific) was added to decrease the overall salt concentration to ~250 mM NaCl. The diluted sample was then mixed with 5 mL methanol and 1.25 mL chloroform by vortexing to form a milky emulsion. Samples were then centrifuged at 4,000 rpm for 20 min at 4 °C in a swinging bucket rotor to facilitate phase separation. The top layer was removed with a pipette without disturbing the protein layer at the interface. 5 mL methanol was then added and samples were mixed gently by inversion. The protein samples were then pelleted by centrifuging at 4,000 rpm for 15 min at 4 °C. The supernatant was removed and the pellets were washed twice with 1 mL ice cold methanol and pelleted by centrifugation at 4,000 rpm for 10 min at 4 °C. The supernatant was removed by pipetting and the pellet was air dried for 10 min at room temperature. Pellets were then resuspended in 30-40 μL solubilization buffer (100 mM ammonium bicarbonate, 8 M urea, 0.1 M dithiothreitol, prepared daily) and transferred to a 1.5 mL Eppendorf microcentrifuge tube. A Bradford assay (Bio-Rad) was performed to determine the protein concentration of each sample. Samples were then snap frozen in liquid nitrogen and stored at -80 °C until further processing.

Sample processing, mass spectrometry, and data analysis: All solvents and acids were Optima LC/MS Grade (Fisher Scientific). Samples were further reduced with DTT at a final total concentration of 5.5 mM at room temperature for 40 min. Proteins were alkylated with iodoacetamide at a concentration of 15 mM in the dark at room temperature for 30 min. Prior to digestion, samples were diluted five-fold in 100 mM ammonium bicarbonate and digested using sequencing grade trypsin (6 ng/μl, Promega Corp. V511) overnight at 37 °C. Following digestion, samples were acidified to pH<2 with TFA, kept on ice for 15 min to precipitate lipids and centrifuged for 15 min at 3,000 rpm. Samples were desalted using Nestgroup C18 Macrospin columns, lyophilized, and redissolved in 2% acetonitrile, 0.1% formic and subjected to LC/MS/MS analysis. A Q Exactive HF (Orbitrap) mass spectrometer (Thermo Scientific) was operated in positive ion mode using data-dependent acquisition. Peptides were loaded on a 75 µm ID x 2 cm Acclaim PepMap C18 trap column prior to separation at 5 µL/min for 3 min. Separation was performed with a 75 µm ID x 50 cm Acclaim PepMap C18 column on an Ultimate 3000 RSLCnano liquid chromatograph. Flow rate was 300 nL/min with an acetonitrile/formic acid gradient. Mobile Phase A was 0.1% formic acid in water and mobile phase B was 20% water, 80% acetonitrile with 0.1% formic acid. Two chromatograms were recorded for each of six biological replicates for a total of 12 runs. All mass spectrometry raw files generated in this work have been publicly deposited as described in the data and materials availability.

The results were analyzed with Rosetta Elucidator Protein Expression Analysis System (V. 4.0.0.2.31 Ceiba Solutions/ PerkinElmer) and aligned based on accurate mass and retention time of detected features using the PeakTeller algorithm in Elucidator. Label-free quantitation was based on comparison of MS1 feature volume of aligned features across all runs. Data were searched against *Escherichia coli* (strain B / BL21-DE3) UniProt database (release-2018_10 from 11/7/18 with 8,398 sequences and 2,628,066 residues) with isoforms. This database also had appended common laboratory contaminants and four recombinant protein sequences: GFP(+36), SlyD::CBD, Can::CBD, GlmS6Ala. Searches were conducted on an in-house Mascot server (V. 2.5.1, Matrix Science Ltd., London, UK). The three fusion proteins (along with Arn::CBD) were engineered into commercial strains by the manufacturer for protein purification purposes (New England Biolabs). Search parameters included fixed modification on Cys (carbamidomethyl) and mass accuracy limits of 10 ppm for MS and 0.02 Da for MS/MS. Variable modifications were Oxidation (Met), Gln>pyro-Glu (N-term Q) and Acetyl (Protein N-term). Identified features were annotated at 1.0% false discovery rate (FDR) using the Peptide Teller algorithm in Elucidator. Overall, 239,789 features were detected by Elucidator with identifications returned for 1,412 proteins. Of the 1,412 proteins, one added protein (trypsin), five laboratory contaminants (keratin) and 337 proteins represented by a single peptide were excluded from the analysis. The remaining 1,069 proteins represented by two or more peptides were included in the analysis. All proteomics results are summarized in Supplementary Data S1.

Analysis of *E. coli* proteome

All protein sequences for *E. coli* strain BL21-DE3 (UP000002032) were retrieved from UniProt. We calculated the expected net charge for each protein at a pH of 7.4 using the following set of ionizable residues (with indicated average $pK_a$ values): N-terminus (9.42), C-terminus (2.18), arginine (12.48), lysine (10.53), histidine (6.00), aspartic acid (3.65), glutamic acid (4.25), and tyrosine (10.07). The Henderson-Hasselbach equation was then used to calculate the predicted net charge for each protein. This data was then used to generate a histogram using R Studio (Figure 1B). This process was similar to a previously reported protocol (*29*). The protein sequence data was also analyzed for other biochemical characteristics (e.g. ratio of negative-to-positive residues, % of residues that are aromatic) for comparison of proteins identified in the proteomics experiment.

Cloning fusion proteins for proteomics validation

*rraB* and *ecnAB* genes were cloned by colony PCR using Phusion high-fidelity DNA polymerase (NEB). A genomic DNA extraction was performed to clone genes that could not be amplified by colony PCR (see below). To generate arabinose-inducible mScarlet-I fusions, parts were assembled stepwise by HiFi assembly of multiple fragments:

- Gene: genes were amplified from the *E. coli* genome (BL21). Primers for gene amplification were designed according to sequences obtained from KEGG (Kyoto Encyclopedia of Genes and Genomes).
- mScarlet-I: pEB2-mScarlet-I (Addgene)
- Arabinose inducible vector: p2-39 was a gift from the Dickinson lab (*45*).

| Gene fusion | DNA template | Primer | Primer Sequence (5’--> 3’) |
|---|---|---|---|
| mScarlet-I | pEB2-mScarlet-I | mScarlet-I_fwd | aatgaccgctATGAGTAAAGGAGAAGCTGTG |
| | | mScarlet-I_rev | ttaattaagtTTATTTGTATAGTTCATCCATGCC |
| | p2-39 | vector_fwd | ctatacaaataaACTTAATTAACGGCACTCCTC |
| | | vector_rev | ctttactcatAGCGGTCATTTATGTCAGAC |
| rraA- mscarlet-I | BL21 *E. coli* colony | rraA_fwd | aatgaccgctATGAAATACGATACTTCCG |
| | | rraA_rev | ctcctttactaccgcttccacccgacgtTTCAATATCCAGC-GGATC |
| | pEB2-mScarlet-I | mScarlet-I_fwd | acgtcgggtggaagcggtAGTAAAGGAGAAGCTGTG |
| | | mScarlet-I_rev | ttaattaagtTTATTTGTATAGTTCATCCATGC |
| | p2-39 | vector_fwd | ctatacaaataaACTTAATTAACGGCACTC |
| | | vector_rev | cgtatttcatAGCGGTCATTTATGTCAG |
| rnr- mScarlet-I | BL21 *E. coli* colony | rnr_fwd | aatgaccgctATGTCACAAGATCCTTTC |
| | | rnr_rev | ctcctttactaccgcttccacccgacgtCTCTGC-CACTTTTTTCTTC |
| | pEB2-mScarlet-I | mScarlet-I_fwd | acgtcgggtggaagcggtAGTAAAGGAGAAGCTGTG |
| | | mScarlet-I_rev | ttaattaagtTTATTTGTATAGTTCATCCATGC |
| | p2-39 | vector_fwd | ctatacaaataaACTTAATTAACGGCACTC |
| | | vector_rev | cttgtgacatAGCGGTCATTTATGTCAG |
| hldD- mScarlet-I | BL21 *E. coli* colony | hldD_fwd | aatgaccgctATGATCATCGTTACCGGCGG |
| | | hldD_rev | ctcctttactaccgcttccacccgacgtTGCGTCGCGAT-TCAGCCA |
| | pEB2-mScarlet-I | mScarlet-I_fwd | acgtcgggtggaagcggtAGTAAAGGAGAAGCTGTG |
| | | mScarlet-I_rev | ttaattaagtTTATTTGTATAGTTCATCCATGC |
| | p2-39 | vector_fwd | ctatacaaataaACTTAATTAACGGCACTC |
| | | vector_rev | cgatgatcatAGCGGTCATTTATGTCAG |
| speA- mScarlet-I | BL21 *E. coli* colony | speA_fwd | aatgaccgctATGTCTGACGACATGTCTATG |
| | | speA_rev | ctcctttactaccgcttccacccgacgtCTCATCTTCAAGA-TAAGTATAACC |
| | pEB2-mScarlet-I | mScarlet-I_fwd | acgtcgggtggaagcggtAGTAAAGGAGAAGCTGTG |
| | | mScarlet-I_rev | ttaattaagtTTATTTGTATAGTTCATCCATGC |
| | p2-39 | vector_fwd | ctatacaaataaACTTAATTAACGGCACTC |
| | | vector_rev | cgtcagacatAGCGGTCATTTATGTCAG |

| | | | |
|---|---|---|---|
| rraB- mScarlet-I | BL21 *E. coli* colony | rraB_fwd | aatgaccgctATGGCAAACCCGGAACAAC |
| | | rraB_rev | accgcttccacccgacgtGTGGCGAACCCCGTCATC |
| | pEB2-mScarlet-I | mScarlet-I_fwd | ggttcgccacacgtcgggtggaagcggtAGTAAAGGAGAAGCTGTG |
| | | mScarlet-I_rev | ttaattaagtTTATTTGTATAGTTCATCCATGC |
| | p2-39 | vector_fwd | ctatacaaataaACTTAATTAACGGCACTC |
| | | vector_rev | ggtttgccatAGCGGTCATTTATGTCAG |
| ecnAB-mScarlet-I | BL21 *E. coli* colony | ecnAB_fwd | aatgaccgctATGGTGAAGAAGACAATTGCAG |
| | | ecnAB_rev | accgcttccacccgacgtTTGCTGCGCTTTCGTTGC |
| | pEB2-mScarlet-I | mScarle-tI_fwd | agcgcagcaaacgtcgggtggaagcggtAGTAAAGGAGAAGCTGTG |
| | | mScarlet-I_rev | ttaattaagtTTATTTGTATAGTTCATCCATGC |
| | p2-39 | vector_fwd | ctatacaaataaACTTAATTAACGGCACTC |
| | | vector_rev | tcttcaccatAGCGGTCATTTATGTCAG |

The araBAD promoter in p2-39 was then replaced with the araBAD promoter from mOrange2-pBAD (Addgene). Sequence confirmation is available upon request.

| **Gene fusion** | **DNA template** | **Primer** | **Primer Sequence (5’--> 3’)** |
|---|---|---|---|
| mScarlet-I | mOrange2-pBAD | insert pBAD promoter_fwd | cttcagccatACTTTTCATACTCCCGCC |
| | | insert pBAD promoter_rev | AGAACCCCGCATATGTATATC |
| | p2-39 mScarlet-I | p2-39 backbone_F | atatacatatgcggggttctATGAGTAAAGGAGAAGCTGTG |
| | | p2-39 backbone_R | tatgaaaagtATGGCTGAAGCGCAAAATG |
| rraA- mScarlet-I | mOrange2-pBAD | insert pBAD promoter_fwd | cttcagccatACTTTTCATACTCCCGCC |
| | | insert pBAD promoter_rev | AGAACCCCGCATATGTATATC |
| | p2-39 rraA-mScarlet-I | rraA p2-39 backbone_fwd | atatacatatgcggggttctATGAAATACGATACTTCCGAGC |
| | | rraA p2-39 backbone_rev | tatgaaaagtATGGCTGAAGCGCAAAATG |

| | | | |
|---|---|---|---|
| rnr-mScar-let-I | mOrange2-pBAD | insert pBAD promoter_fwd | cttcagccatACTTTTCATACTCCCGCC |
| | | insert pBAD promoter_rev | AGAACCCCGCATATGTATATC |
| | p2-39 rnr-mScar-let-I | rnr p2-39 back-bone_fwd | atatacatatgcggggttctATGTCACAAGATCCTTTC-CAG |
| | | rnr p2-39 back-bone_rev | tatgaaaagtATGGCTGAAGCGCAAAATG |
| hldD- mS-carlet-I | mOrange2-pBAD | insert pBAD promoter_fwd | cttcagccatACTTTTCATACTCCCGCC |
| | | insert pBAD promoter_rev | AGAACCCCGCATATGTATATC |
| | p2-39 hldD-mS-carlet-I | hldD p2-39 backbone_fwd | atatacatatgcggggttctATGATCATCGTTACCGGC |
| | | hldD p2-39 backbone_rev | tatgaaaagtATGGCTGAAGCGCAAAATG |
| speA- mS-carlet-I | mOrange2-pBAD | insert pBAD promoter_fwd | cttcagccatACTTTTCATACTCCCGCC |
| | | insert pBAD promoter_rev | cgtcagacatAGAACCCCGCATATGTATATC |
| | p2-39 speA-mS-carlet-I | speA-mScar-let-I_fwd | gcggggttctATGTCTGACGACATGTCTATG |
| | | speA p2-39 backbone_rev | tatgaaaagtATGGCTGAAGCGCAAAATG |
| rraB- mS-carlet-I | mOrange2-pBAD | insert pBAD promoter_fwd | cttcagccatACTTTTCATACTCCCGCC |
| | | insert pBAD promoter_rev | AGAACCCCGCATATGTATATC |
| | p2-39 rraB-mS-carlet-I | rraB p2-39 backbone_fwd | atatacatatgcggggttctATGGCAAACCCGGAA-CAAC |
| | | rraB p2-39 backbone_rev | tatgaaaagtATGGCTGAAGCGCAAAATG |
| ecnAB-mScarlet-I | mOrange2-pBAD | insert pBAD promoter_fwd | cttcagccatACTTTTCATACTCCCGCC |
| | | insert pBAD promoter_rev | AGAACCCCGCATATGTATATC |
| | p2-39 ec-nAB-mS-carlet-I | ecnAB p2-39 backbone_fwd | atatacatatgcggggttctATGGTGAAGAAGA-CAATTGC |
| | | ecnAB p2-39 backbone_rev | tatgaaaagtATGGCTGAAGCGCAAAATG |

Amino acid sequences of all fusion proteins are provided below. mScarlet-I is shown in pink, endogenous *E. coli* protein in grey, and a linker in black.

*hldD-mScarlet-I:*
MIIVTGGAGFIGSNIVKALNDKGITDILVVDNLKDGTKFVNLVDLNIADYMDKEDFLIQIMAGEEFGD-VEAIFHEGACSSTTEWDGKYMMDNNYQYSKELLHYCLEREIPFLYASSAATYGGRTSDFIESREYEK-PLNVYGYSKFLFDEYVRQILPEANSQIVGFRYFNVYGPREGHKGSMASVAFHLNTQLNNGESP-KLFEGSENFKRDFVYVGDVADVNLWFLENGVSGIFNLGTGRAESFQAVADATLAYHKKGQIEYIP-FPDKLKGRYQAFTQADLTNLRAAGYDKPFKTVAEGVTEYMAWLNRDATSGGSGSKGEAVIKEFM-RFKVHMEGSMNGHEFEIEGEGEGRPYEGTQTAKLKVTKGGPLPFSWDILSPQFMYGSRAFIKHPADIP-DYYKQSFPEGFKWERVMNFEDGGAVTVTQDTSLEDGTLIYKVKLRGTNFPPDGPVMQKKTMGWEA-STERLYPEDGVLKGDIKMALRLKDGGRYLADFKTTYKAKKPVQMPGAYNVDRKLDITSHNEDYTV-VEQYERSEGRHSTGGMDELYK

*rraB-mScarlet-I:*
MANPEQLEEQREETRLIIEELLEDGSDPDALYTIEHHLSADDLETLEKAAVEAFKLGYEVTDPEELE-VEDGDIVICCDILSECALNADLIDAQVEQLMTLAEKFDVEYDGWGTYFEDPNGEDGDDEDF-VDEDDDGVRHTSGGSGSKGEAVIKEFMRFKVHMEGSMNGHEFEIEGEGEGRPYEGTQTAKLKVT-KGGPLPFSWDILSPQFMYGSRAFIKHPADIPDYYKQSFPEGFKWERVMNFEDGGAVTVTQDT-SLEDGTLIYKVKLRGTNFPPDGPVMQKKTMGWEASTERLYPEDGVLKGDIKMALRLKDGGRYLAD-FKTTYKAKKPVQMPGAYNVDRKLDITSHNEDYTVVEQYERSEGRHSTGGMDELYK

*speA-mScarlet-I:*
MSDDMSMGLPSSAGEHGVLRSMQEVAMSSQEASKMLRTYNIAWWGNNYYDVNELGHISVCPDP-DVPEARVDLAQLVKTREAQGQRLPALFCFPQILQHRLRSINAAFKRARESYGYNGDYFLVYPIKVN-QHRRVIESLIHSGEPLGLEAGSKAELMAVLAHAGMTRSVIVCNGYKDREYIRLALIGEKMGHKVYL-VIEKMSEIAIVLDEAERLNVVPRLGVRARLASQGSGKWQSSGGEKSKFGLAATQVLQLVETLREA-GRLDSLQLLHFHLGSQMANIRDIATGVRESARFYVELHKLGVNIQCFDVGGGLGVDYEGTRSQS-DCSVNYGLNEYANNIIWAIGDACEENGLPHPTVITESGRAVTAHHTVLVSNIIGVERNEYTVPTA-PAEDAPRALQSMWETWQEMHEPGTRRSLREWLHDSQMDLHDIHIGYSSGIFSLQERAWAEQLYLS-MCHEVQKQLDPQNRAHRPIIDELQERMADKMYVNFSLFQSMPDAWGIDQLFPVLPLEGLDQVPER-RAVLLDITCDSDGAIDHYIDGDGIATTMPMPEYDPENPPMLGFFMVGAYQEILGNMHNLFGDTEA-VDVFVFPDGSVEVELSDEGDTVADMLQYVQLDPKTLLTQFRDQVKKTDLDAELQQQFLEEFEAG-LYGYTYLEDETSGGSGSKGEAVIKEFMRFKVHMEGSMNGHEFEIEGEGEGRPYEGTQTAKLKVT-KGGPLPFSWDILSPQFMYGSRAFIKHPADIPDYYKQSFPEGFKWERVMNFEDGGAVTVTQDT-SLEDGTLIYKVKLRGTNFPPDGPVMQKKTMGWEASTERLYPEDGVLKGDIKMALRLKDGGRYLAD-FKTTYKAKKPVQMPGAYNVDRKLDITSHNEDYTVVEQYERSEGRHSTGGMDELYK

*rraA-mScarlet-I:*
MKYDTSELCDIYQEDVNVVEPLFSNFGGRASFGGQIITVKCFEDNGLLYDLLEQNGRGRVLVVDG-GGSVRRALVDAELARLAVQNEWEGLVIYGAVRQVDDLEELDIGIQAMAAIPVGAAGEGIGESD-VRVNFGGVTFFSGDHLYADNTGIILSEDPLDIETSGGSGSKGEAVIKEFMRFKVHMEGSMNGHEFEI-EGEGEGRPYEGTQTAKLKVTKGGPLPFSWDILSPQFMYGSRAFIKHPADIPDYYKQSFPEGFKW-ERVMNFEDGGAVTVTQDTSLEDGTLIYKVKLRGTNFPPDGPVMQKKTMGWEASTERLYPEDGVLK-GDIKMALRLKDGGRYLADFKTTYKAKKPVQMPGAYNVDRKLDITSHNEDYTVVEQYERSEGRHST-GGMDELYK

*rnr-mScarlet-I:*
MSQDPFQEREAEKYANPIPSREFILEHLTKREKPASRDELAVELHIEGEEQLEGLRRRLRAM-ERDGQLVFTRRQCYALPERLDLVKGTVIGHRDGYGFLRVEGRKDDLYLSSEQMKTCIHGD-QVLAQPLGADRKGRREARIVRVLVPKTSQIVGRYFTEAGVGFVVPDDSRLSFDILIPPDQIMGARMG-FVVVVELTQRPTRRTKAVGKIVEVLGDNMGTGMAVDIALRTHEIPYIWPQAVEQQVAGLKEEVPEE-AKAGRVDLRDLPLVTIDGEDARDFDDAVYCEKKRGGGWRLWVAIADVSYYVRPSTPLDREARN-

RGTSVYFPSQVIPMLPEVLSNGLCSLNPQVDRLCMVCEMTVSSKGRLTGYKFYEAVMSSHARL-TYTKVWHILQGDQDLREQYAPLVKHLEELHNLYKVLDKAREERGGISFESEEAKFIFNAERRIER-IEQTQRNDAHKLIEECMILANISAARFVEKAKEPALFRIHDKPSTEAITSFRSVLAELGLELPGGNK-PEPRDYAELLESVADRPDAEMLQTMLLRSMKQAIYDPENRGHFGLALQSYAHFTSPIRRYPDLT-LHRAIKYLLAKEQGHQGNTTETGGYHYSMEEMLQLGQHCSMAERRADEATRDVADWLKCDFM-LDQVGNVFKGVISSVTGFGFFVRLDDLFIDGLVHVSSLDNDYYRFDQVGQRLMGESSGQTYRLG-DRVEVRVEAVNMDERKIDFSLISSERAPRNVGKTAREKAKKGDAGKKGGKRRQVGKKVNFEPD-SAFRGEKKTKPKAAKKDARKAKKPSAKTQKIAAATKAKRAAKKKVAETSGGSGSKGEAVIKEFM-RFKVHMEGSMNGHEFEIEGEGEGRPYEGTQTAKLKVTKGGPLPFSWDILSPQFMYGSRAFIKHPADIP-DYYKQSFPEGFKWERVMNFEDGGAVTVTQDTSLEDGTLIYKVKLRGTNFPPDGPVMQKKTMGWEA-STERLYPEDGVLKGDIKMALRLKDGGRYLADFKTTYKAKKPVQMPGAYNVDRKLDITSHNEDYTV-VEQYERSEGRHSTGGMDELYK

*ecnAB-mScarlet-I:*

MVKKTIAAIFSVLVLSTVLTACNTTRGVGEDISDGGNAISGAATKAQQTSGGSGSKGEAVIKEFM-RFKVHMEGSMNGHEFEIEGEGEGRPYEGTQTAKLKVTKGGPLPFSWDILSPQFMYGSRAFIKHPADIP-DYYKQSFPEGFKWERVMNFEDGGAVTVTQDTSLEDGTLIYKVKLRGTNFPPDGPVMQKKTMGWEA-STERLYPEDGVLKGDIKMALRLKDGGRYLADFKTTYKAKKPVQMPGAYNVDRKLDITSHNEDYTV-VEQYERSEGRHSTGGMDELYK

Extraction of *E. coli* genomic DNA

To improve the efficiency of amplification of the selected *E. coli* genes (*rraA*, *rnr*, *hldD* and *speA*), genomic DNA was extracted and used for PCR. Briefly, a single colony (NiCo21(DE3)) was placed in 500 µL sterile dd-$H_2O$ in a 1.7 mL eppendorf tube and centrifuged for 1 min at 11,000 rpm. The supernatant was discarded and the pellet was resuspended in an additional 500 µL sterile dd-$H_2O$. The sample was vortexed and then centrifuged for 1 min at 1,300 rpm. The supernatant was discarded and the pellet was resuspended in 100 µL of sterile dd-$H_2O$. The sample was then sonicated using a microtip for 10 min (cycle: 2 s on, 4 s off) with the sample submerged in a -20 °C aluminum bead bath. After sonication the sample was allowed to settle to the bottom of the tube and was then used directly for PCR.

Proteomics validation assays

Preparation of cells for protein expression was performed as described in the section above (Microscopy of *E. coli* cells). Briefly, a saturated overnight culture was used to inoculate 25 mL sterilized LB media in a 125 mL erlenmeyer flask to an $OD_{600}$ = 0.1 with 0.2% L(+)-arabinose (Acros organics). The culture was grown to $OD_{600}$ = 0.8 - 1.0 at 37 ºC, with shaking at 250 rpm and then induced with 1 mM IPTG. Cultures were then incubated at 25 ºC, with shaking at 250 rpm and imaged after 18 h using the protocols described above. Only cells expressing both GFP and mScarletI fusion proteins were considered for analysis. A small percentage of cells expressing speA-mScarletI exhibited low GFP expression and were wider and longer than cells co-expressing sfGFP and speA-mScarletI at similar levels. Moreover, speA-mScarletI was excluded from local regions in these larger cells resulting in irregular spatial distributions. These anomalous cells were excluded from further analysis.

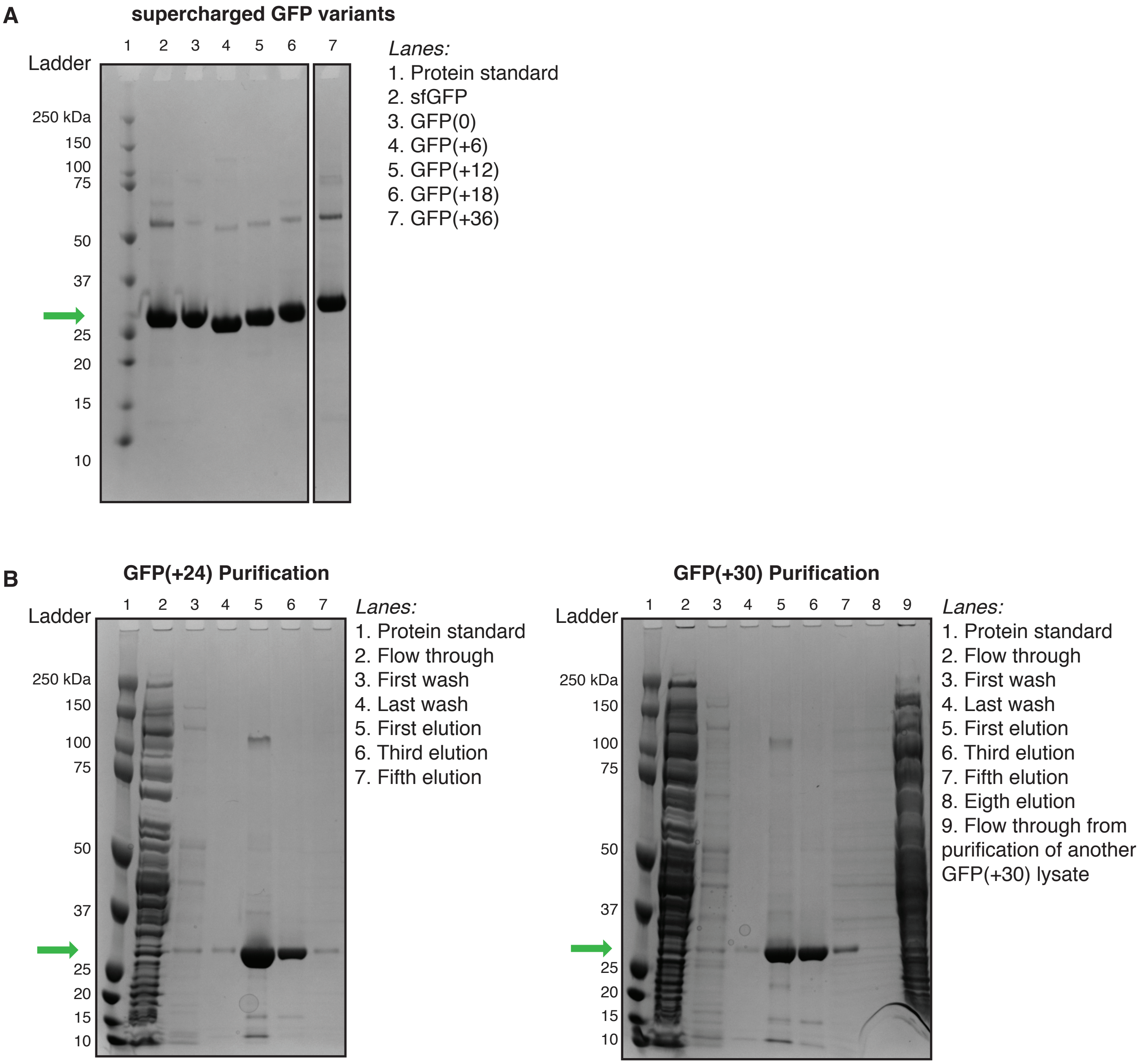


**Figure S1.**

**SDS-PAGE analysis of purified supercharged GFP variants.** (A) Each supercharged variant was purified by affinity chromatography and loaded on a SDS-PAGE gel from a 0.25 mg/mL protein stock. GFP variants were observed to increase in apparent molecular weight with increased supercharging, consistent with their expected molecular weights. All samples were loaded onto the same gel but irrelevant lanes were excluded for clarity. (B) Examples of purified fractions used for turbidimetry assays and construction of phase diagrams. Elutions 1-5 were collected for both GPF(+24) and GFP(+30), dialyzed in physiological buffer, and tested for their ability to phase separate with RNA. Background bands observed in lane 7 on the GFP(+30) SDS-PAGE gel (right) were contamination from lane 9. In all gels, Lane 1 corresponds to precision plus protein dual color standard (Bio-rad). Green arrows indicate bands corresponding to monomeric GFP.

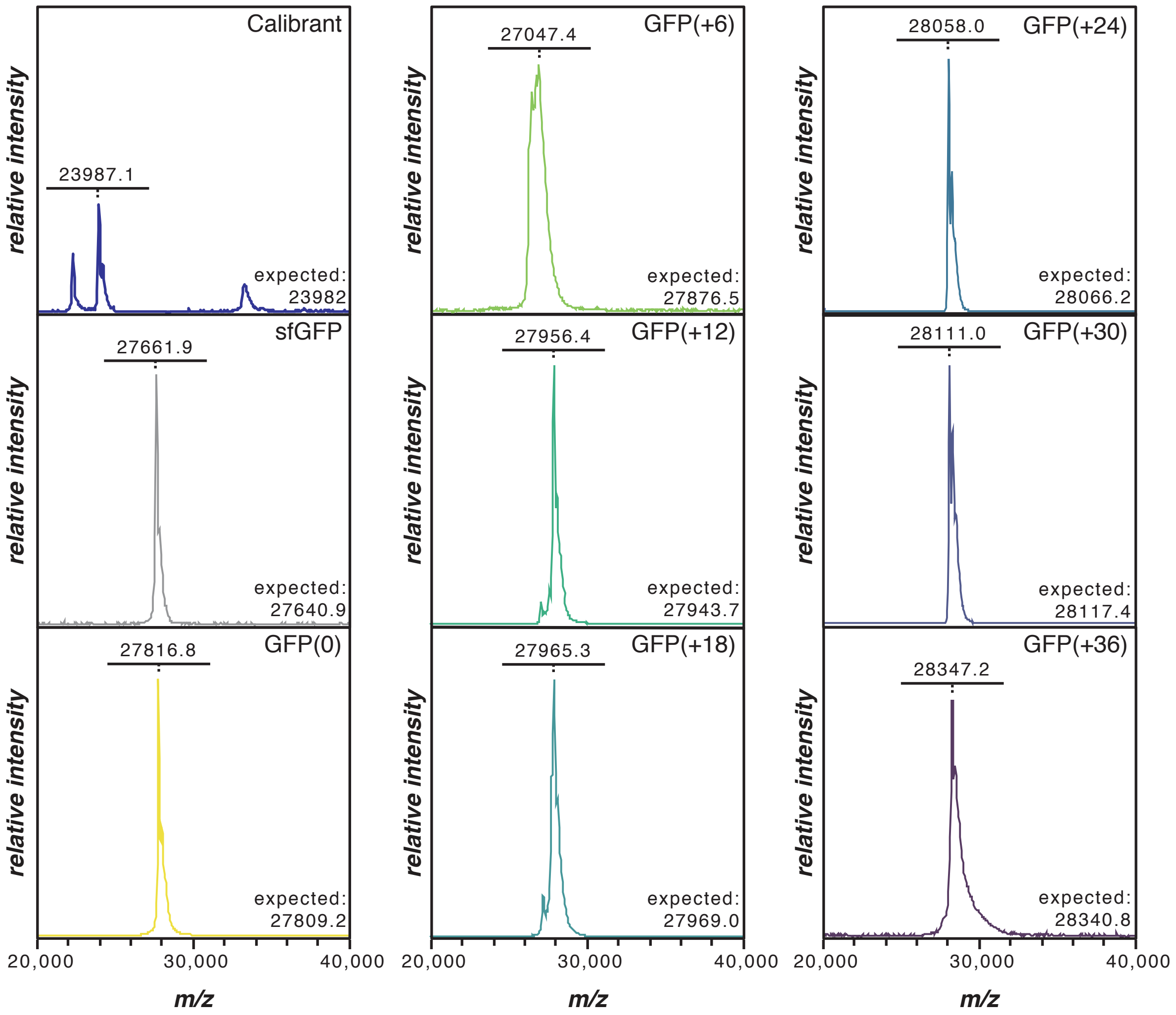


**Figure S2.**

**MALDI-TOF MS of supercharged GFP variants.** Mass spectra for supercharged GFP variants were obtained from MALDI-TOF MS. The spectra were calibrated using an external standard (Bruker Protein Standard II). Observed (underlined) and expected molecular weights are reported for each GFP variant. Expected molecular weights reported here reflect the cleavage of the initial methionine and loss of water due to chromaphore cyclization. For GFP(+6), removal of the N-terminal His tag accounted for the difference between expected and observed molecular weights.

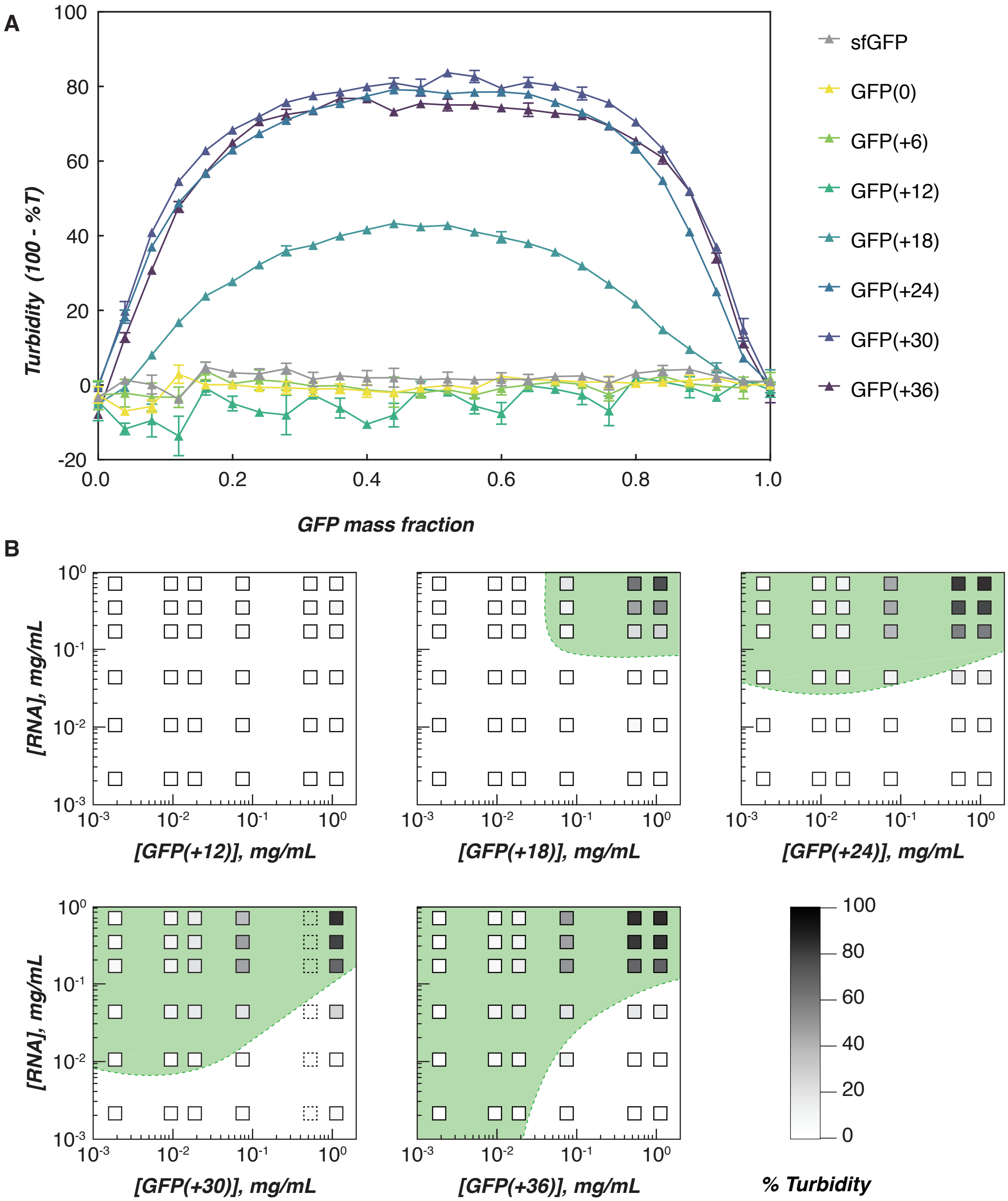


**Figure S3.**

***In vitro* complex coacervation of RNA with supercharged GFPs.** Turbidity assays at (A) fixed macromolecule concentrations and (B) phase diagrams in a physiological buffer that mimics the intracellular ion concentration in E. coli (70 mM $K_2HPO_4$, 60 mM KCl, 40 mM NaCl, pH 7.4). Each supercharged GFP variant was mixed with total RNA from torula yeast at various mixing ratios and at (A) constant 1 mg/mL total macromolecule concentration or (B) varied macromolecule concentrations. Phase separation was detected by turbidimetry measurements at 600 nm and confirmed with optical microscopy. Green dashed lines and green shading represent the observed phase boundaries and two phase regions, respectively. Dashed boxes indicate concentrations for which turbidity measurements were not obtained due to extenuating circumstances caused by COVID-19.

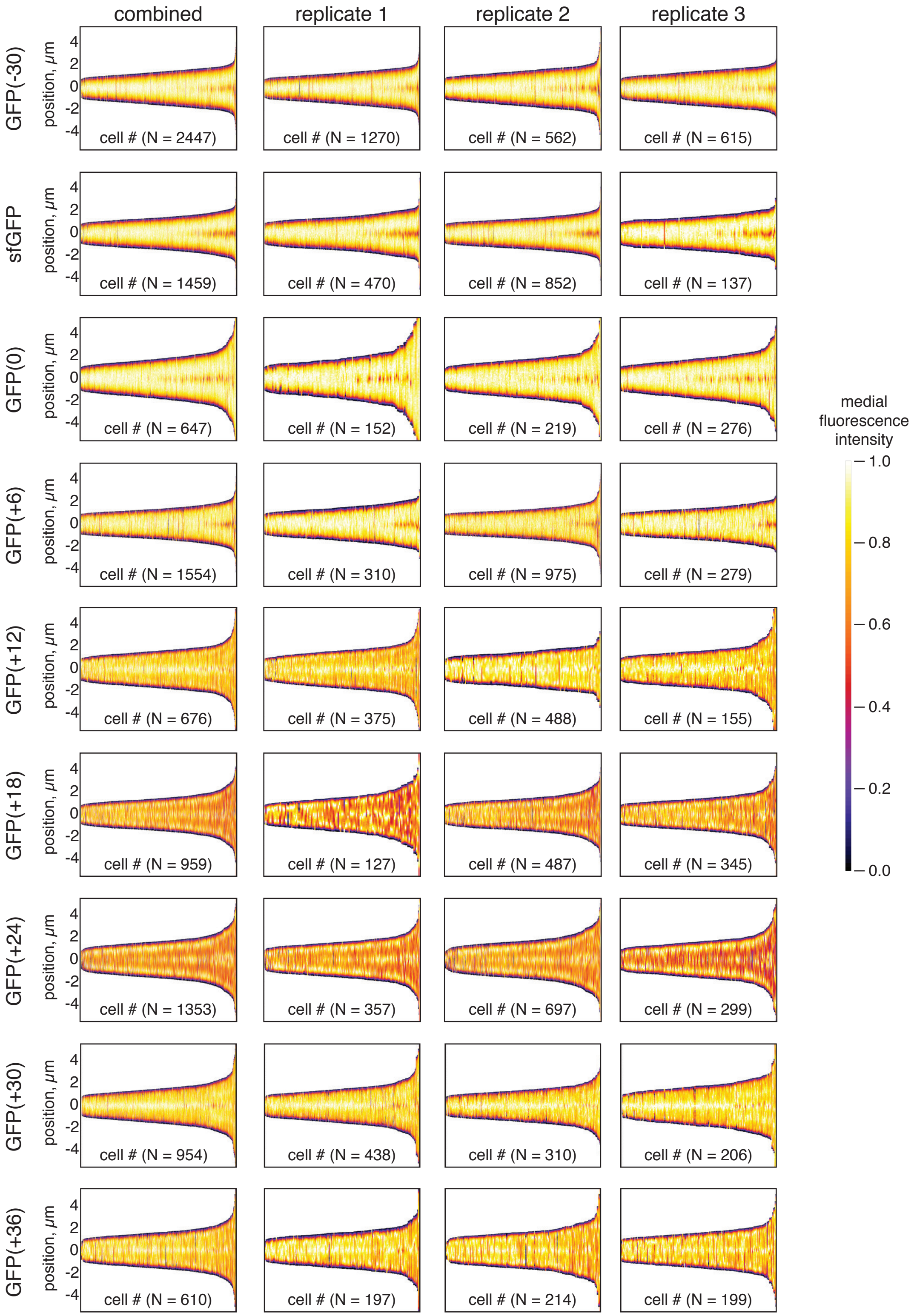


**Figure S4.**

**Spatial GFP distribution of all three biological replicates for each supercharged GFP variant at 2 h post-induction.** Demographs depict GFP localization for N cells. The position, 0 μm, represents the cell center. Images from each replicate were analyzed separately (replicate) or all together (combined) to generate the demographs.

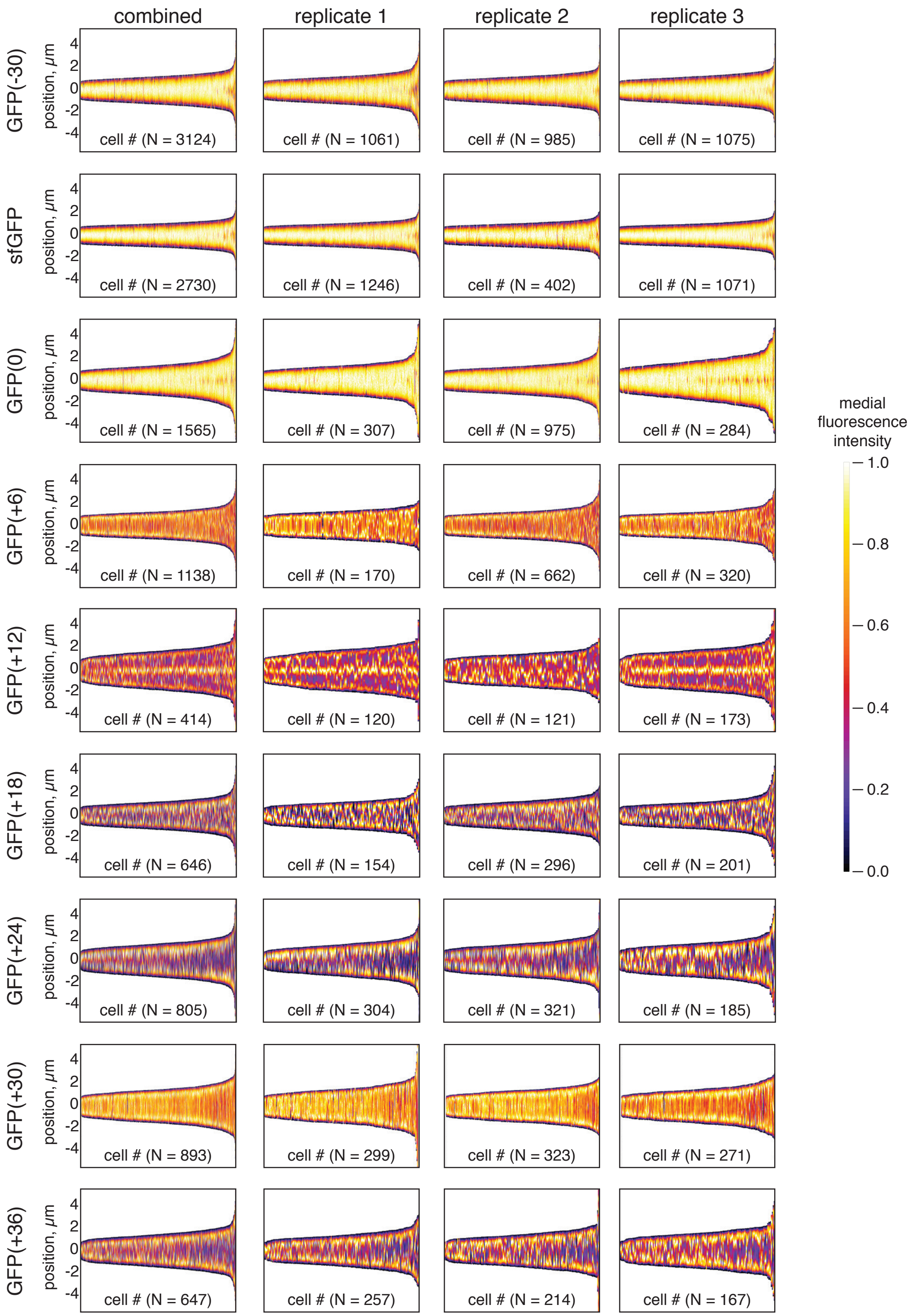


**Figure S5.**

**Spatial GFP distribution of all three biological replicates for each supercharged GFP variant at 8 h post-induction.** Demographs depict GFP localization for N cells. The position, 0 μm, represents the cell center. Images from each replicate were analyzed separately (replicate) or all together (combined) to generate the demographs.

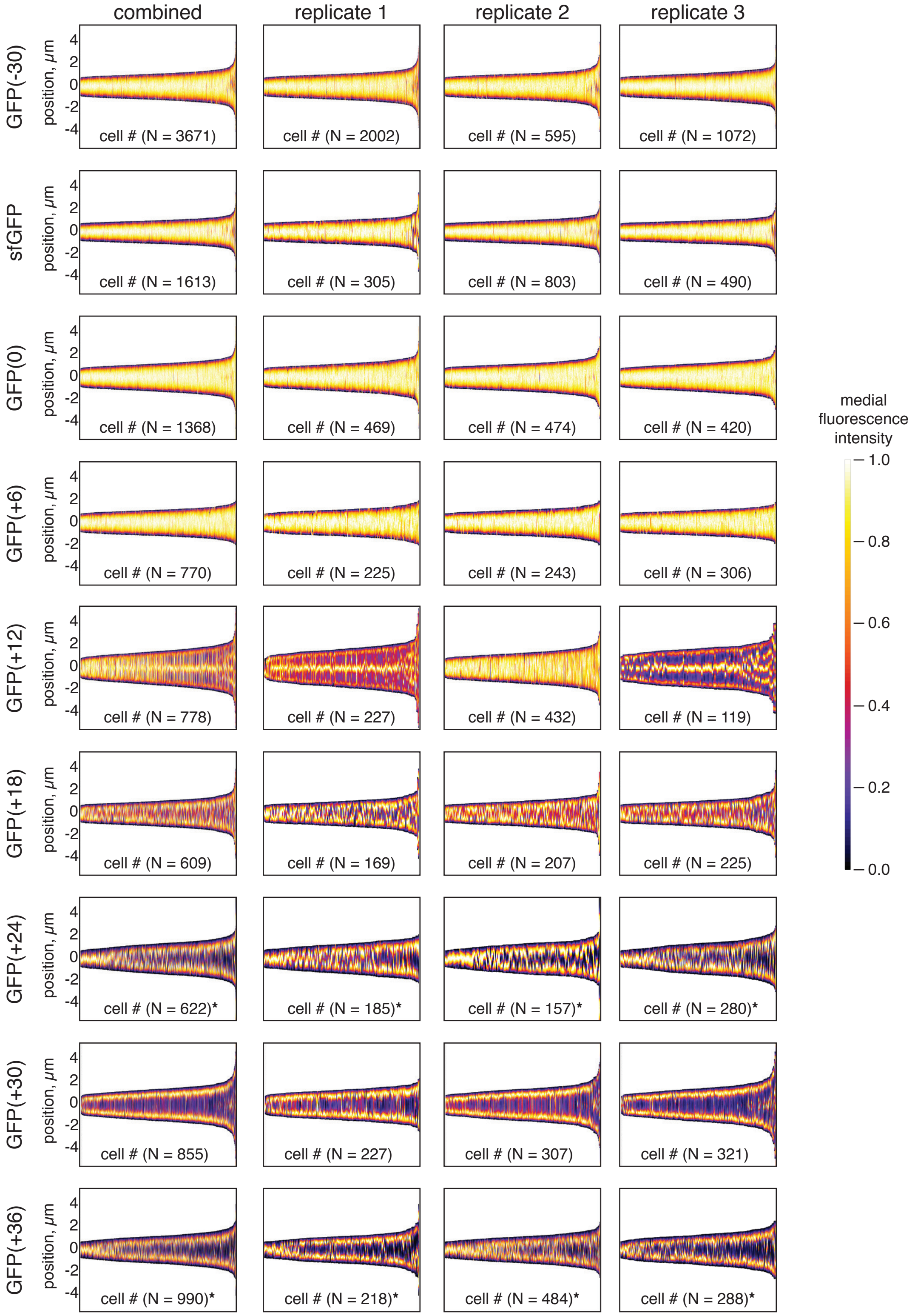


**Figure S6.**

**Spatial GFP distribution of all three biological replicates for each supercharged GFP variant at 24 h post-induction.** Demographs depict GFP localization for N cells. The position, 0 μm, represents the cell center. Images from each replicate were analyzed separately (replicate) or all together (combined) to generate the demographs. *Thresholding settings were adjusted for image analysis of cells expressing GFP(+24) and GFP(+36) at 24 h to improve the identification of cells in microbeJ.

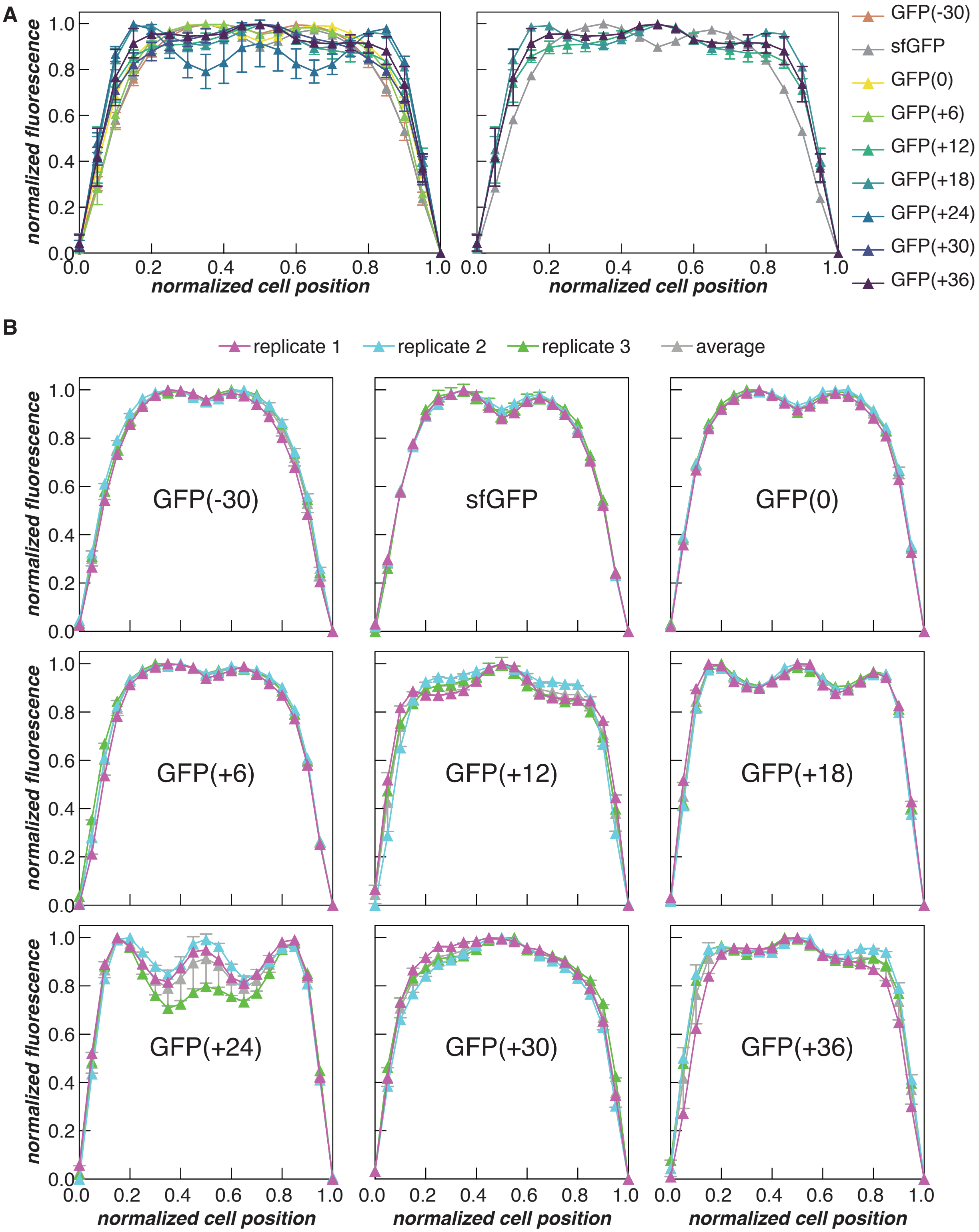


**Figure S7.**

**Normalized fluorescence intensity plots for all three biological replicates at 2 h post-induction.** Cell length and fluorescence intensity were normalized for each GFP variant. (A) Averages and standard deviations of normalized fluorescence intensities are depicted for all GFP variants (left) and variants shown in Figure 2C (right). (B) The average and standard deviation of normalized fluorescence for individual biological replicates is depicted for all GFP variants as well as the average across all replicates (grey).

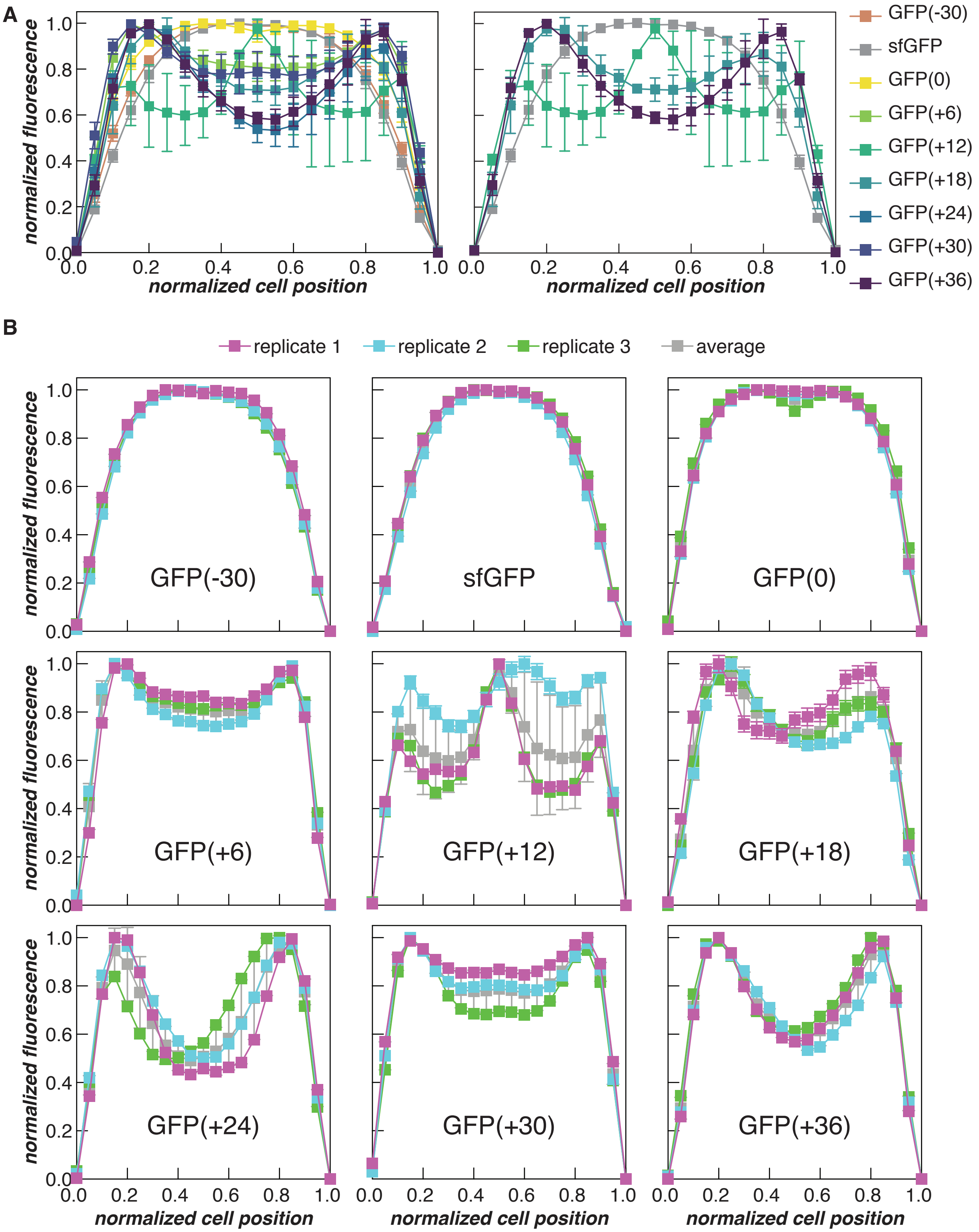


**Figure S8.**

**Normalized fluorescence intensity plots for all three biological replicates at 8 h post-induction.** Cell length and fluorescence intensity were normalized for each GFP variant. (A) Averages and standard deviations of normalized fluorescence intensities are depicted for all GFP variants (left) and variants shown in Figure 2C (right). (B) The average and standard deviation of normalized fluorescence for individual biological replicates is depicted for all GFP variants as well as the average across all replicates (grey).

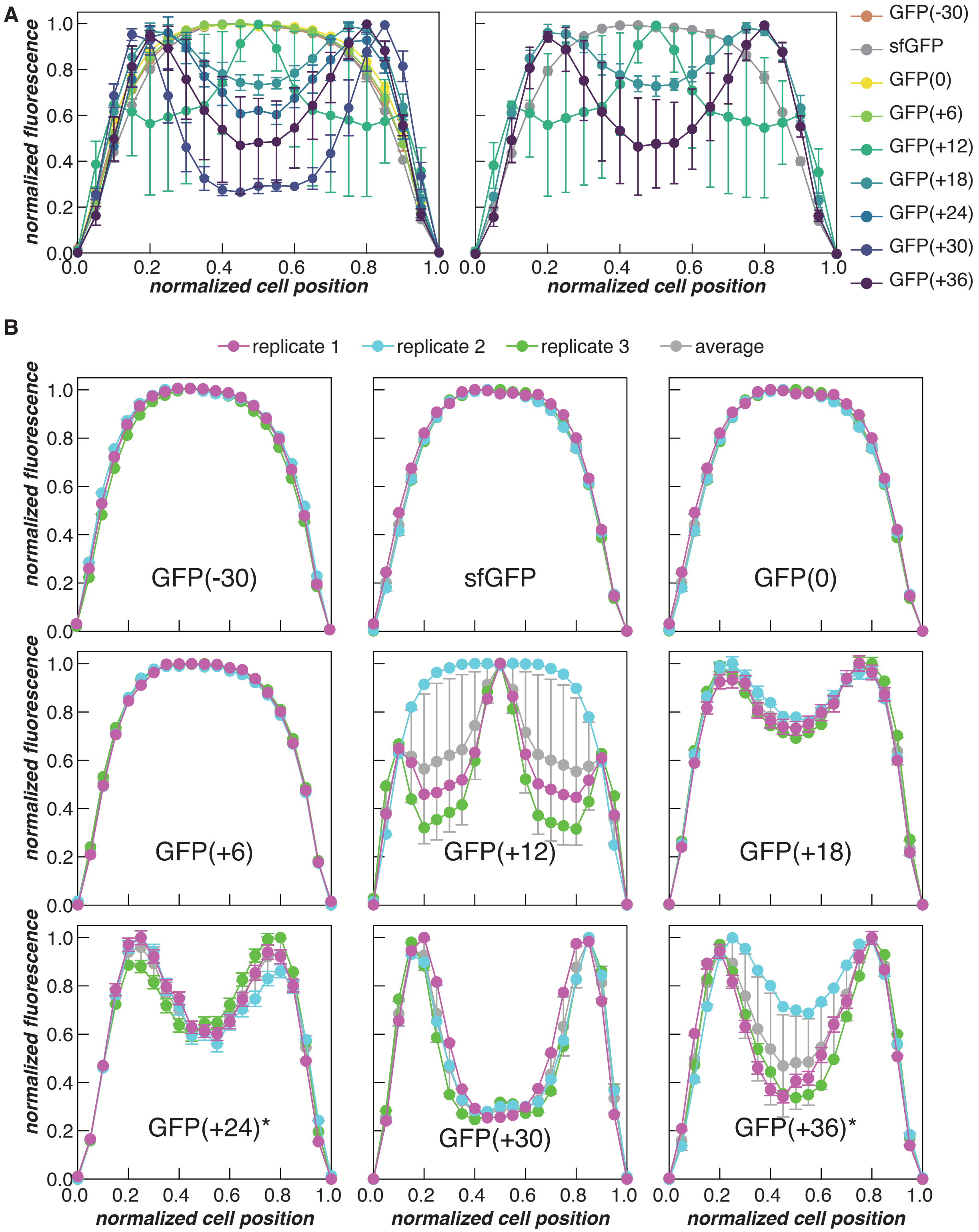


**Figure S9.**

**Normalized fluorescence intensity plots for all three biological replicates at 24 h post-induction.** Cell length and fluorescence intensity were normalized for each GFP variant. (A) Averages and standard deviations of normalized fluorescence intensities are depicted for all GFP variants (left) and variants shown in Figure 2C (right). (B) The average and standard deviation of normalized fluorescence for individual biological replicates is depicted for all GFP variants as well as the average across all replicates (grey).*Thresholding settings were adjusted for image analysis of cells expressing GFP(+24) and GFP(+36) at 24 h to improve the identification of cells in microbeJ.

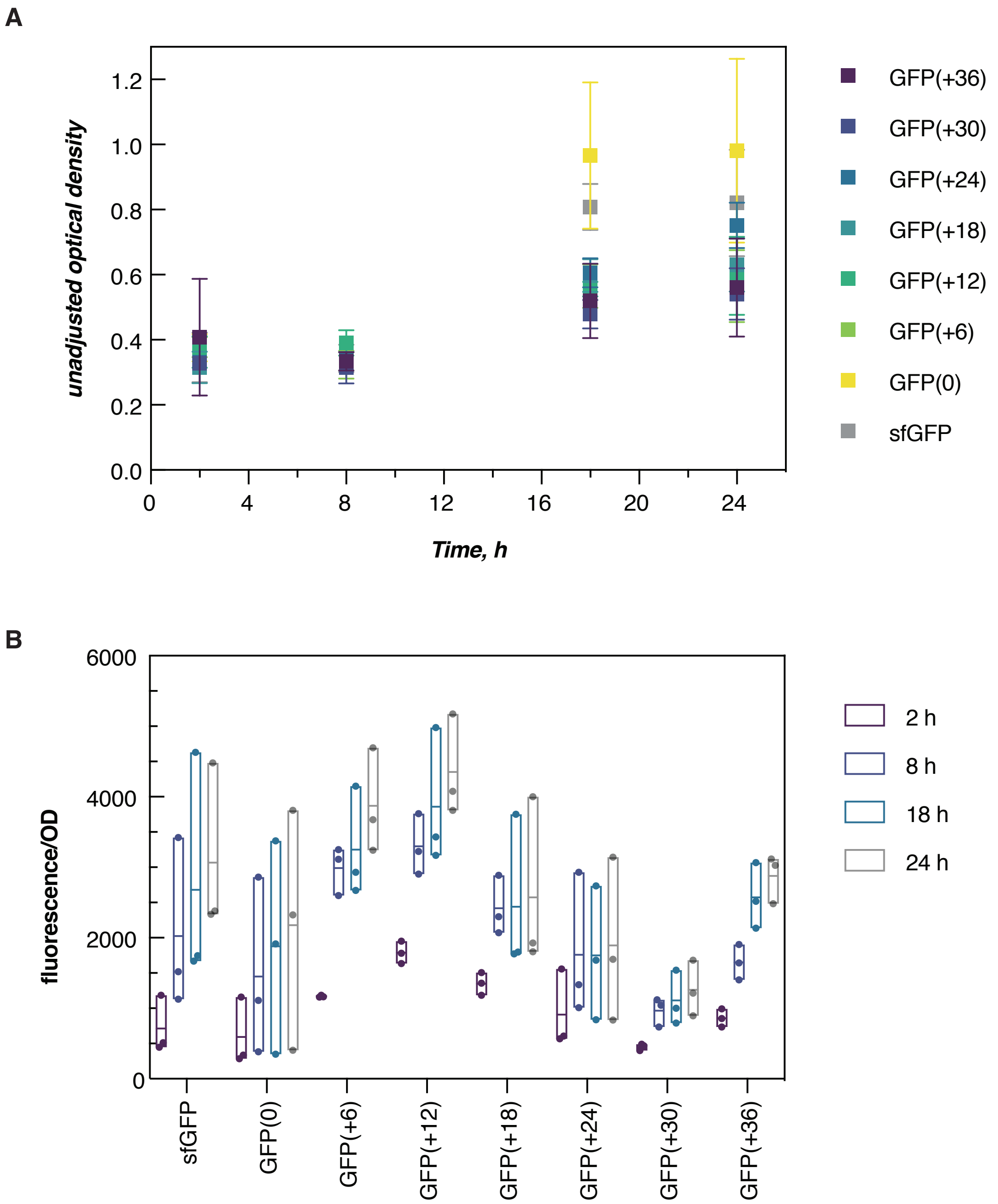


**Figure S10.**

**Cell viability and protein expression.** Optical density measurements at 600 nm and fluorescence intensity ($\lambda_{ex}$ = 488 nm; $\lambda_{em}$ = 530 nm) were obtained for all supercharged variants 2, 8, 18, and 24 h post-induction. (A) The average $OD_{600}$ and standard deviation of all variants grown at 25 °C at 2 h, 8 h, 18 h and 24 h post-induction. All variants demonstrate an increase in optical density after 24 h, with the lower charge variants growing slightly better than the rest. (B) Fluorescence intensity was normalized to $OD_{600}$ and showed an increase in intracellular GFP concentration over 24 h. Individual data points are plotted in addition to the average normalized fluorescence, and range.

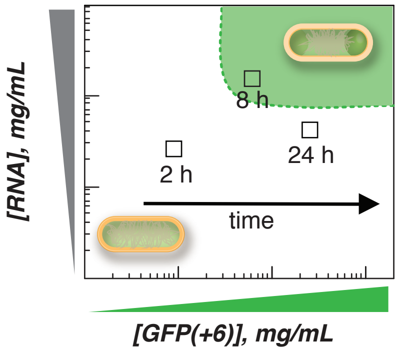


**Figure S11.**

**Schematic of traversing the phase boundary in bacterial cells.** Formation of condensates at 8 h and their disassembly at 24 h is likely due to changes in intracellular protein and RNA concentration over the course of the cell growth cycle.

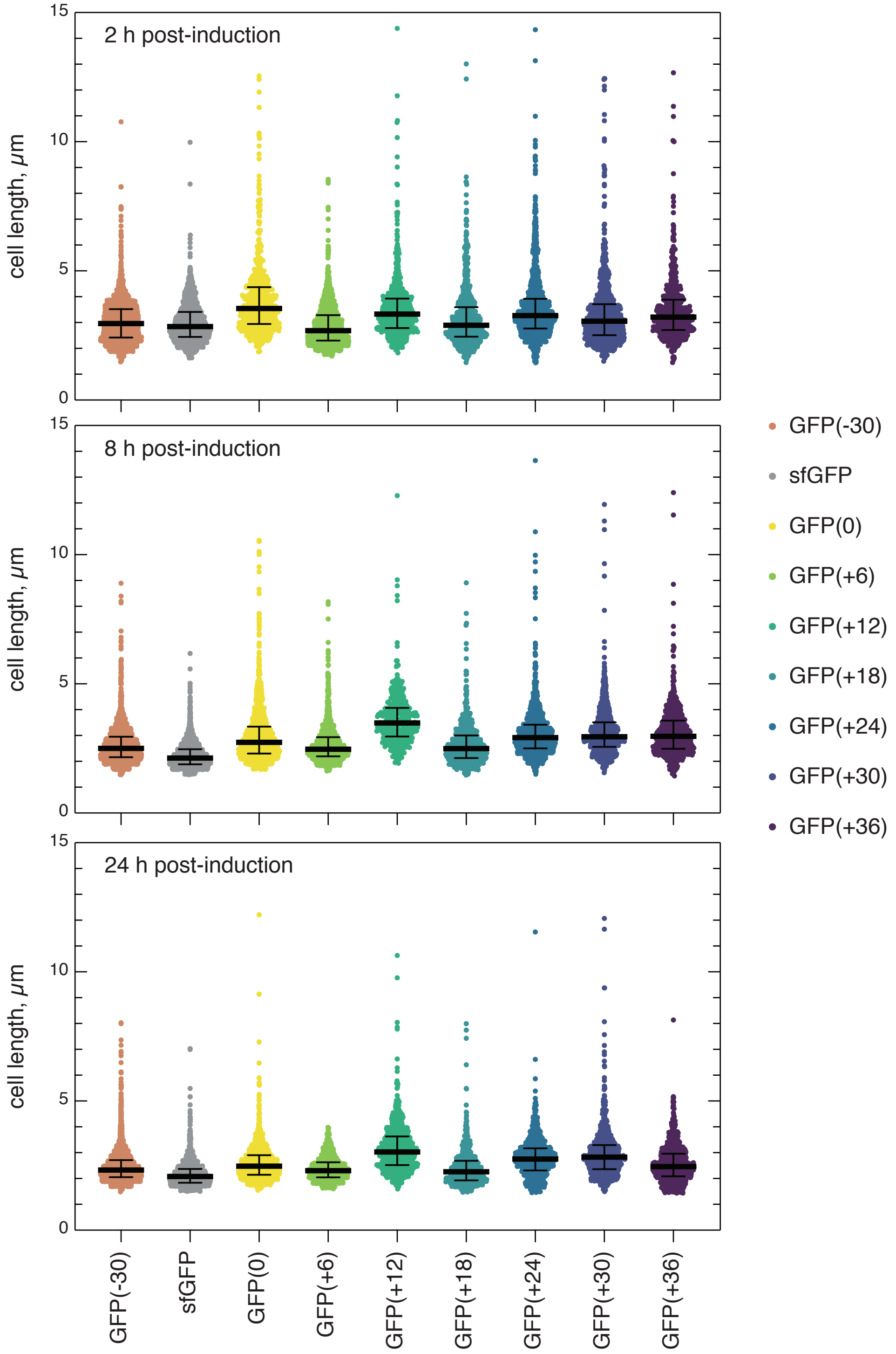


**Figure S12.**

**Cell length distribution of combined replicates at 2, 8, and 24 h.** Cell lengths were obtained from microbeJ and scatterplots were generated using GraphPad. The median and interquartile ranges are depicted for all variants. In general, the length of cells expressing different supercharged GFP variants were comparable. At longer time points, cells expressing GFP(+12) were slightly longer.

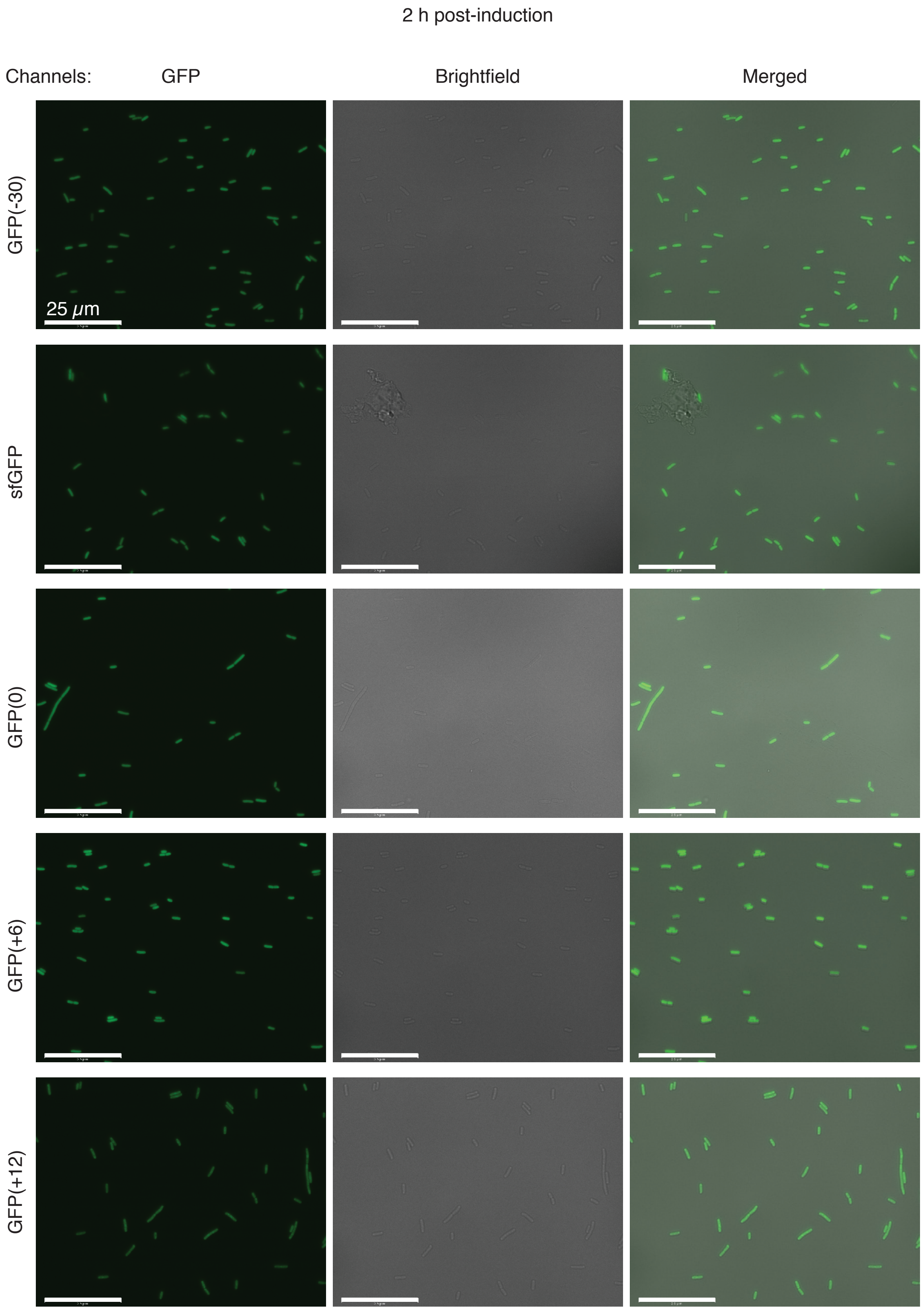
2 h post-induction
Channels:
GFP
Brightfield
Merged
GFP(-30)
sfGFP
GFP(0)
GFP(+6)
GFP(+12)
25 μm

2 h post-induction (continued)

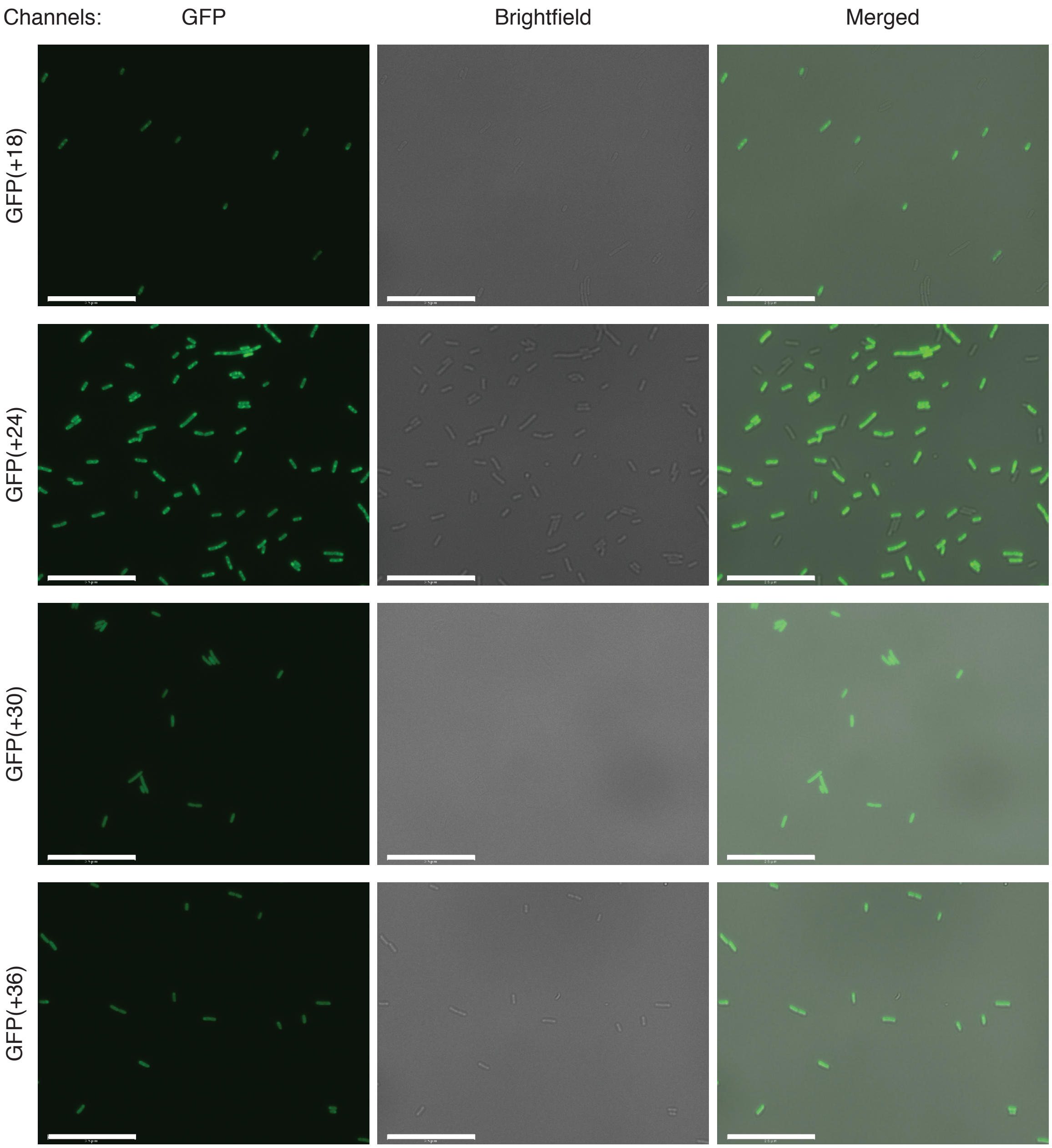


**Figure S13.**

**Representative images acquired from microscopy of cells expressing each supercharged variant at 2 h post-induction.** Representative cell populations are shown on GFP, brightfield, and merged channels. Scale bars are 25 μm.

8 h post-induction

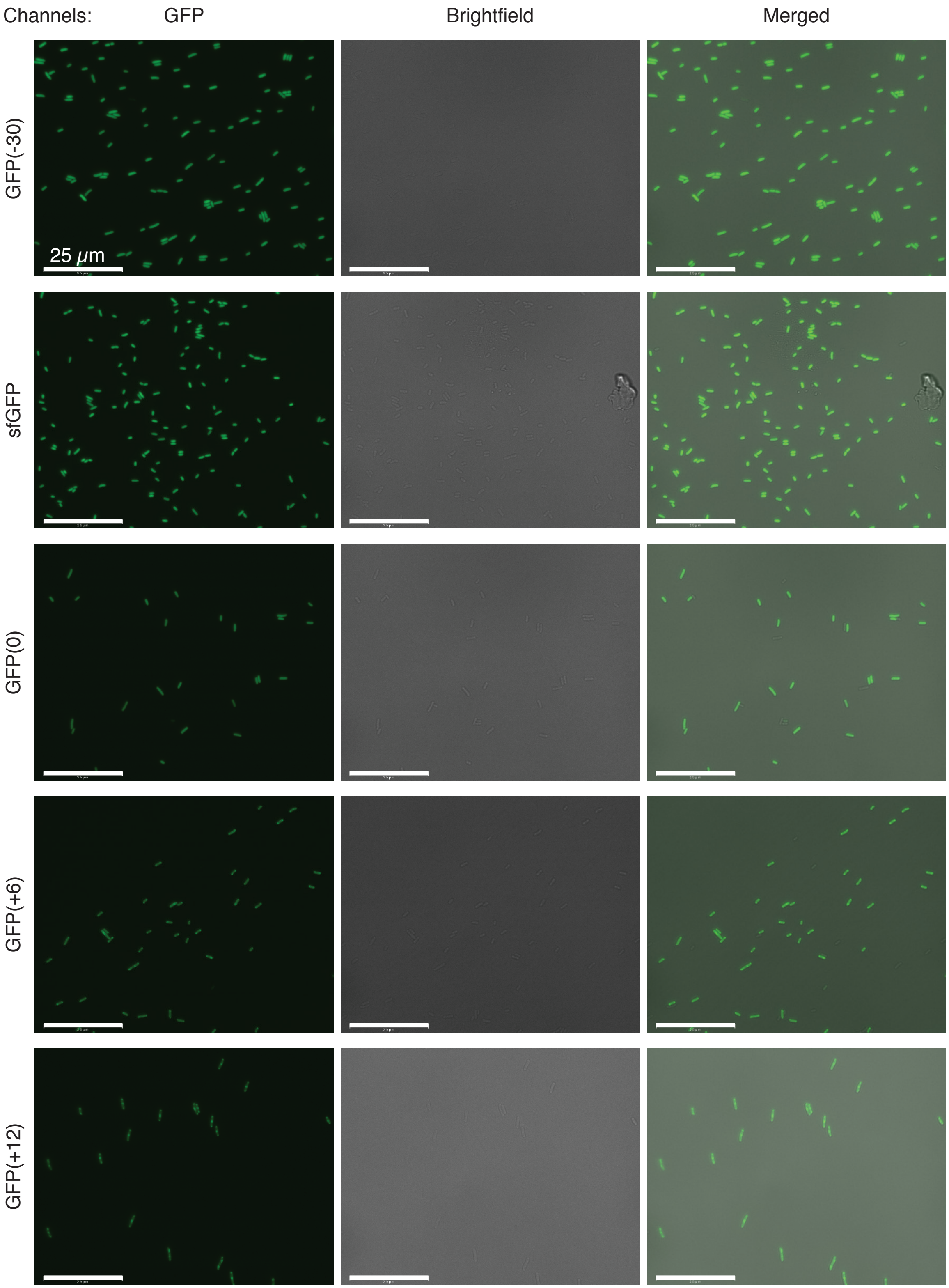

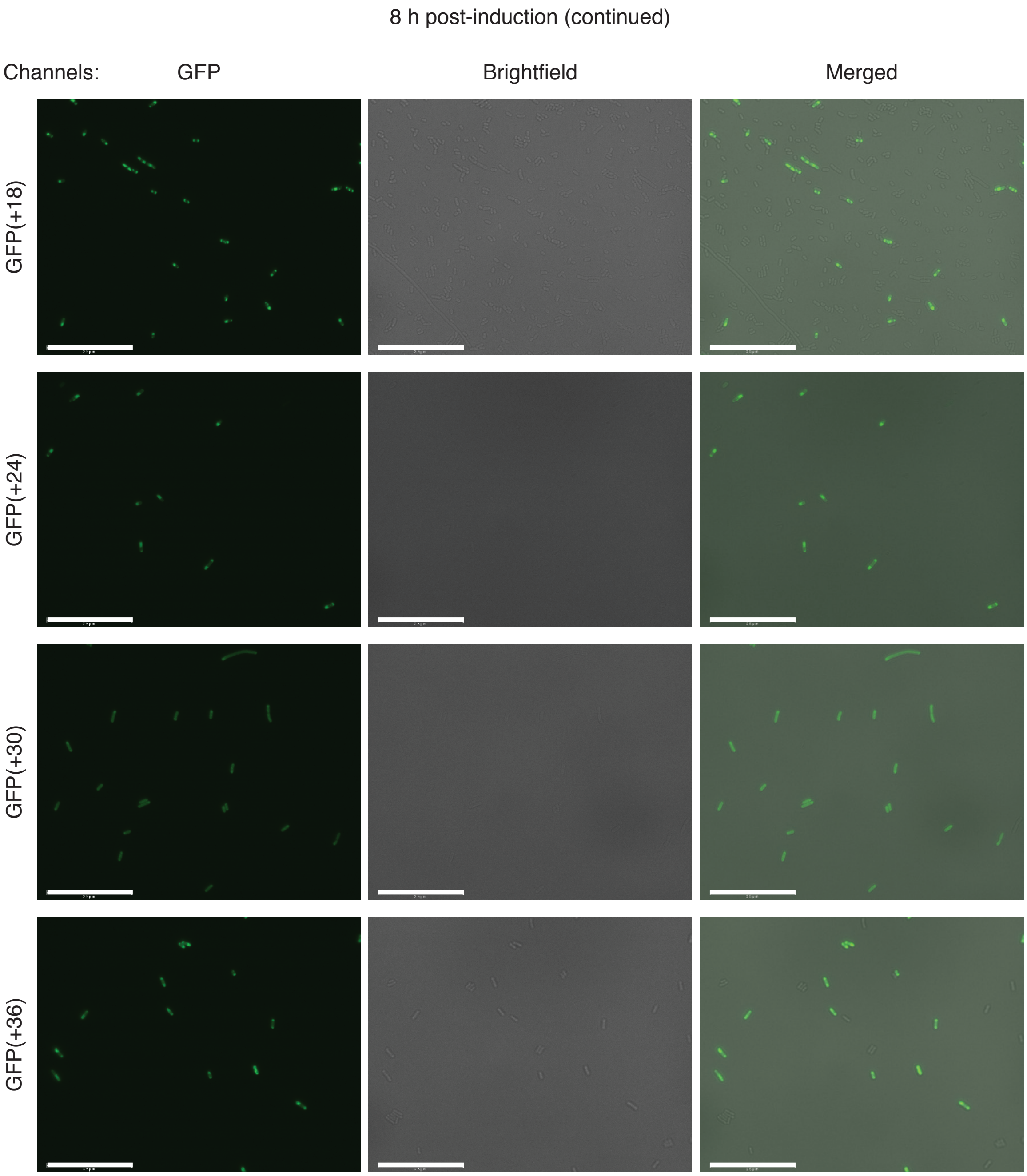


**Figure S14.**

**Representative images acquired from microscopy of cells expressing each supercharged variant at 8 h post-induction.** Representative cell populations are shown on GFP, brightfield, and merged channels. Scale bars are 25 μm.

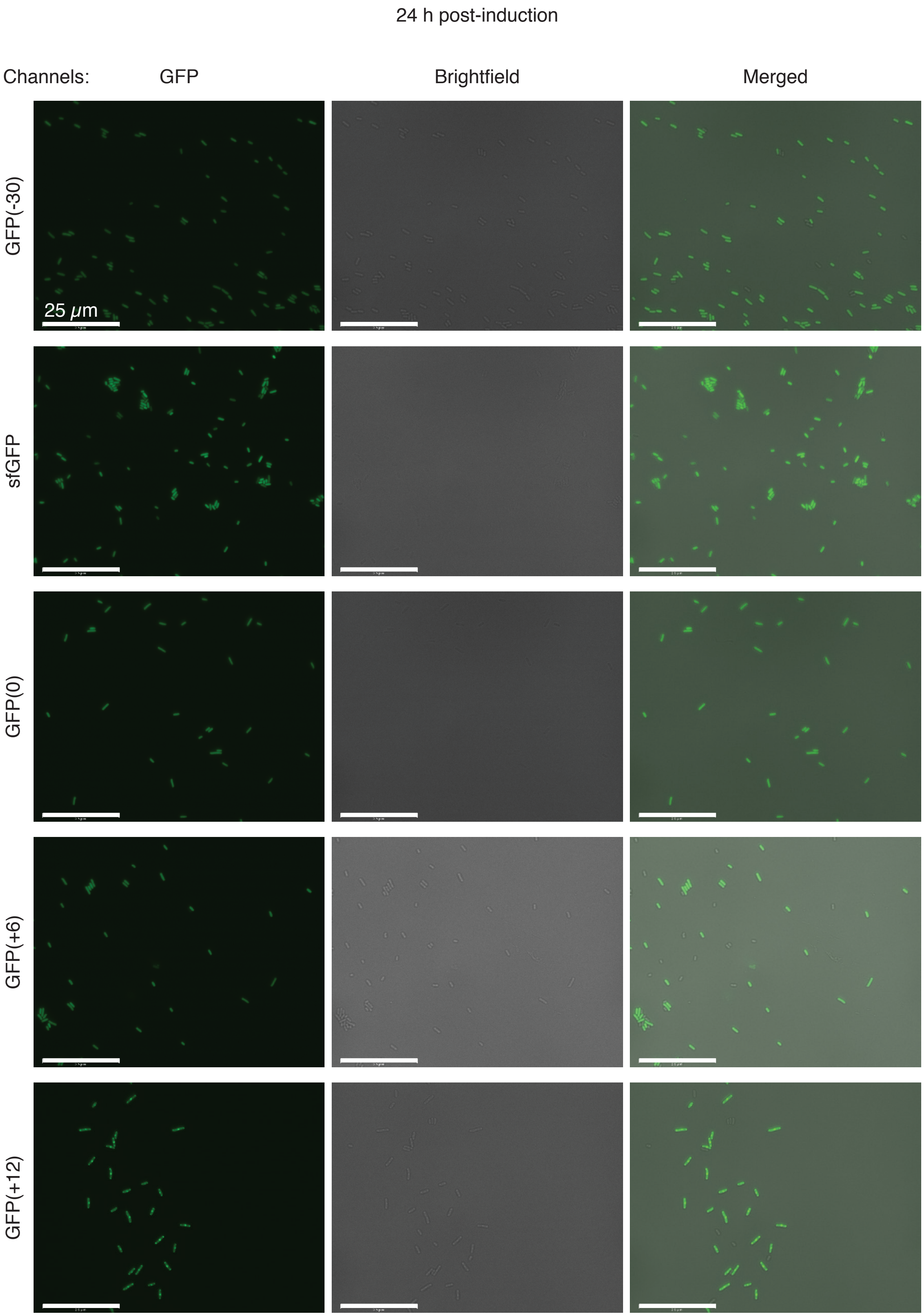

24 h post-induction
Channels:
GFP
Brightfield
Merged
GFP(-30)
sfGFP
GFP(0)
GFP(+6)
GFP(+12)
25 μm

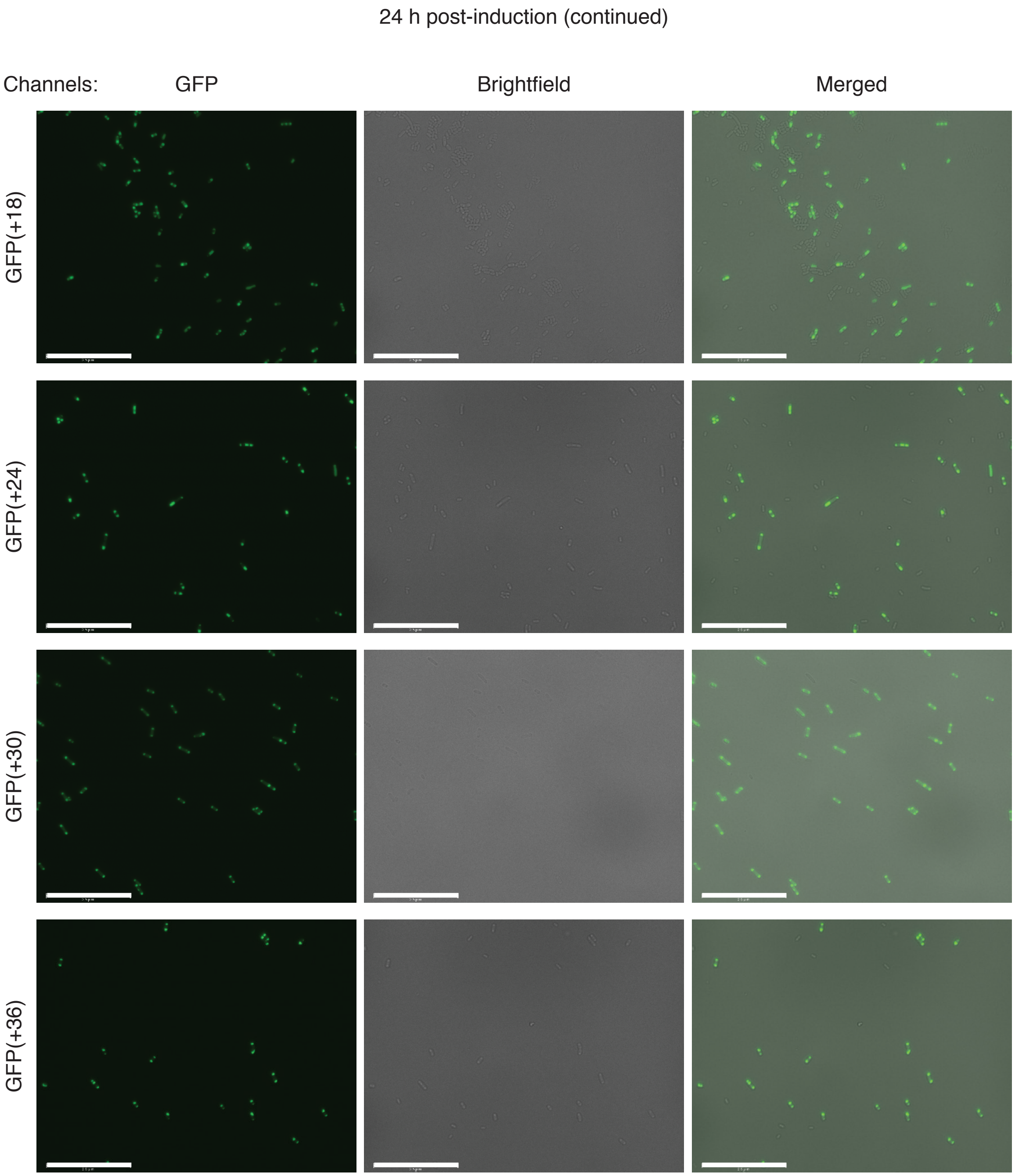


**Figure S15.**

**Representative images acquired from microscopy of cells expressing each supercharged variant at 24 h post-induction.** Representative cell populations are shown on GFP, brightfield, and merged channels. Scale bars are 25 μm.

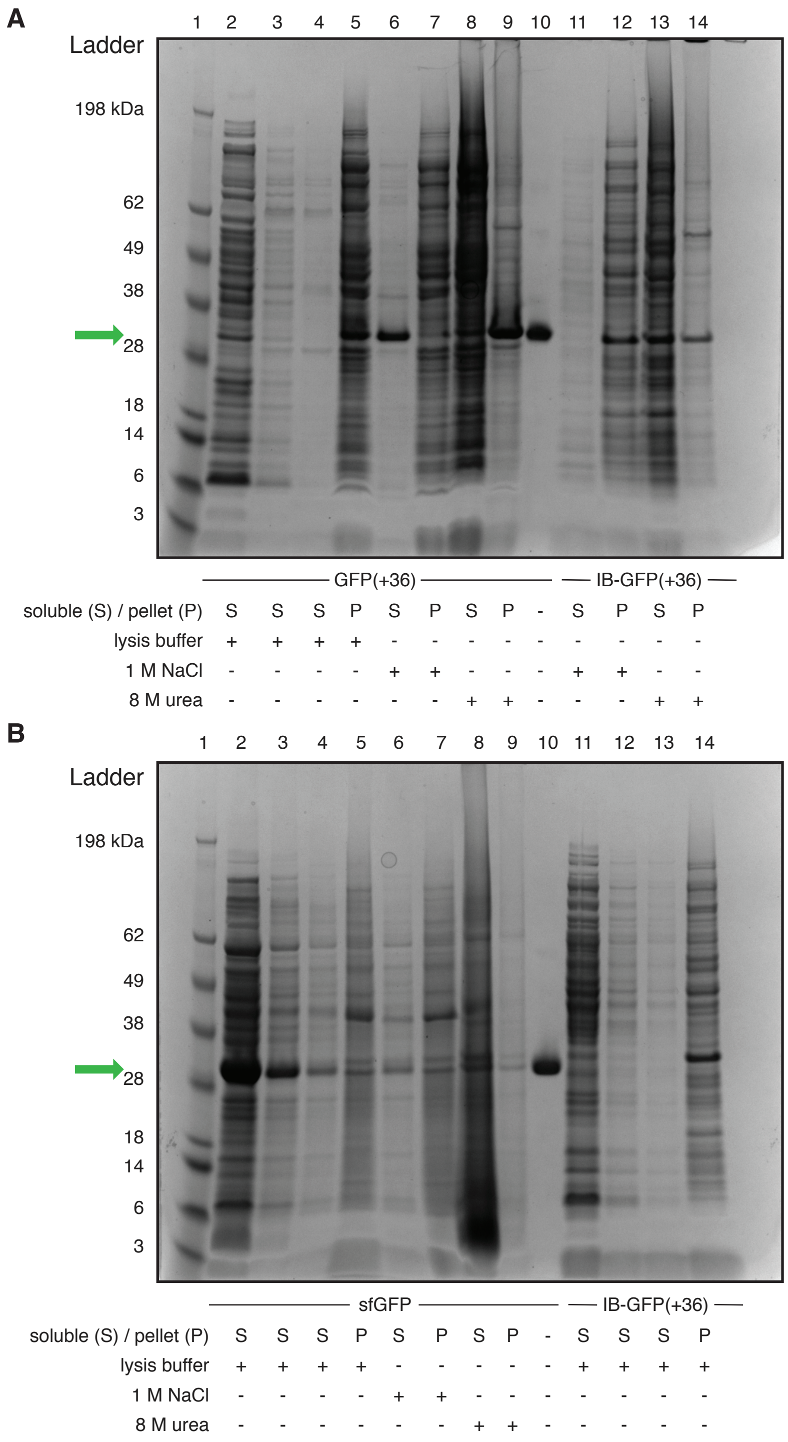


**Figure S16.**

**Solubility of soluble sfGFP, GFP coacervates, and GFP inclusion bodies.** SDS-PAGE gels depicting differences in solubility between coacervate-based condensates and inclusion bodies. "+" indicates treatment with the specified buffer (lysis buffer, 1 M NaCl or 8 M urea) and "S" and "P" refer to the soluble fraction and pellet, respectively. For both gels, lane 1 corresponds to SeeBlue pre-stained protein standard (Fisher Scientific) and lane 10 corresponds to purified protein. The band corresponding to GFP is indicated by the green arrow. (A) Lanes 2 and 3 depict the pre- and first washes of the GFP(+36) lysates, respectively. Lanes 4-9 depict different buffer treatments of GFP(+36) lysates. (B) sfGFP samples were loaded in the same order as GFP(+36) samples. Lanes 11-14 on both gels (A and B) show IB-GFP(+36) samples. Pre- and first washes of IB-GFP(+36) lysates were loaded in lanes 11 and 12 in (B). All other lanes depict treatments of IB-GFP(+36) lysates.

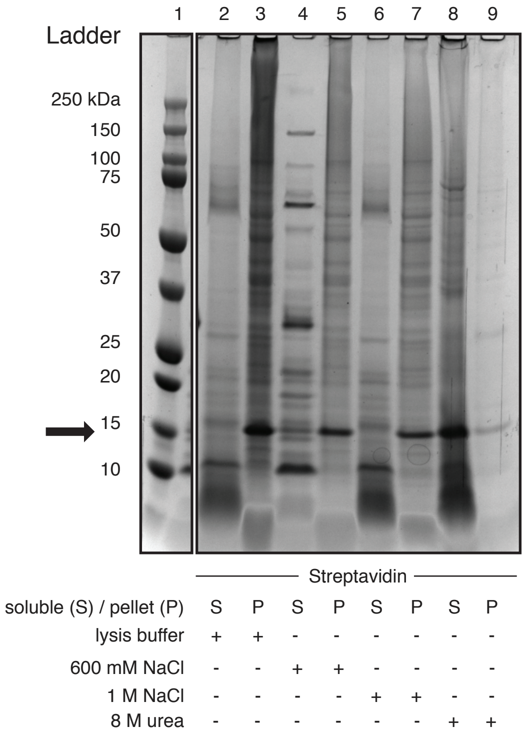


**Figure S17.**

**Solubility of streptavidin inclusion bodies.** SDS-PAGE was used to test the solubility of streptavidin, a recombinant protein known to form inclusion bodies in bacteria (*24*), in the presence of different buffers. Note that the precision plus dual color protein standard (Bio-rad) in lane 1 was loaded on the same gel as the subsequent samples but several irrelevant lanes were excluded for clarity. Various treatments of stretavidin lysates were loaded in lanes 2-9 as indicated below the gel image. "+" indicates treatment with the specified buffer and "S" and "P" refer to the soluble fraction and pellet, respectively. The band corresponding to streptavidin is indicated by the black arrow.

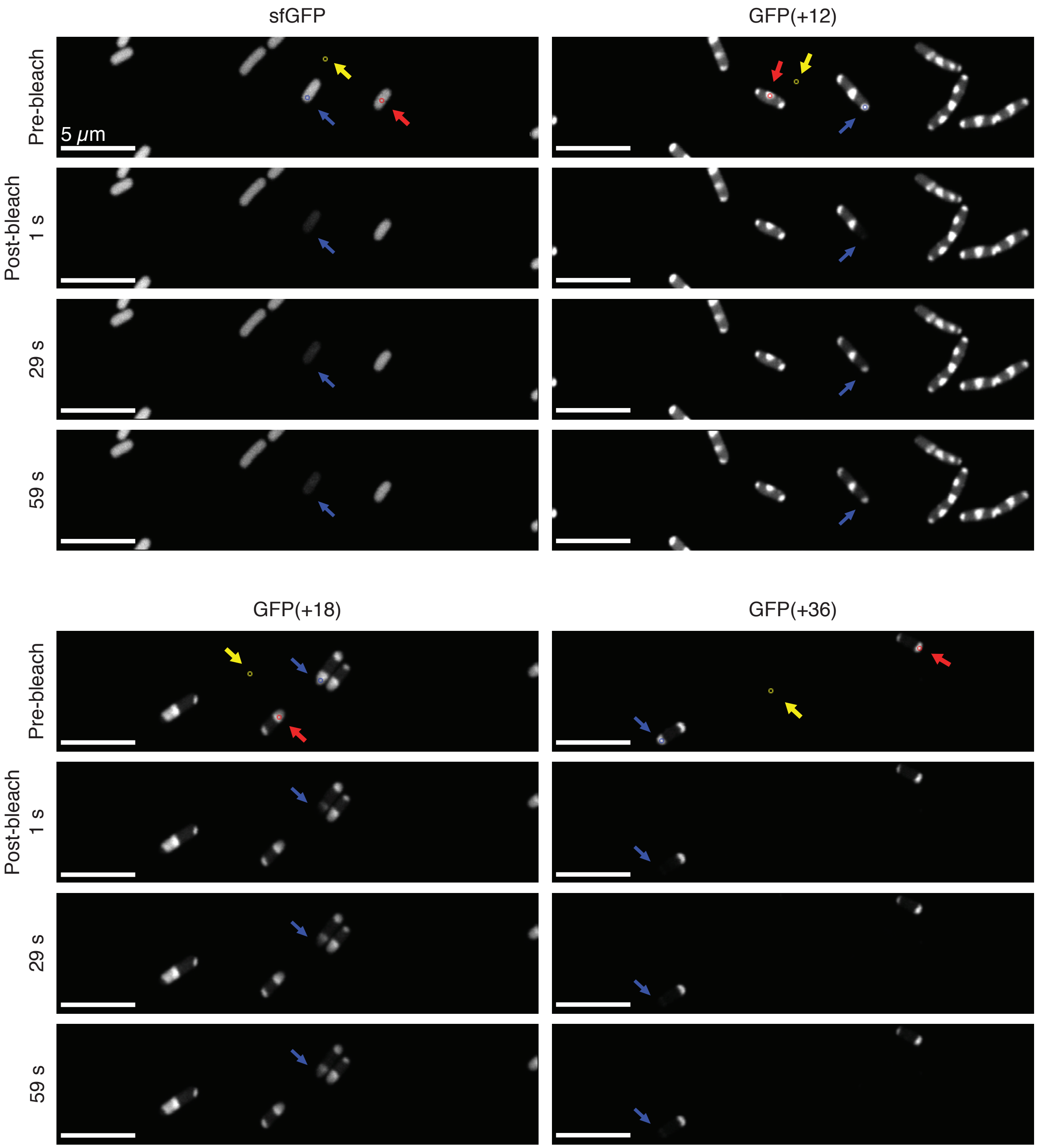


**Figure S18.**

**Representative images from FRAP experiments with condensates.** The bleached region is indicated by the blue circle and blue arrow guide, the background ROI is shown in yellow, and an unbleached cell in the same frame was chosen as a photobleaching control (shown in red). Scale bars are 5 μm.

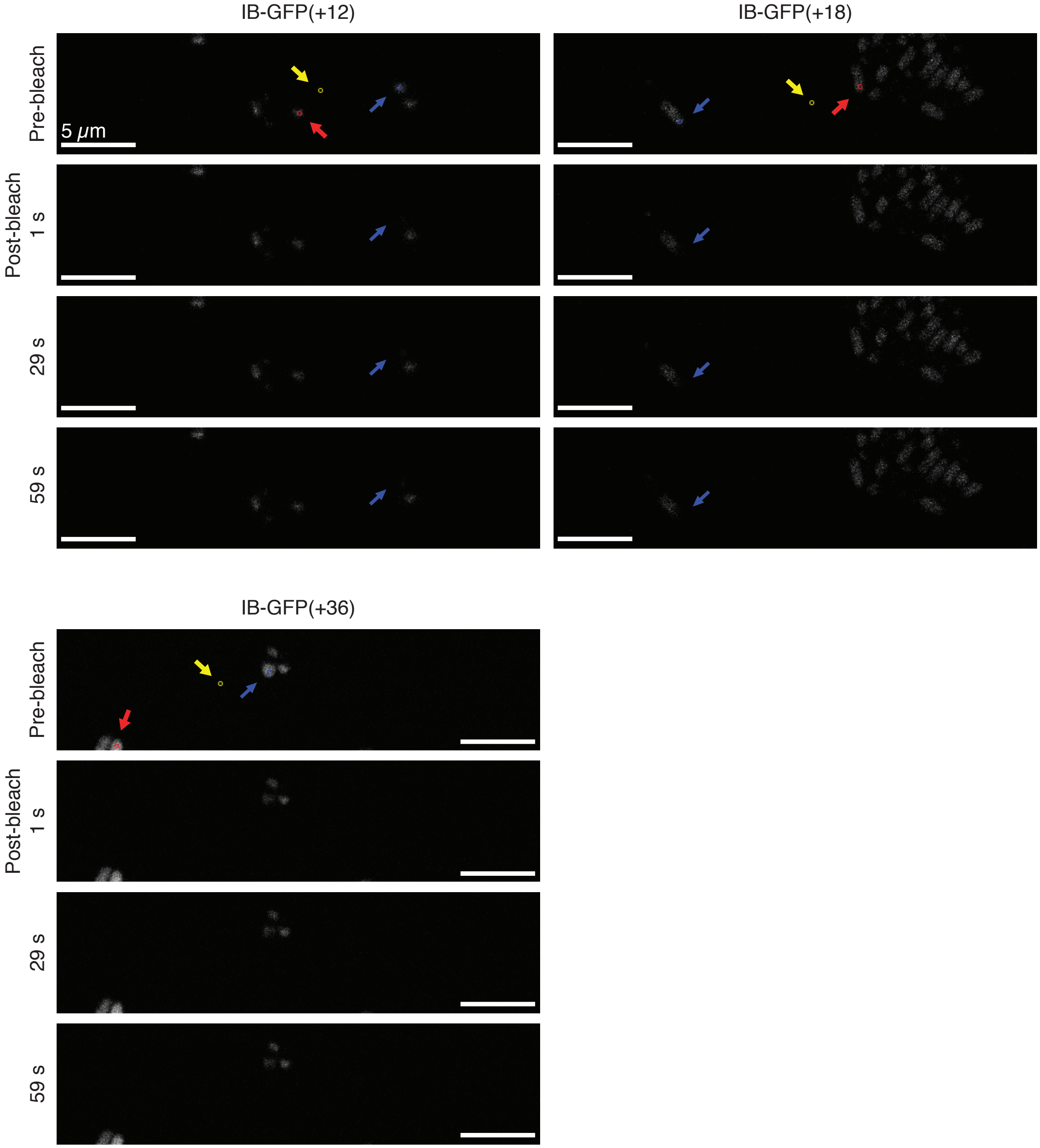


**Figure S19.**

**Representative images from FRAP experiments with inclusion bodies.** The bleached region is indicated by the blue circle and blue arrow guide, the background ROI is shown in yellow, and an unbleached cell in the same frame was chosen as a photobleaching control (shown in red). The overall fluorescence intensity of cells expressing each IB-GFP was significantly lower than their non-IB forming counterparts (Figure S18). Scale bars are 5 μm.

**A**

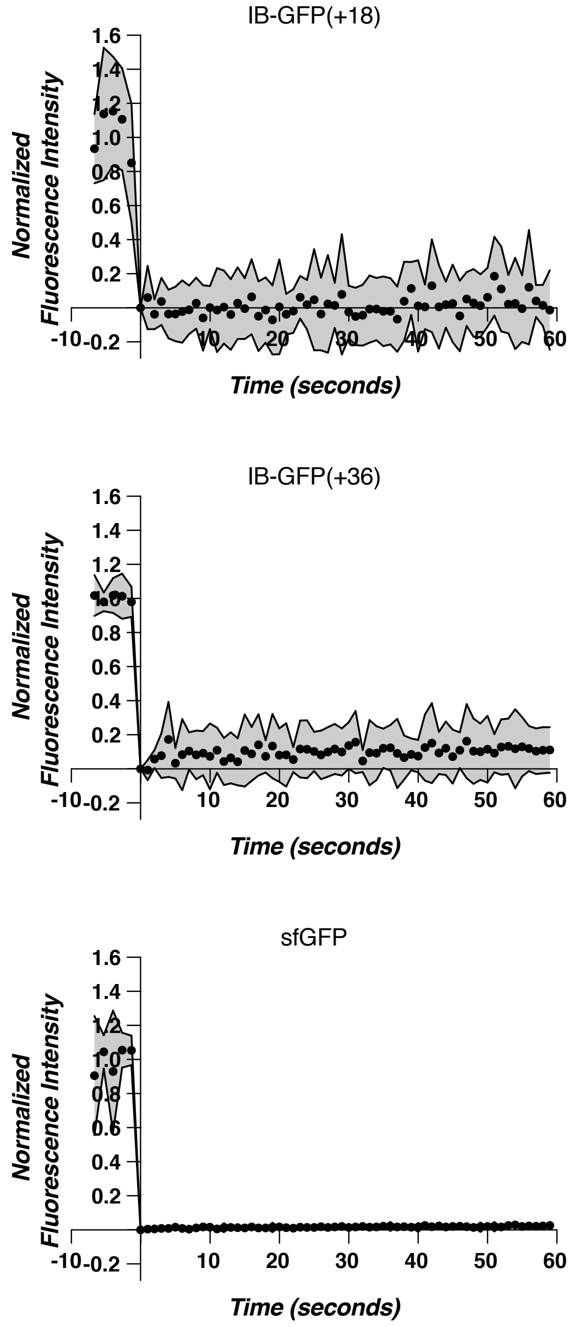
IB-GFP(+18)
Normalized Fluorescence Intensity
Time (seconds)
IB-GFP(+36)
Normalized Fluorescence Intensity
Time (seconds)
sfGFP
Normalized Fluorescence Intensity
Time (seconds)

B

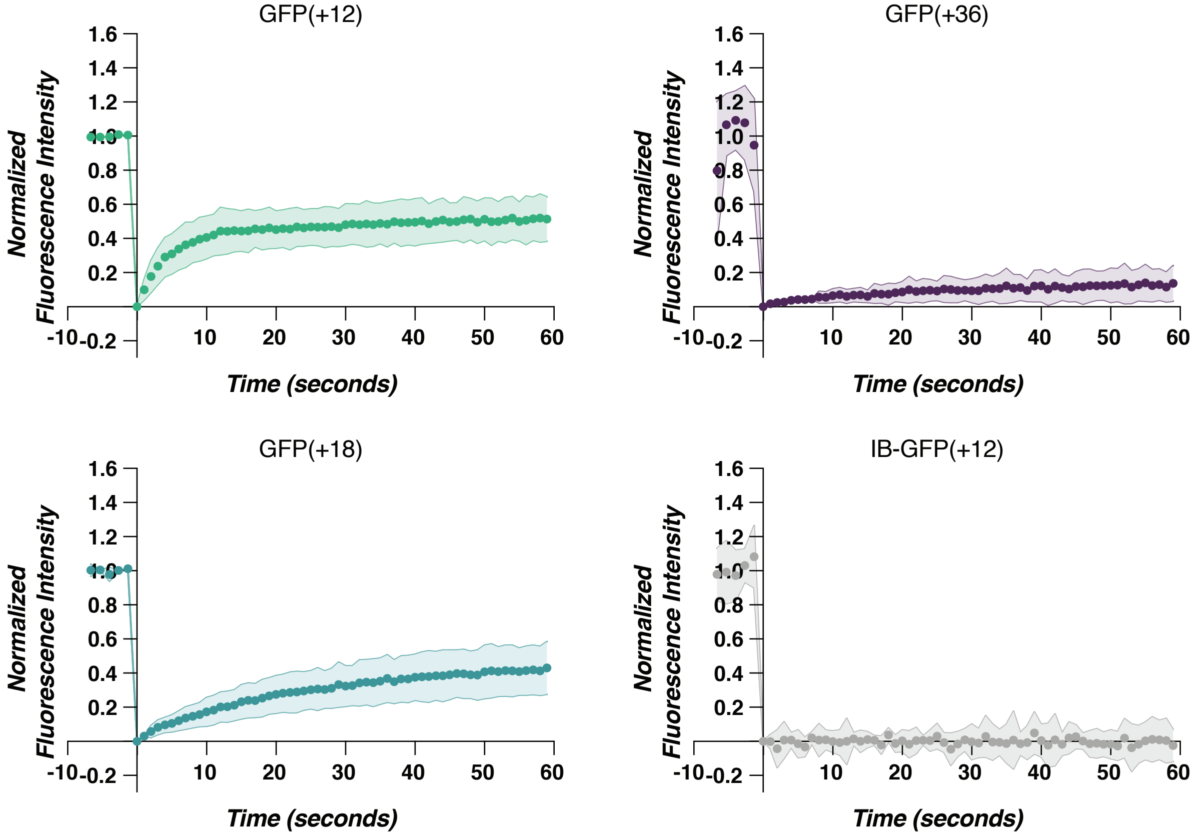


C

| GFP variant | # cells (n) | Mobile fraction | $t_{1/2}$ (sec) | $R^2$ |
|---|---|---|---|---|
| sfGFP | 9 | 0.02 | --- | 0.67 |
| GFP(+12) | 7 | 0.49 | 3.96 | 0.98 |
| GFP(+18) | 8 | 0.45 | 16.3 | 1 |
| GFP(+36) | 9 | 0.14 | 16.07 | 0.96 |
| IB-GFP(+12) | 5 | 0 | --- | 0 |
| IB-GFP(+18) | 7 | 0.01 | --- | 0.03 |
| IB-GFP(+36) | 8 | 0.1 | --- | 0.28 |

**Figure S20.**

**FRAP analysis of (A) inclusion body dynamics and (B) individual replicates.** (A) Inclusion bodies do not exhibit fluorescence recovery when bleached because they do not engage in material exchange with the surrounding cytoplasm. Depicted are the averages of normalized fluorescence intensities (dots), and the standard deviation (gray). The FRAP settings used were not optimized for the fast diffusion of sfGFP in the cytoplasm, which explains why fluorescence recovery is not observed as all of the GFP in the cell was bleached during the bleaching event. (B) FRAP curves of individual GFP variants depicted in Figure 2B. (C) Summary table of all GFP variants, sample sizes, mobile fractions, half-lives, and R-squared values obtained from single exponential fits. The recovery curves of sfGFP and IB-GFP variants did not fit to a single exponential function as indicated by their R-squared values. As a result, their half-lives were not reported.

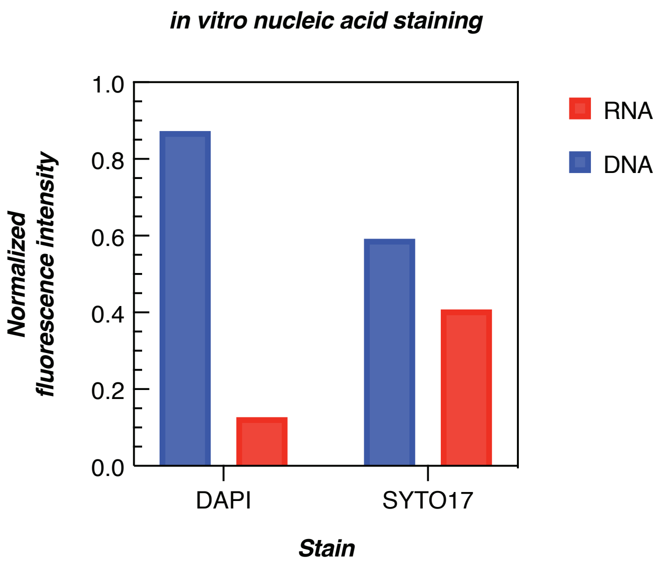


**Figure S21.**

***In vitro* nucleic acid staining.** DAPI and SYTO17 staining of DNA or RNA. *In vitro* staining of DNA or RNA with DAPI or SYTO17 demonstrates that DAPI exhibits a 7-fold fluorescence intensity increase in the presence of DNA (relative to RNA) while SYTO17 exhibits a 1.5-fold increase. Fluorescence intensities were normalized for each dye.

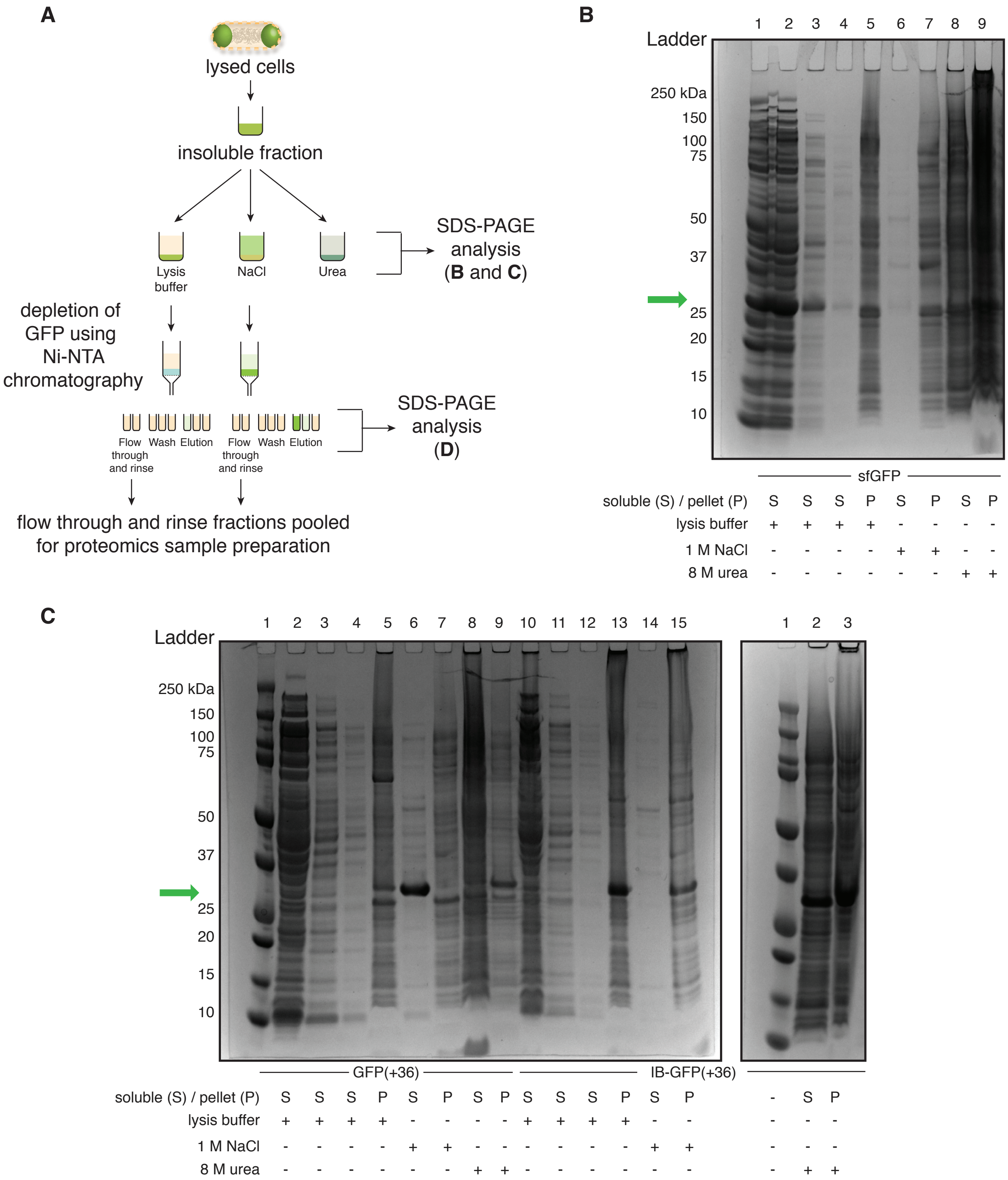
A
lysed cells
insoluble fraction
Lysis buffer
NaCl
Urea
SDS-PAGE analysis (B and C)
depletion of GFP using Ni-NTA chromatography
Flow through and rinse
Wash
Elution
SDS-PAGE analysis (D)
flow through and rinse fractions pooled for proteomics sample preparation
B
Ladder
1 2 3 4 5 6 7 8 9
250 kDa
150
100
75
50
37
25
20
15
10
sfGFP
soluble (S) / pellet (P) S S S P S P S P
lysis buffer + + + + - - - -
1 M NaCl - - - - + + - -
8 M urea - - - - - - + +
C
Ladder
1 2 3 4 5 6 7 8 9 10 11 12 13 14 15
1 2 3
250 kDa
150
100
75
50
37
25
20
15
10
GFP(+36)
IB-GFP(+36)
soluble (S) / pellet (P) S S S P S P S P S S S P S P - S P
lysis buffer + + + + - - - - + + + + - - - - -
1 M NaCl - - - - + + - - - - - - + + - - -
8 M urea - - - - - - + + - - - - - - - + +

D

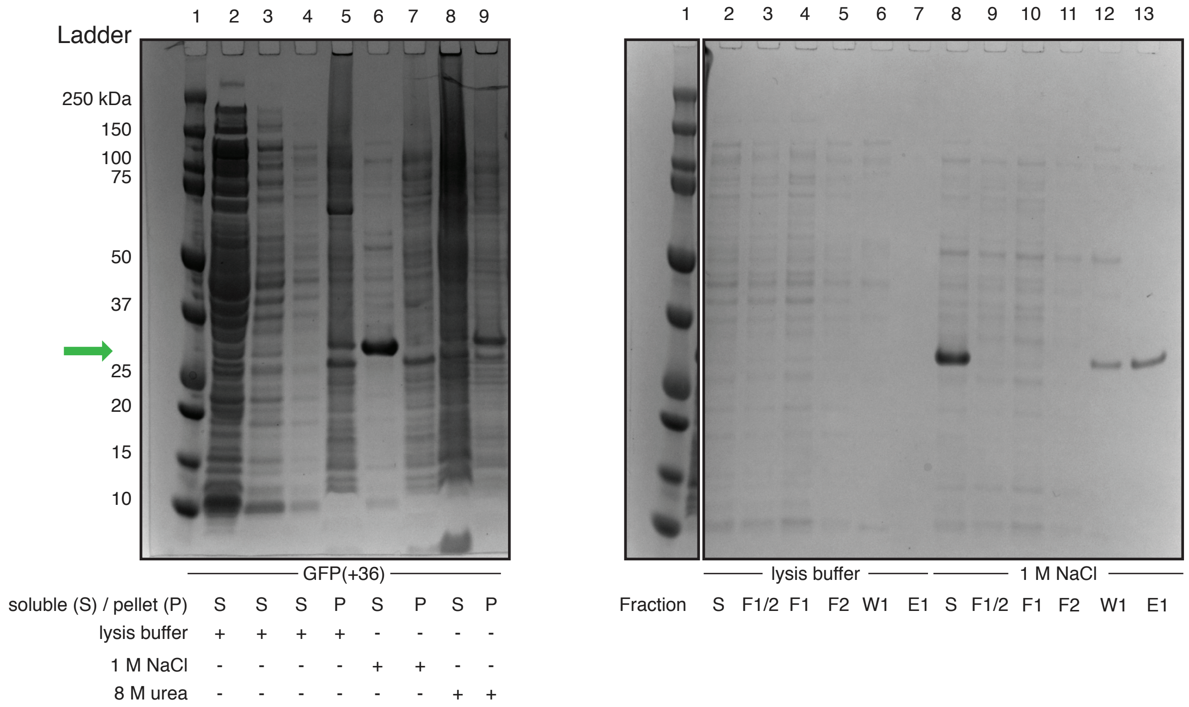


**Figure S22.**

**Representative SDS-PAGE analysis of samples submitted for quantitative proteomics.** Results here replicate findings from the solubility assay shown in Figure S16. (A) General schematic for isolation of the different fractions and removal of GFP from soluble fractions. Flow through rinse fractions were pooled and prepared for proteomics analysis. (B and C) SDS-PAGE gels showing the isolation of fractions. (D) SDS-PAGE gels depicting the Ni-NTA removal of GFP from the isolated soluble fractions. The band corresponding to GFP is indicated by the green arrow. (B) The pre- and first washes of sfGFP lysates are shown in lanes 2-3 and various treatments are depicted in lanes 4-9. Similarly, GFP(+36) lysates are depicted in lanes 2-9 (C, left), and IB-GFP(+36) lysates are depicted in lanes 10-15 (C, left) and 2-3 (C, right). Only soluble fractions from GFP(+36) treated with lysis buffer (lane 4) or NaCl (lane 6) were further processed for proteomics analysis. In all gels, lane 1 corresponds to precision plus protein dual color standard (Bio-rad) and “S” and “P” refer to the soluble fraction and pellet, respectively. (D) Isolation of GFP(+36) soluble fractions in different buffers (left); gel is reproduced from part (C). Fractions from Ni-NTA purification of soluble fractions obtained from resuspension in either lysis buffer or 1 M NaCl (right). Shown are the soluble fraction before Ni-NTA purification “S”, flow through fraction “F1”, rinse (with lysis buffer) fraction “F2”, first wash fraction “W1”, and first elution fraction “E1”. “F1/2” is the pooled F1 and F2 fraction. Irrelevant lanes were removed for visual clarity.

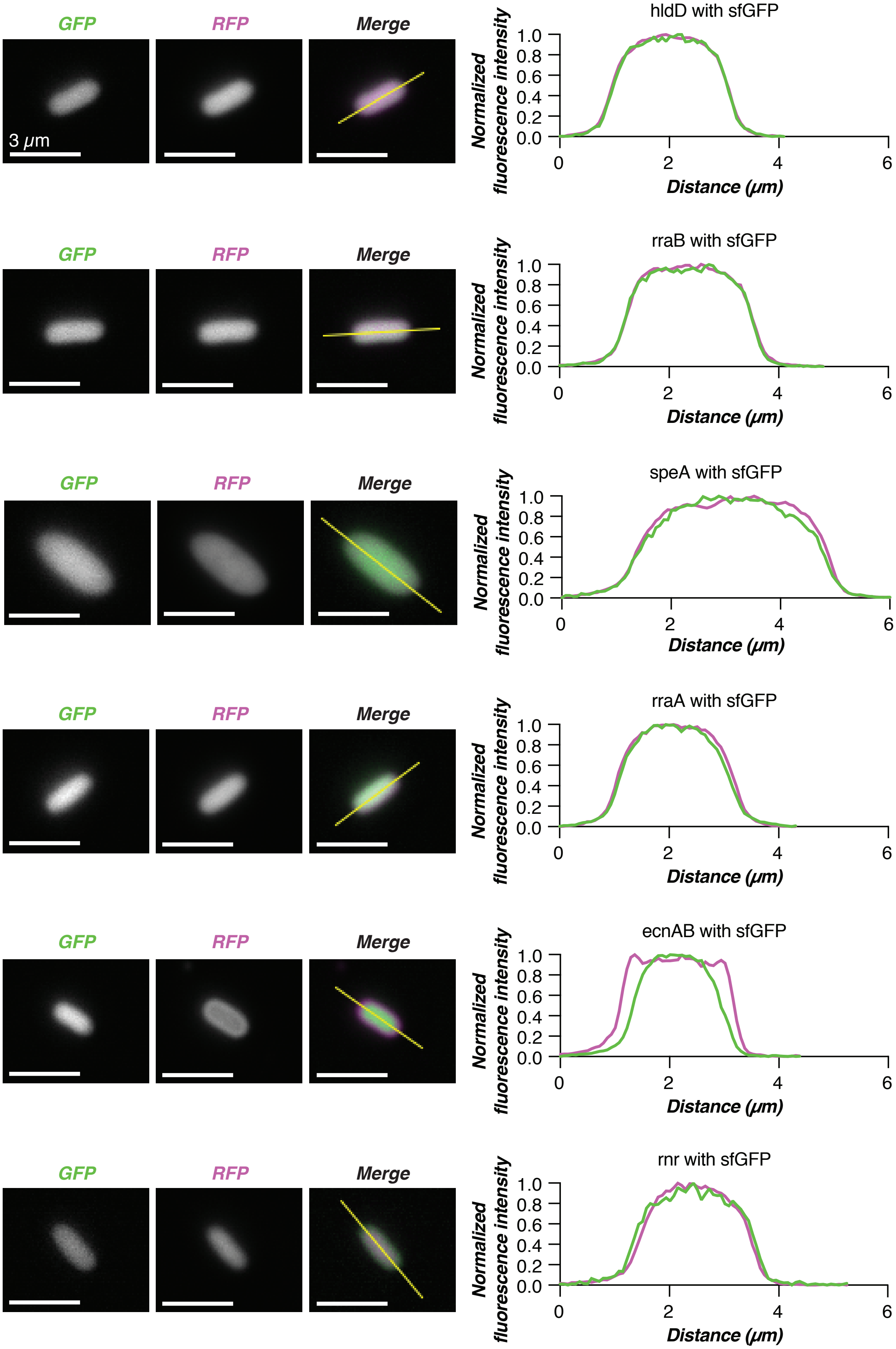
GFP
RFP
Merge
3 μm
hldD with sfGFP
rraB with sfGFP
speA with sfGFP
rraA with sfGFP
ecnAB with sfGFP
rnr with sfGFP
Normalized fluorescence intensity
Distance (μm)
1.0
0.8
0.6
0.4
0.2
0.0
0
2
4
6

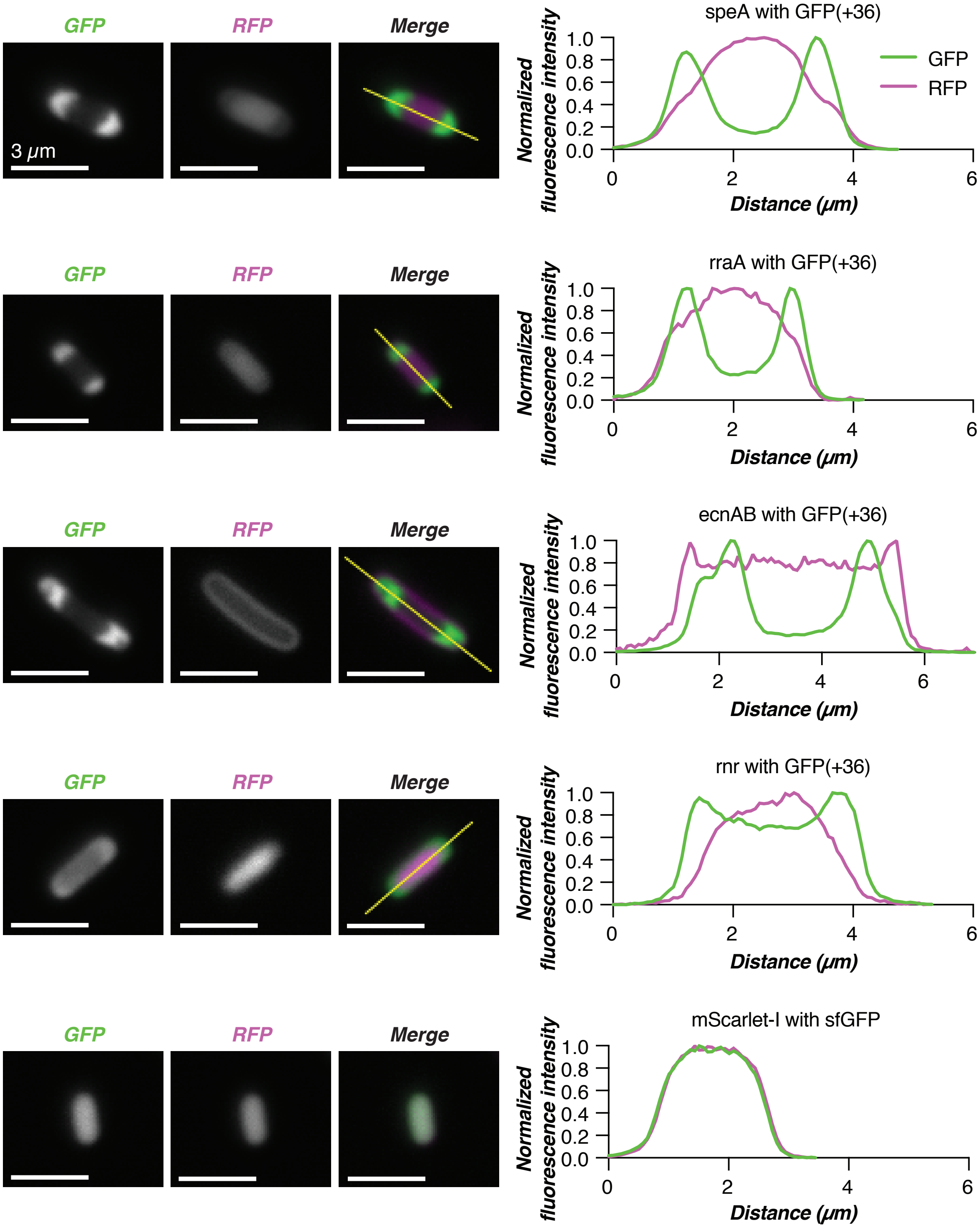


**Figure S23.**

**Additional proteomics validation candidates.** Co-expression of speA-, rraA-, ecnAB-, and rnr-mScarlet-I fusions with GFP(+36) or sfGFP. Additional sfGFP controls were conducted for mScarlet-I, hldD-mScarlet-I and rraB-mScarlet-I. Microscopy images (left) depict representative cells from co-expression with GFP and line-cuts demonstrate the extent of co-localization (right). Scale bars are 3 μm.

**Data S1. (separate file)**

**Protein enrichment in the condensate as determined by quantitative proteomics.** UniProt accession numbers, fold-changes, and p-values are provided for *E. coli* proteins identified. Data represents three biological replicates for each group.
